# SpaceExpress enables differential spatial transcriptomics with natural coordinate systems

**DOI:** 10.1101/2024.12.19.628720

**Authors:** Yeojin Kim, Abhishek Ojha, Alex Schrader, Juyeon Lee, Zijun Wu, Ian M Traniello, Saumya Jain, Gene E Robinson, Hee Sun Han, Sihai D Zhao, Saurabh Sinha

## Abstract

Spatial transcriptomics (ST) technologies have enabled new explorations into the spatial organization of tissues and their functional implications. However, one of the most fundamental analyses – comparative analysis of spatial gene expression across phenotypes – remains a formidable challenge. We introduce SpaceExpress, a novel statistical tool for detecting phenotype-associated changes in spatial expression patterns. SpaceExpress employs a neural network to embed multiple ST samples in a common latent space, enabling robust cross-sample comparisons despite structural and technical variations. It then uses spline regression to test differential spatial expression of genes between conditions, identifying specific regions of the tissue where expression patterns diverge and quantifying the magnitude of those differences, with rigorous false discovery control and handling of multiple replicates per condition. It includes visualization tools to help interpret spatial pattern differences. We demonstrate the tool’s effectiveness on synthetic and real ST datasets, revealing mechanistic insights into behavior- and development-related neurogenomic changes in honey bees and mice. Our work extends the highly influential paradigm of differential gene expression analysis to spatial omics.

## INTRODUCTION

The rapid development of highly multiplexed spatially resolved omics technologies, including spatial transcriptomics (ST)^1,2^, has opened new avenues of inquiry into the interplay of intercellular communication and intracellular gene regulatory mechanisms, as well as the spatio-temporal organization of tissues and its relationship to function. As ST technologies mature and costs decrease, research groups will collect ST measurements from many samples to understand how the fine global structure of spatial gene expression might be associated with phenotypes such as disease states and quantitative traits.

The field therefore needs tools for comparative ST analyses. These tools must be able to quantitatively describe associations between spatial gene expression and phenotypes in an interpretable way, as well as test for their statistical significance. This will be an important step in advancing from the largely descriptive nature of current ST analytics towards a more mechanistic understanding of how biological systems function. For non-spatial transcriptomics data, rigorous detection of differential^3^ or trait-associated^4^ gene expression has been central to mechanistic studies. In contrast, ST data call for a new comparative analysis, which we call “differential *spatial* expression” (DSE) analysis: to detect and describe alterations in the fine spatial distribution of gene expression across phenotypes. However, to the best of our knowledge, no such tools currently exist for spatial gene expression, posing a significant barrier to achieving the full potential of ST technologies. Most existing methods simply test for differential expression between spatial domains across ST samples^5^, which can only capture coarse forms of differential spatial patterning.

DSE analysis is a challenging problem for both technical and conceptual reasons. Technically, ST samples can vary for reasons that are not of direct interest, e.g., natural variation in tissue structure or differences in sectioning. Conceptually, it is difficult to even define a quantitative construct that can flexibly describe the innumerable ways in which the spatial structure of gene expression may associate with other traits. Available methods for describing spatial gene expression patterns in a tissue^6^ do not furnish a way to compare those patterns across tissues, especially in the face of structural variations noted above. Simply aligning multiple ST samples^7,8^ or mapping cells across samples^9,10^ does not provide a useful way to quantitatively describe differential patterns or rigorously test for associations with phenotypic traits, because the Cartesian coordinate system is not appropriate for tissues with complex geometric and topological structures. Furthermore, relying on automated inter-tissue alignments^11^] makes the comparisons vulnerable to the effects of misalignment.

We propose a novel approach to comparative analysis for spatial biology and use it to develop a statistical tool for inferring cross-sample or phenotype-associated changes in spatial expression patterns. Our new tool, called SpaceExpress, learns a new coordinate system that can most naturally describe locations within the tissue reflecting its intrinsic biological structure. This natural coordinate system is *shared* across analyzed samples, in the sense that biologically analogous regions of different samples are assigned similar coordinates. We demonstrate the accuracy of this common coordinate system through extensive evaluations on synthetic data and four different ST datasets obtained by three different technologies of varying resolution. Included visualization tools aid in interpreting the coordinate system. SpaceExpress directly compares spatial gene expression patterns between ST samples from different conditions along each axis of this coordinate system. It employs spline regression modeling to capture how expression along these natural axes differs across phenotypes and to formulate a rigorous hypothesis testing framework for differential spatial expression (DSE). The DSE concept introduced here is distinct from and complementary to the widely used differential expression^12^ (DE) strategy, which tests for changes in average gene expression levels between groups (**Figure 1a**). We apply SpaceExpress to compare three distinct spatial transcriptomic brain datasets (**Figure 1b**). First, we compare whole brain sections and specific brain substructures between a forager and soldier honey bee, using ST data that we generated, and identify statistically significant DSE genes that yield mechanistic hypotheses about behavior-related neurogenomic changes. Second, we compare visual cortex samples from normally reared and dark-reared mice, revealing spatial transcriptional patterns related to sensory experience. Third, we analyze two groups of ST samples from the mouse hypothalamic preoptic area, highlighting spatial expression differences associated with aggressive behavior. In summary, we present a novel approach to multi-sample ST data analysis capable of identifying spatial axes capturing the fine global structure shared by those samples, and rigorously testing the differential spatial expression of genes.

**Figure 1.**
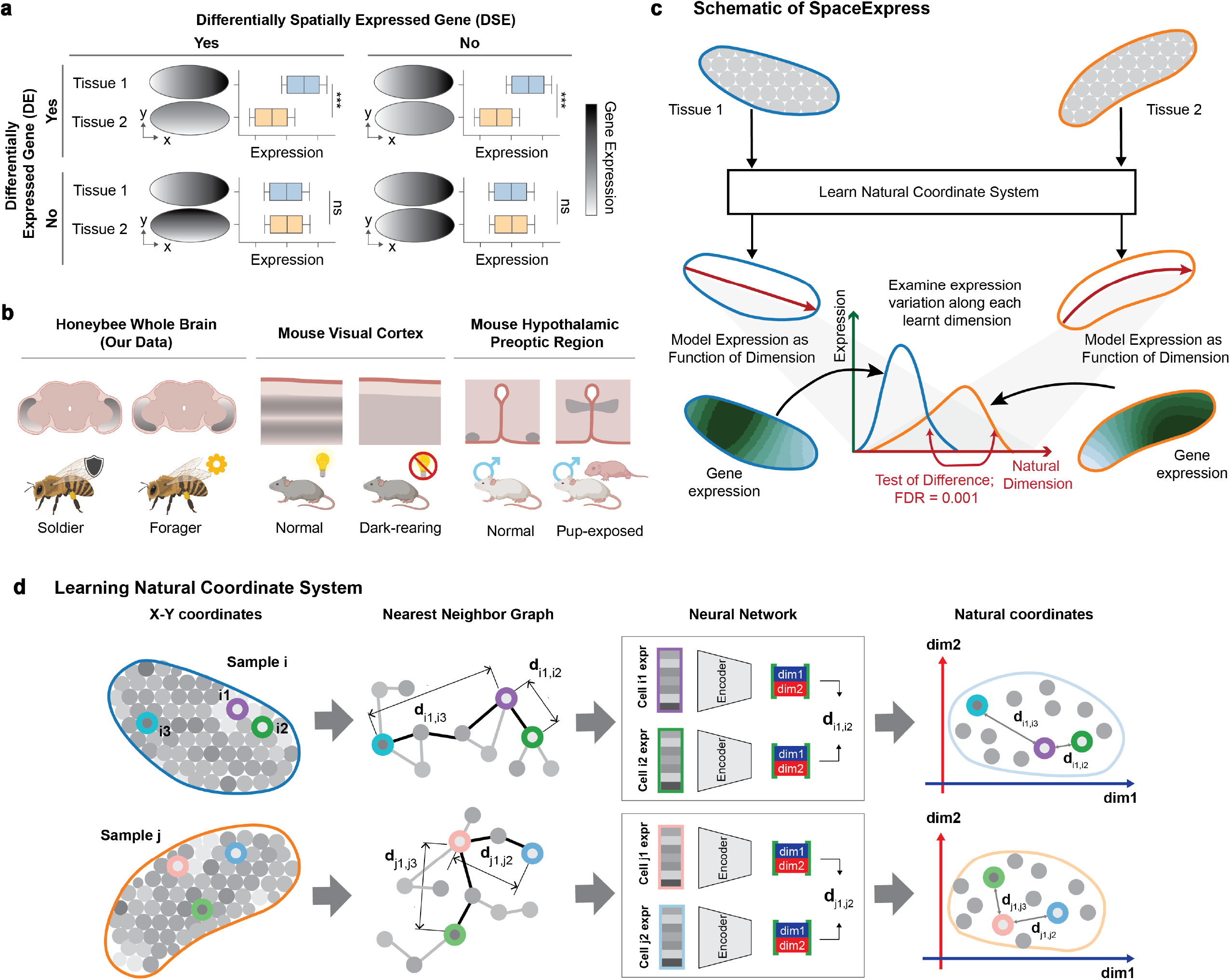
Overview of SpaceExpress Framework and Evaluations of SpaceExpress cell embeddings on synthetic data. **(a)** Schematic contrasting conventional differential expression (DE) analysis, which tests for changes in average gene expression levels between two samples, with differential spatial expression (DSE) analysis, which tests for alterations in the spatial distribution of gene expression across samples. In each scenario the left subpanel shows two idealized tissue regions colored by expression intensity and the right subpanel shows boxplots of gene expression for Tissue 1 (blue) and Tissue 2 (orange). Asterisks indicate significant differences and “ns” indicates no significant spatial change. **(b)** Application of SpaceExpress to three spatial transcriptomic brain datasets comparing two phenotypes. First, whole brain sections of honey bee workers in soldier and forager roles are compared using custom MERFISH data. Second, mouse visual cortex samples from normally reared and dark reared animals are compared. Third, samples from the mouse hypothalamic preoptic area are compared between naïve males and males exposed to pups. **(c)** SpaceExpress detects differential spatial expression between two tissue samples. In its first step (bottom), SpaceExpress maps (“embeds”) cells of each tissue sample in a common latent space based on their locations as well as gene expression profiles. Dimensions of the learnt latent space define a natural coordinate system shared by the samples, assigning similar coordinates to analogous tissue structures. In step 2 (top), each gene *G* is analyzed for differential spatial expression (DSE) between the tissue samples: spline regression is used to model the gene’s expression in a cell as a function of the cell’s coordinate along a particular dimension, and a statistical test is performed to assess whether this function differs between the samples, with the results quantified as a false discovery rate (FDR). (This is repeated for each dimension.) The same approach also applies to detecting a gene’s DSE between two groups of samples. **(d)** Schematic of step 1 of SpaceExpress. Spatial information in given tissue sample(s) is represented by a nearest neighbor graph of cells, with distances between cells (e.g., *d*_*ij*_ between cells *i* and *j*) defined as shortest path lengths between them and reflecting their physical proximity in the tissue. Expression profile of each cell (*g*_*i*_) is mapped by an encoder (neural network) to a low-dimensional embedding, so that Euclidean distances between cells in the embedding space (e.g., *p*_*i*_ − *p*_*j*_ ∨) approximate their spatial distances (e.g., *d*_*ij*_). For simplicity, three dimensions are shown in the embedding space.

## RESULTS

### Overview of SpaceExpress

SpaceExpress is the first approach that can rigorously test whether and describe how a gene’s spatial expression pattern differs between samples. More generally, it can test if spatial differences in gene expression are associated with a trait across multiple spatial samples. Furthermore, it can characterize these associations in a way that aligns with the tissue’s inherent biological organization. It is challenging to compare expression patterns along conventional x-, y- and z-axes, since the correspondence between standard Cartesian coordinates and biologically relevant spatial features – such as developmental axes or tissue layers – varies across ST samples. SpaceExpress overcomes this fundamental limitation by learning a more natural coordinate system for the tissue that enables more meaningful comparisons of spatial gene expression patterns.

SpaceExpress first constructs a coordinate-free representation of the spatial locations of each cell in each sample using nearest neighbor graphs. It then learns a *d*-dimensional vector representation of each cell such that a pair of cells close (resp. far) in a sample’s graph have similar (resp. different) representations (**Figure 1c,d**, Methods). (The dimensionality *d* is user-configurable and typically set to a small value.) This step involves training a neural network to obtain a cell’s *d*-dimensional representation as a function of its gene expression profile. This is motivated by the insight that gene expression encodes spatial location, arising from the literature on “positional information”^13^. The learnt dimensions constitute a natural coordinate system (set of axes) that captures the fine global structure of the tissue as encoded in its transcriptome. This reflects a key insight behind our approach: since gene expression, when properly normalized, is comparable across samples, the learnt axes, produced by the same neural network for every tissue sample, are automatically comparable as well. These axes are invariant to technical variation such as rotation or scaling, adapt to complex tissue geometries, and can be easily interpreted by visualizing the learnt coordinates of each cell in a tissue sample.

SpaceExpress uses its axes to characterize whether and how a gene is differentially spatially expressed with respect to discrete or quantitative traits. For this, it projects all cells onto each dimension of the shared coordinate system and models a gene’s expression pattern as a mathematical function of the dimension. A gene is defined as differentially spatially expressed (DSE) with respect to a trait if this function varies across samples with different trait values (**Figure 1c**, Methods). Such an association is assessed in a statistical hypothesis testing framework. Differences in mean expression (DE^12^) between samples are already accounted for in the model design, ensuring that they do not confound the identification of DSE. It is possible for a gene to be DSE for some dimensions but not others. This is a major advantage of SpaceExpress: it can identify which intrinsic tissue structures are involved in a gene’s DSE.

### SpaceExpress learns natural tissue coordinate system across multiple samples

The *d-*dimensional vector representations learnt by SpaceExpress describe cellular locations across multiple spatial tissue samples in a common coordinate system, assigning analogous cells in different samples to similar coordinates, e.g., functional domains, specific positions on a biological axis, etc., are assigned similar coordinates. Our first tests of this crucial capability involved synthetic data sets. We synthesized an “early Drosophila embryo slice” with cells laid out uniformly in an elliptical area, with a subset of genes varying in expression along the “anterior-posterior” (“AP”) axis and/or “dorso-ventral” (“DV”) axis (**Figure 2a, Supplementary Table S1**); these are the natural axes of this simple model tissue. We generated two such synthetic embryos with identical cellular layouts, and also introduced inter-embryo differences in spatial patterns of a subset (20%) of genes (Figure 2a, Methods, **Supplementary Figure S1**), to mimic inter-sample biological differences expected in a comparative analysis scenario. The level of DSE of these genes was a variable parameter (*μ*).

**Figure 2.**
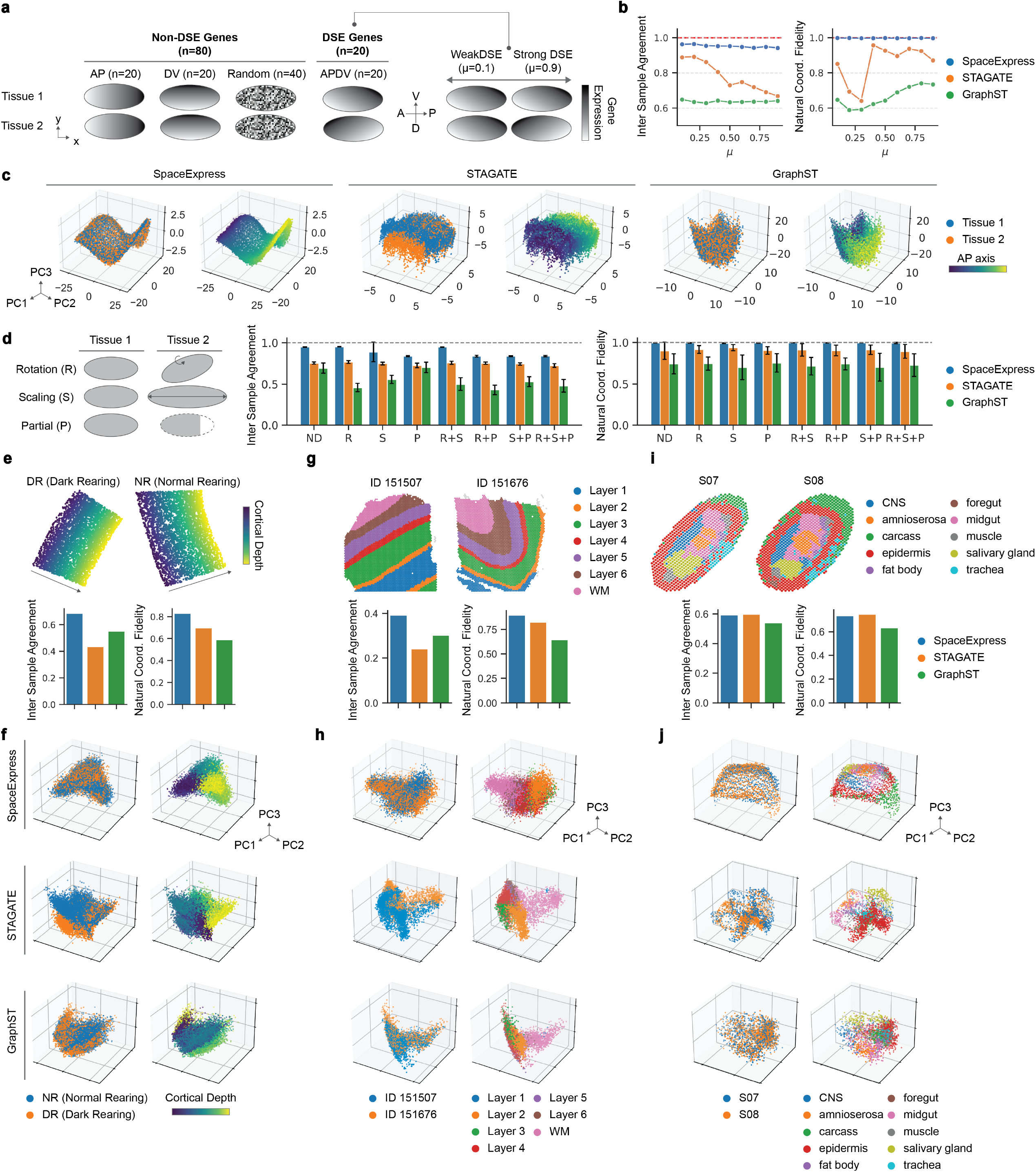
SpaceExpress learns a shared coordinate system for multi-sample comparative analysis. **(a)** Schematic of synthetic “early Drosophila embryo” data set, comprising two “tissue samples”, each with 3,915 cells arranged in an ellipse and gene expression patterns varying along the anterior–posterior (AP) and dorso–ventral (DV) axes; a subset (20%) of genes exhibits inter-sample differences and parameter *μ* controls DSE strength of these genes. **(b)** Inter-sample agreement (left, assesses if the learnt coordinate system is “shared”) and natural coordinate fidelity (right, evaluates if the learnt coordinate preserves the original AP axis) for SpaceExpress, STAGATE and GraphST across varying *μ*. Red dashed line indicates the maximum score of 1. PCA visualization of learnt coordinate system for data set at *μ* = 0.6. For each method evaluated, left panel shows cells colored by sample identity and right panel shows cells colored by ground-truth AP coordinate. **(d)** Evaluation of robustness to spatial distortions: rotation (30°), scaling (1.5×), partial truncation (20% removal) and their combinations. Bar plots show inter-sample agreement (left) and natural coordinate fidelity (right). Means across 10 replicates of synthetic data sets are shown, error bars indicate standard deviation. **(e-j)** Evaluations of shared coordinate system learnt for each of three real spatial transcriptomics datasets across diverse technologies and biological contexts: MERFISH profiles of normal-reared (NR) and dark-reared (DR) mouse visual cortex (e,f), Visium profiles of human DLPFC samples from two different individuals (g,h), and Stereo-seq profiles of adjacent slices of a Drosophila late-stage embryo (14-16h after egg laying) (i,j). Panels e,g,I (bottom) show agreement metrics – Inter-sample agreement (left) and natural coordinate fidelity (right) – for each method. Agreement score captures the extent to which learnt coordinate system preserves biologically meaningful correspondences between samples – based on cortical depth in the mouse visual cortex, and spatial domains in the DLPFC and Drosophila datasets. Panels f,h,j show PCA visualization of learnt coordinate systems. In each case, plot on left shows cells colored by sample identity and plot on right shows cells colored by ground-truth cortical depth / region annotation. In all cases, SpaceExpress achieves the highest inter-sample agreement and natural coordinate fidelity, demonstrating strong generalizability across platforms and tissue types.

We trained SpaceExpress on the two embryos jointly to obtain a 4-dimensional coordinate system and evaluated it using two complementary metrics: “inter-sample agreement” and “natural coordinate fidelity”. Inter-sample agreement assesses whether corresponding cells from different samples (as per the “ground truth”) are proximal in the learned coordinate system, i.e., it quantifies the “shared” nature of the coordinate system (see Methods). Natural coordinate fidelity, by contrast, measures how well spatial relationships within each sample are preserved in the learned coordinate system, i.e., it assesses if the coordinate system truly reflects spatial organization (see Methods). To benchmark performance, we compared SpaceExpress with two prominent ST tools – STAGATE^14^ and GraphST^15^. While neither method is designed to learn a natural coordinate system, these were relevant comparators for this evaluation as they support multi-sample learning of cellular representations for domain identification, by combining transcriptomic profiles of cells with their spatial location.

Our benchmarking shows that as DSE strength (*μ*) in the synthetic data is increased, making the task harder, SpaceExpress consistently maintains higher inter-sample agreement scores than both comparators, with STAGATE showing a marked drop at higher *μ* settings (**Figure 2b**, left). SpaceExpress also achieves the highest natural coordinate fidelity across all values of *μ*, as indicated by scores near 1 (**Figure 2b**, right). In contrast, STAGATE and GraphST show lower and more variable fidelity depending on *μ*, indicating that only SpaceExpress robustly captures the true underlying spatial axis even under divergent expression patterns.

Visualization of the joint embedding using PCA further illustrates these findings (**Figure 2c**). The left panels for each method depict the overlap of two synthetic samples in the learned coordinate space, for the data synthesized with μ = 0.6. SpaceExpress achieves near-complete overlap between samples, reflecting strong inter-sample agreement, while STAGATE and GraphST embed the two samples to overlapping but somewhat distinct regions in the shared space. The right panels show cells colored by their original anterior-posterior (AP) coordinate, allowing visual assessment of spatial fidelity. Only SpaceExpress recovered a continuous AP gradient, supporting that its learned coordinate system preserves biologically meaningful structure within each sample. (Similar visualizations of the DV axis are shown in **Supplementary Figure S2**.) We further tested robustness to spatial distortion, including rotation (30°), scaling (1.5x), and partial truncation (removal of 20% of one side of the tissue), as well as combinations thereof (**Figure 2d**). For these experiments, we also introduced variation in spatial expression patterns using *μ* = 0.5. Across all seven perturbation settings, SpaceExpress retained the highest inter-sample agreement and robust natural coordinate fidelity. In contrast, both STAGATE and GraphST were more sensitive to distortion, especially under combinations involving rotation and partial truncation.

To validate performance on real data, we applied joint training to three diverse ST datasets spanning different biological contexts and experimental platforms: (i) MERFISH profiles of visual cortex from dark-reared (DR) and normal-reared (NR) mice^16^ (**Figures 2e,f**), representing distinct experimental conditions; (ii) human dorso-lateral prefrontal cortex (DLPFC) samples from two individuals profiled using Visium^17^ (**Figures 2g,h**), representing biological replicates under the same condition; and (iii) Stereo-seq data from two adjacent slices of a late-stage Drosophila embryo^18^ (**Figures 2i,j**), representing technical replicates from the same specimen. In each case, SpaceExpress achieved the highest inter-sample agreement and natural coordinate fidelity across methods.

These results demonstrate that SpaceExpress can reliably learn a shared natural coordinate system for multiple ST samples, which is key to our goal of identifying DSE genes. The method is robust to both signal heterogeneity and spatial distortions, and broadly applicable to comparative analysis across biological conditions, replicates, or individuals. The learned coordinate system captures salient aspects of tissue organization, whether it is in the form of axes (synthetic embryos, visual cortex) or discrete regions (DLPFC, late embryo). *Note:* see **Supplementary Note 1** for additional evaluations of SpaceExpress-learnt coordinate systems, in the context of single-sample analysis, and **Supplementary Note 2** for further assessment of the model’s generalizability, showing that SpaceExpress trained on one sample can reconstruct spatial organization in unseen tissues based solely on gene expression.

### SpaceExpress identifies and characterizes differentially spatially expressed genes

The second step of SpaceExpress involves a rigorous statistical procedure for detecting differential spatial expression (DSE), achieved via nonparametric regression modeling and significance testing with false discovery rate (FDR) control (**Figure 3a-d, Supplementary Figure S3**, Methods). Once cells of all tissue samples have been embedded in a common *d*-dimensional space, each gene is tested for DSE along each of the dimensions. This involves fitting a spline regression model of a gene’s expression in a cell and using likelihood ratio statistics to assess if the contribution of the cell’s coordinate value to the model is sample-specific (or in the case of more than two samples, group-specific). FDR control is achieved using an empirical null fitting procedure^19^. The first step of learning a shared coordinate system may be prone to being compromised by DSE genes, since such genes, by definition, violate the premise of gene expression encoding space in a “universal” manner. To address this, once DSE genes have been identified, the neural network for mapping cells to a coordinate space is retrained without using those genes, and the statistical testing procedure is repeated a second time. We justified the individual steps of our testing procedure through an ablation analysis (**Figure 3e)**.

**Figure 3.**
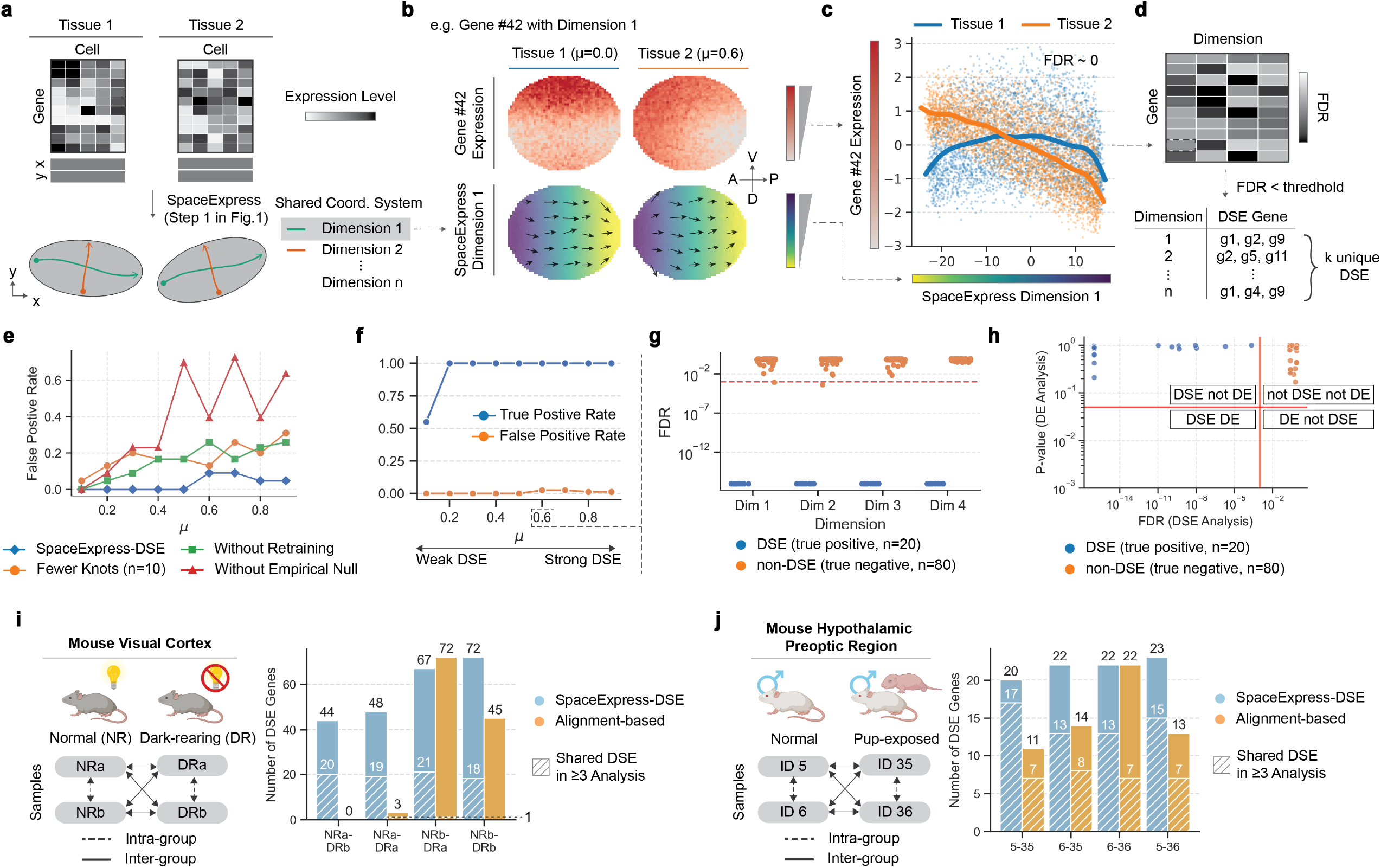
Statistical inference of differential spatial expression (DSE) by SpaceExpress. **(a-c)** Schematic overview of DSE testing, illustrated with the synthetic early embryo data set. Cells from two synthetic tissue samples are embedded in a shared coordinate system (a) by SpaceExpress, and each gene is tested for sample-specific variation along each dimension. Panels (b) show an example gene whose expression pattern differs between Tissue 1 and Tissue 2 (top). Part of this difference manifests in the gene’s expression variation along the AP axis, which is captured by SpaceExpress-learned dimension 1 (bottom). Panel c shows this difference in detail, with the gene’s expression in each cell (of either tissue) plotted against the cell’s coordinate value in dimension 1. The tissue-specific trends of this variation are visualized by the bold curves, which represent tissue-specific components of a spline regression model fit to all cells. The marked difference between these trends results in the gene being assigned an FDR ∼0 for dimension 1; all the gene-dimension combinations are similarly tested and assigned FDR values (panel d). Finally, all significant genes are reported for each dimension. **(e)** False positive rate at varying levels of signal strength (μ), showing performance of the SpaceExpress-DSE pipeline compared to ablated versions that lack the second retraining step, utilize fewer knots in spline model (n = 10 compared to n = 300), or lack the empirical null fitting step. **(f)** True positive rate and false positive rate for DSE detection across increasing values of *μ*, demonstrating robust sensitivity even to small signal strengths (*μ*). **(g)** FDR values assigned by SpaceExpress to DSE (n = 20) and non-DSE (n = 80) genes for each of four learned dimensions, indicating clear separation of significant hits by SpaceExpress. The red lines mark significance thresholds DSE analysis (1e−3). Results are shown for data set at μ = 0.6. **(h)** Comparison of DSE and traditional DE results, showing SpaceExpress-assigned FDR and Wilcoxon p-value respectively. (Results are for μ = 0.6.) The data set includes no DE genes, by design; consequently, all genes show nonsignificant p-values in DE analysis. In contrast, in DSE analysis using SpaceExpress with a specific dimension, all 20 DSE genes are correctly detected without false positives. The red lines mark significance thresholds for DE (p-value = 0.05) and DSE (FDR = 1e−3) analyses. **(i,j)** Evaluation of robustness of DSE identified by SpaceExpress in MERFISH profiles of mouse visual cortex under normal and dark-rearing conditions (i) and mouse hypothalamic preoptic region from naive and pup-exposed animals (j). In each case, four inter-group two-sample comparisons (bold arrows) and two intra-group comparisons (dashed arrows) were performed. DSE genes were detected (with statistical significance scores) for each comparison, using SpaceExpress or an alignment-based method. Bar plots show numbers of DSE genes detected (out of 500 genes tested) in the four inter-group comparisons, at a significance level stronger than any seen in the intra-group comparisons by the same method. Each bar also shows the number of genes designated DSE in three or all of the four inter-group comparisons; see Methods for full details.

We performed systematic tests of the sensitivity and precision of our approach using the synthetic data sets comprising pairs of “early embryos”, where 20 of 100 genes are made DSE in a controlled manner, as presented in the previous section (Figure 2a, Methods). Figures 3a-d illustrate how SpaceExpress identifies such a DSE gene. This exemplar gene exhibits a moderate level of DSE (Figure 3b, top), manifested along the AP as well as DV axis (**Supplementary Figure S4a,b**), and SpaceExpress-learnt dimensions capture these axes (Figures 3a,b, **Supplementary Figure S5**). Figure 3c shows the gene’s expression in individual cells plotted against the dimension 1 coordinates, and the fitted spline regression model reveals the distinct expression trends in the two embryos. This divergence is quantified in a high log likelihood ratio statistic, which is then assigned an FDR of approximately 0. We evaluated the performance of SpaceExpress across all 100 genes and across varying levels of DSE strength (*μ*), observing true positive rates ∼1 with false positive rates below 0.02 (**Figure 3f, Supplementary Data S1**) at most values of *μ*. For instance, at *μ* = 0.6, SpaceExpress shows a clear separation in FDR between the two classes across all dimensions (**Figure 3g**). Notably, sensitivity remains high even for subtle spatial changes (*μ* < 0.3), enabling detection of fine-grained DSE that may be visually indiscernible (Supplementary Figure S4c,d). In many cases the DSE was detected for the same gene in multiple dimensions (**Supplementary Data S2**). Additionally, SpaceExpress also identifies DSE genes that are missed by conventional differential expression (DE) analyses (**Figure 3h**), and has a superior accuracy than the alignment-based River method^11^ (Supplementary Figure 31a).

To test DSE detection on real data, we applied SpaceExpress to four pairs of tissue samples of mouse visual cortex under normal (NR) and dark-rearing (DR) conditions (**Figure 3i**, Methods). This identified DSE genes in these four NR–DR comparisons, and 21 genes were significantly DSE in three or more of the analyses. The significance threshold was set at a level not crossed by any gene when tested for DSE between intra-condition replicates (NR-NR and DR-DR comparisons), thus ensuring strong control over false positive. As a comparison point, we implemented an alignment-based approach to DSE detection that does not utilize a shared coordinate system, and repeated the above analysis to find only one gene to be consistently DSE (in ≥ 3 of pairwise comparisons) at equivalent levels of false positive errors. (Also see Supplementary Figure 31b for a comparison to the River method^11^ for ranking DSE genes.) A similar analysis of mouse hypothalamic preoptic area samples from naïve and pup-exposed mice yielded 18 robust DSE genes versus seven with the alignment-based comparator (**Figure 3j**, Methods). Thus, SpaceExpress achieves greater sensitivity in DSE detection with higher consistency from multiple trial replicates compared to alignment-based methods. These results provide objective evidence for the robustness of SpaceExpress in detecting DSE genes across different biological conditions.

### SpaceExpress pinpoints behavior-related changes in the honey bee brain spatial transcriptome

We illustrate the power of SpaceExpress by using it to identify DSE patterns associated with honey bee behavior. Honey bees form complex societies with specific castes responsible for different tasks and previous research has revealed that neurotranscriptomic state is tightly linked to a bee’s occupation in the hive^20-22^. However, these transcriptomic states^20,21,23,24^ have lacked spatial resolution and it remains unclear whether behavioral differences between castes such as soldiers and foragers are associated with regional variations in expression. Studying anatomical locations responsible for this DSE may reveal principles of how complex behaviors emerge from the spatial organization of cells in the bee brain.

To answer this question, we profiled the spatial expression of 130 behavior-related genes^23,24^ in similarly aged soldier and forager brains using our custom MERFISH platform and performed spatial comparisons using SpaceExpress. For the forager brain, we intentionally used a section that was only partially imaged (**Figure 4a**), to mimic the common challenges for differential spatial analysis, including size/morphology variation and partially damaged tissues. **Figure 4b** visualizes a four-dimensional natural coordinate system learned by SpaceExpress on the two brains. Despite dramatic visual differences between the samples, each learned dimension successfully captures anatomical structures previously characterized in honey bee brains, and assigns similar coordinates to the same regions across both brains. Additionally, different embedding dimensions distinguish different structures. For example, dimensions 2 and 4 clearly distinguish between mushroom bodies (MB), the primary region responsible for learning^25^, and the rest of the brain. Dimension 1 differentiates regions within MB, which correspond to different subtypes of neurons called Kenyon cells (**Supplementary Figure S6**). Dimension 4 differentiates between the central complex (middle embedding values) and the optic lobes (low embedding values), which is especially noteworthy because the central complex was not even imaged in the forager brain sample. Additionally, comparisons with GraphST and STAGATE show that SpaceExpress provides a more consistent alignment of these regions, underscoring its effectiveness in generating a shared coordinate system for samples under different conditions (**Supplementary Figure S7**).

**Figure 4.**
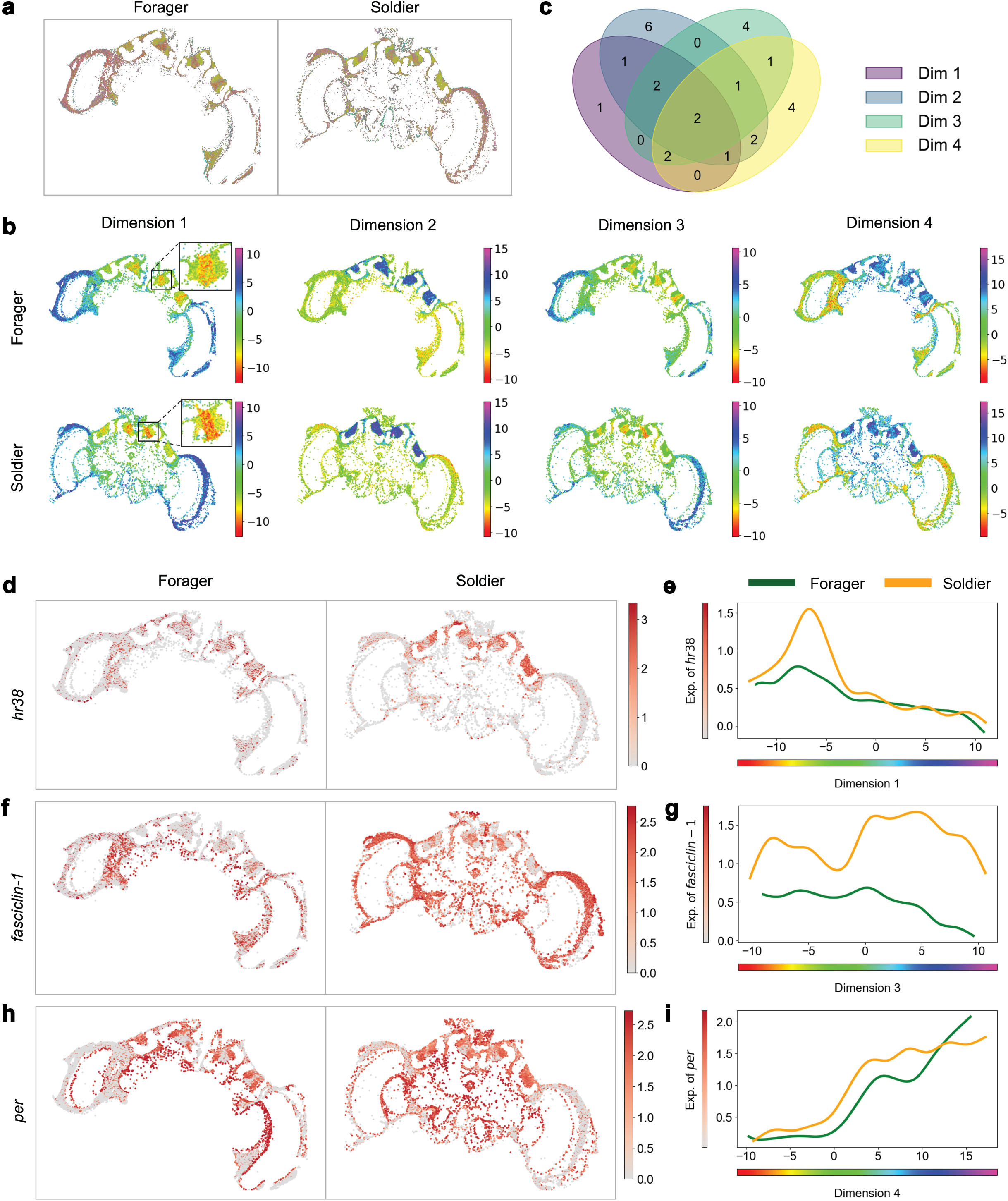
SpaceExpress identifies differential spatial expression of genes in honey bee brain. **(a)** Transcriptome map for forager and soldier honey bee brains. Each spot represents a transcript (n= 11,373 and 10,334 respectively) with color representing gene identity. **(b)** Visualization of SpaceExpress cell embeddings for the two honey bee brain samples, in all four dimensions. Each point corresponds to an individual cell at its spatial location, colored by the embedding value for each dimension. **(c)** Venn diagram showing the overlaps among sets of genes exhibiting Differential Spatial Expression (DSE) in each of the four embedding dimensions (Dim 1-4). **(d, f, h)** Spatial expression visualization of select DSE genes. The expression levels of *hr38* (d), *fasciclin-1* (f), and *per* (h) are shown. Cells with higher expression levels of each gene are colored in darker shades of red, revealing distinct patterns of spatial expression in different brain regions. **(e, g, i)** Gene expression versus embedding dimension plots. These plots display spline model fits for gene expression levels of *hr38* (e), *fasciclin-1* (g), and *per* (i) against the embedding dimension in which DSE was detected as significant. See text for how to interpret DSE based on gene-specific visualizations (d-i) and embedding dimension visualizations (b).

Based on these embeddings, SpaceExpress found 27 unique DSE genes at an FDR of 0.005 across all four embedding dimensions (**Figure 4c**). Our spline model allows for different average expression levels (see **Methods**), ensuring that these findings were not driven by overall differential expression (DE) of a gene between the two samples. To confirm this, we tested for overall differential expression of each gene using a Wilcoxon test, and found DE and DSE ranks of genes to be uncorrelated (**Supplementary Figure S8**).

A unique advantage of the SpaceExpress framework is that its embedding visualizations can be used along with its fitted spline models to interpret DSE patterns and derive biological insight. For each DSE gene, SpaceExpress produces a consolidated report with figures showing the spatial expression patterns in the samples, visualizations of the embedding dimension in which the DSE was detected, and a plot of the best-fit spline models showing how expression varies with the embedding dimension in each sample. We highlight a few of our results below.

Some genes show a similar spatial expression pattern across an embedding dimension between the two phenotypes, except in a localized region. This is evident in the dimension 1 spline plots for *hr38* (DSE FDR 1.8 × 10^−11^), where the expression in the soldier differs from the forager approximately between embedding values -7 and -4, where it shows a peak (**Figure 4e**). The embedding visualization of dimension 1 (Figure 4b) shows that these embedding values roughly correspond to the large Kenyon cells of the MB (Supplementary Figure S6), indicating that *hr38* is more spatially localized to these cells in the soldier than in the forager (**Figure 4d**). Corroborative of these findings, *in situ* hybridization has previously shown that spatial localization of this immediate early gene within the MB varies with aggressive responsiveness in honey bees, though overall expression levels do not change across aggressive or affiliative contexts^26^.

Some genes’ spatial expression patterns look different across most of the embedding dimension. For example, the dimension 3 spline plots for *fasciclin-1* (DSE FDR 0.0) show that in the soldier, the gene has higher expression for embedding values in the range 3 – 5 (blue in Fig. 4b) compared to values around −3 (green in Fig. 4b), while this trend is absent for the forager (**Figure 4g**). Given the embedding visualization of dimension 3 (Figure 4b), this suggests that *fasciclin-1* is spatially more expressed near the optic lobes than large Kenyon cells of MB in the soldier, in contrast to the pattern in the forager (**Figure 4f**). Interestingly, *fasciclin-1* is associated with synaptic plasticity^27^, which is important for behavioral differences.

We also found the circadian clock gene *per* to be DSE along dimension 4 (DSE FDR 2.6 × 10^−4^), and the associated visualizations (**Figure 4b,h,i)** reveal that its expression is concentrated in the MB in both brains, but more so in the soldier. We anticipate substantial variation in timekeeping strategies between soldiers and foragers: foraging behavior is gated by diurnal nectar and pollen availability^28^, whereas soldiers must always be prepared to launch an attack at the earliest indication of a threat. Therefore, variation in circadian-associated localization may indicate a mechanistic underpinning of phenotypic variation across soldiers and foragers.

### SpaceExpress Reveals Experience-Dependent Transcriptomic Gradients in the Mouse Visual Cortex

A common paradigm in transcriptomics is to identify molecular changes, e.g., DE genes, between two groups of replicates representing different biological conditions or treatments. SpaceExpress is the first tool to extend this influential paradigm to spatial transcriptomics in a statistically rigorous manner. The main idea is to include random effects terms into the SpaceExpress spline regression model (Methods).

We next applied SpaceExpress to map vision-dependent transcriptomic patterns with cellular resolution in the mouse primary visual cortex. For this, we used a previously published MERFISH dataset from postnatal day 28 (P28) mice^16^ (**Figure 5a**). This dataset consisted of coronal sections from normal-reared (NR) and dark-reared (DR) mice (two biological replicates per condition). DR mice were raised under normal conditions, but moved to the dark at P21, which marks the beginning of a critical period where development of the visual cortex is particularly sensitive to visual deprivation^29^. For each animal, one anterior and one posterior slice were profiled for a panel of 500 genes.

**Figure 5.**
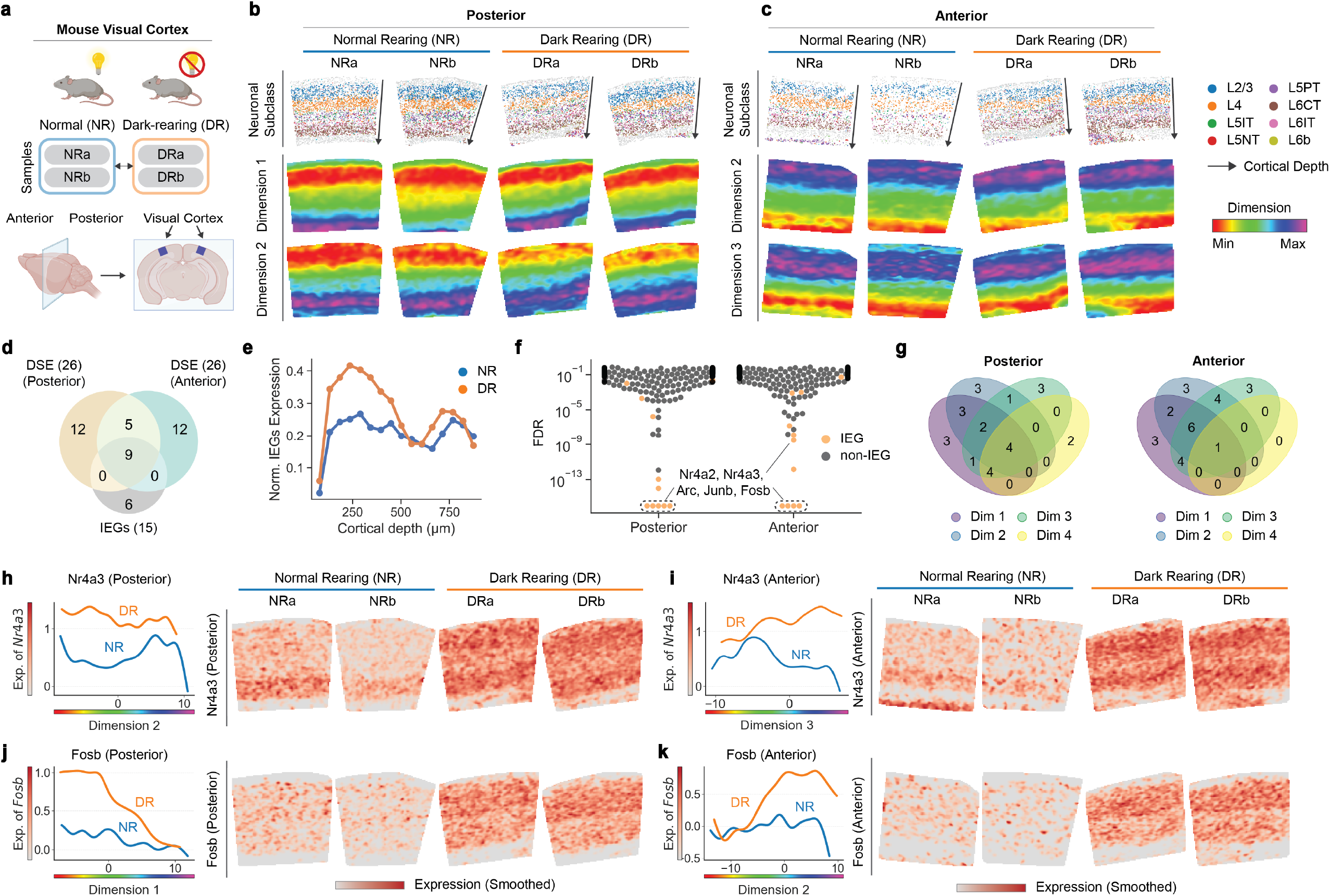
SpaceExpress analysis reveals vision-dependent spatial gene expression changes in the mouse visual cortex. **(a)** Schematic of experimental design. Primary visual cortex (V1) samples were collected from normally reared (NR) and dark-reared (DR) mice at postnatal day 28, including anterior and posterior slices (∼550 *μ*m apart). Each group includes two biological replicates (NRa, NRb, DRa, DRb). **(b, c)** Visualization of neuronal subclass annotation and SpaceExpress dimensions across (b) posterior and (c) anterior V1 samples. Top: neuronal subclass annotation overlaid on the tissue sections, with arrows indicating the direction of cortical depth. Bottom: Smoothed heatmaps generated using kernel density estimation, displaying the learned natural coordinate values as continuous color gradients for each dimension. Venn diagram showing overlap of differentially spatially expressed (DSE) genes in anterior and posterior analyses. Both regions yielded 26 DSE genes, with 14 shared genes, 9 of which are immediate early genes (IEGs). **(e)** Normalized expression of the 9 shared IEGs plotted as a function of cortical depth, averaged across replicates. DR samples show elevated expression between 100–500 *μ*m. **(f)** Beeswarm plot showing the FDR for each gene in posterior and anterior analyses. FDR for each gene is calculated by the minimum FDR values across all dimensions. IEGs are highlighted in orange. **(g)** Venn diagrams showing the number of DSE genes identified per dimension in posterior (left) and anterior (right) analyses. **(h-k)** Differential spatial expression of (h, i) Nr4a3 and (j, k) Fosb, two immediate early genes identified as DSE in both posterior and anterior analyses. Each panel shows, from left to right, spline model fits and spatial expression heatmaps for posterior or anterior samples. Spatial expression heatmaps are smoothed across replicates using kernel density estimation.

We conducted two separate analyses using SpaceExpress: one for the posterior V1 sections and one for the anterior V1 sections, each using all four P28 datasets (NR – replicates NRa, NRb, and DR – replicates DRa, DRb) to learn a shared, four-dimensional common coordinate system. In the posterior V1, dimensions 1 and 2 formed smooth gradients that closely tracked cortical depth along the pial-ventral axis, in all four replicates (**Figure 5b**). By contrast, in the anterior V1, dimensions 2 and 3 exhibited inverse gradients, with coordinate values increasing toward the pial surface (**Figure 5c**). These concordant embedding patterns confirm that SpaceExpress recovers the intrinsic laminar organization of V1 across rearing conditions^30^.

We next tested each gene for differential spatial expression (DSE) along each dimension, performing this analysis separately for the posterior and anterior datasets. We used the multi-replicate DSE analysis version of SpaceExpress (see Methods), which models variation across biological replicates and controls for sample-to-sample variation and heterogeneity in cell type composition. The number of DSE genes identified in each dimension for both analyses is summarized in **Figure 5g**. Aggregating across all four dimensions, both anterior and posterior analyses identified 26 DSE genes (**Figure 5d**), with 14 genes common to both regions. Notably, nine of these shared genes are canonical immediate-early genes (IEGs) that are known to be controlled by neuronal activity^31^. Using differential gene expression analysis, IEGs were previously reported to be upregulated under dark-rearing conditions across several neuronal subclasses in the visual cortex^16^ (**Figures 5d,e**); however, their condition-related difference in spatial pattern had not been quantified. To summarize their SpaceExpress-detected DSE pattern, we plotted the normalized expression of the 9 DSE IEGs against cortical depth (**Figure 5e**). This analysis revealed that IEGs are particularly upregulated in upper cortical layers in DR. This pattern is exemplified by the IEGs Nr4a3 and Fosb (**Figure 5h-k**). Under DR, Nr4a3 shows upregulation in low values of dimension 2 and in higher values of dimension 3 in posterior and anterior slices respectively (**Figures 5h,i**). (Note that dimension 2 in posterior slices increases with cortical depth, while dimension 3 in the anterior slices is anti-correlated with cortical depth.) Observed changes in Fosb are also consistent with specific upregulation in upper layers of the cortex (**Figures 5j,k**). As upper-layer cortical excitatory neurons have been suggested to be highly susceptible to sensory deprivation^32^, results from SpaceExpress are consistent with expected roles for IEGs in mediating sensory input-dependent development of upper-layer cortical neurons during the critical period.

### SpaceExpress pinpoints behavior-related changes in the mouse hypothalamic preoptic area spatial transcriptome

We next applied SpaceExpress to identify genes whose spatial expression patterns in the hypothalamic preoptic area of virgin male mice differ based on pup-exposure, which is associated with aggressive behavior. We analyzed MERFISH data from one control and one pup-exposed mouse, collected by Moffitt et al.^33^. The four-dimensional natural coordinate system learned by SpaceExpress (**Figure 6c**) captures meaningful tissue structure despite dramatic differences between the samples. Roughly, dimension 1 identifies the dorsal-ventral axis, dimension 2 highlights a gradient that increases from the medial preoptic area (MPA) and fornix (fx) to the periventricular zone (PV) and bed nucleus of the stria terminalis (BST), dimension 3 spans third ventricle (V3) and fx on one extreme and MPA on the other, while dimension 4 distinguishes the median preoptic nucleus (MPN) from the rest of the MPA. Importantly, these embeddings do not simply cluster known brain regions, they also identify continuous gradients within each region. For example, the dimension 3 embeddings exhibit a gradient within the V3 region. To support these embedding interpretations, we transferred expression-based brain region annotations from Li et al.^34^ using the tool SLAT^8^ (**Figure 6a**, Methods).

**Figure 6.**
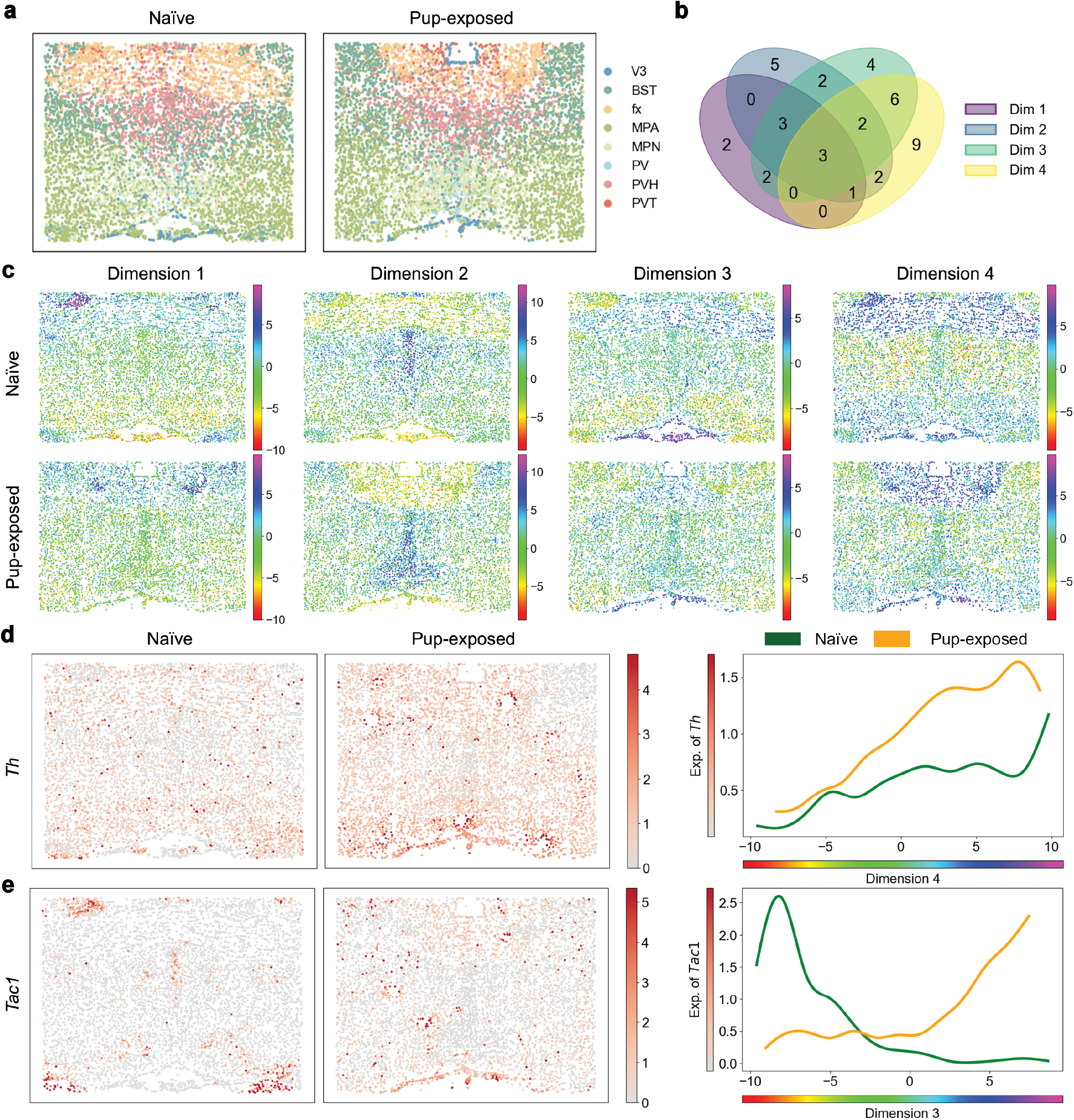
SpaceExpress identifies differential spatial expression of genes in mouse hypothalamic preoptic region related to pup-directed aggression. **(a)** Neuroanatomical annotation of cells in the hypothalamic preoptic region for naïve and a pup-exposed aggressive mouse, obtained computationally by mapping the cells to a reference tissue sample with manual annotations (Methods). Cells are color-coded according to their respective regional annotations, including the third ventricle (V3), bed nuclei of the stria terminalis (BST), columns of the fornix (fx), medial preoptic area (MPA), medial preoptic nucleus (MPN), periventricular hypothalamic nucleus (PV), paraventricular hypothalamic nucleus (PVH), and paraventricular nucleus of the thalamus (PVT). **(b)** Venn diagram illustrating the overlaps among sets of DSE genes across the four embedding dimensions (Dim 1 to 4). **(c)** SpaceExpress embeddings visualized across four dimensions for both the naïve and pup-exposed mouse brains. Each cell is represented by a point, positioned according to its spatial location and colored by the corresponding embedding value for each dimension. **(d, e)** Spatial expression visualization and gene expression versus embedding dimension plots for select genes. Shown are the expression levels (grey-red colormaps) of *Th* (d) and *Slc17a6* (e) in both naïve and pup-exposed samples, alongside spline model fits for gene expression along the respective embedding dimensions where significant DSE was detected.

Based on these embeddings, SpaceExpress identified 41 unique DSE genes at an FDR of 0.005 (**Figure 6b**). These include the gene *Th* (tyrosine hydroxylase), which is DSE with FDR ∼0 in dimension 3. **Figure 6d** and **Supplementary Figure S9** show that the contrast in *Th* expression between fx and V3 regions versus BST and MPN regions is stronger in the pup-exposed mouse compared to the naïve mouse. Th is crucial for neurotransmitter production, including dopamine and norepinephrine, which regulate emotional states including aggression^35,36^. V3 surrounds key nuclei in the hypothalamus that regulate aggression^37^ while fx is linked to emotional regulation through connections to hypothalamus and hippocampus^38^. The fx- and V3-specific difference in *Th* expression thus suggests catecholamine signaling as a contributor to aggression toward pups^39^. A similar case of DSE is observed for the gene *Tac1*, which exhibits elevated expression in the V3 region, especially in the pup-exposed mouse **(Figure 6e, Supplementary Figure S10**): its expression along dimension 3 has opposing trends in the two samples (FDR ∼ 0). *Tac1* encodes “substance P”, a neuropeptide that is a key modulator of neurotransmission in regions involved in aggression such as V3^40^, and our finding suggests that it may heighten aggression by amplifying the hypothalamic response to stressful stimuli such as pup-exposure. Many other genes well-known to be involved in aggression and parental behavior were also identified as DSE, including *oxytocin*^41,42^, *urocortin-3*^43^, and *galanin*^44^ (**Supplementary Figure S11**).

We also analyzed this dataset in the multi-replicate mode comparing three control and three pup-exposed mice and found additional DSE genes, e.g., *Gad1*, that were not identified above (**Supplementary Note 3**).

## DISCUSSION

SpaceExpress enables the identification and biological interpretation of genes whose spatial expression patterns correlate with phenotypic traits. SpaceExpress can compare the spatial structure of gene expression across samples of highly organized tissue, capturing the fine spatial features encoded in the tissue’s transcriptome. This contrasts with existing tools that operate at coarser resolutions such as spatial domains^5^, which can be used to identify differential gene expression in specific spatial domains across tissues^45-47^. It also differs from tools that study local spatial contexts, such as spatial “niches”^48^ and “microenvironments”^49^, which are more suitable for comparing tissues with repeating niche features such as tumor samples.

A possible alternative to SpaceExpress is to align tissue samples and compare gene expression between aligned cells. However, this strategy faces technical challenges posed by structural variations across tissue samples, which makes alignment difficult^50^, and it is hard to scale beyond pairwise comparisons (e.g., handling replicates) without treating one sample as a reference^51^. Our approach of leveraging positional information encoded in gene expression offers a more robust solution. Furthermore, it has the added advantage that we can visualize expression along the axes of a natural coordinate system adapted to intrinsic tissue structure and thus biologically interpret the statistically detected DSE. A recent method called “River”^11^ uses alignments between ST samples to describe their cellular locations in a common reference frame, and then uses a machine learning approach to identify any gene whose spatial pattern is informative of the sample’s phenotypic group. However, DSE detection by this strategy is prone to errors propagated from the initial alignment step, which can be challenging in many real scenarios^52^, does not provide a rigorous statistical test of difference, and is based on the assumption that every cell in one condition is distinguishable (in its spatial transcriptome) from every cell in the other condition.

SpaceExpress, like other ST analysis tools, combines transcriptomic and spatial information to obtain low-dimensional embeddings of cells. A unique property of these embeddings is that the expression vector fully determines the embedding. This property enables the tool to embed cells profiled in non-spatial single-cell experiments as well. It can thus be used for “spatial reconstruction”^9,10^ without explicitly mapping to an arbitrary reference sample as is done today. The ability to generalize to unseen spatial data is also key to the model’s robustness for learning natural coordinate systems (**Supplementary Figure S12**).

The statistical methodology of DSE detection in SpaceExpress is capable of associating spatial expression patterns with quantitative as well as discrete traits. We have described results for binary traits but the underlying statistical framework allows quantitative traits to be accommodated as well. The model is highly flexible and can optionally accommodate cell type information through additional terms in the model (Methods).

An important direction of future research is to systematically explore model complexity via tuning of hyperparameters, including the number of dimensions of the embedding space. Though the embedding space is meant to capture space (typically two- or three-dimensional), there can be value to using more than two embedding dimensions as each such dimension is meant to capture a biologically meaningful spatial pattern and there may be more than two such patterns, just as how the intrinsic dimension of a network object can be arbitrarily large.

## METHODS

### SpaceExpress Embeddings

The first step of SpaceExpress is to represent cells in a spatial transcriptomic (ST) dataset as low-dimensional vectors (“embedding” vectors or “embeddings”). Formally speaking, let the ST data set comprise a collection of cells *C* with each cell *c* ∈ *C* being described by a gene expression vector *g*(*c*) (of dimensionality *G* = number of genes) and a spatial location vector *x*(*c*) of dimensionality two or three. Thus, the data set may be denoted by *D*_*ST*_ = {*c, g*(*c*), *x*(*c*)}_*c*∈*C*_. Given *D*_*ST*_ and an integer *d*, the dimensionality of desired embedding vectors, SpaceExpress learns a function *F*: *R*^*G*^ → *R*^*d*^ that can map any given gene expression vector *g*(*c*) to its *d*-dimensional embedding *p*(*c*) such that for any two cells *c*_*i*_, *c*_*j*_ in *D*_*ST*_, the Euclidean distance between *p*(*c*_*i*_) and *p*(*c*_*j*_) reflects the spatial proximity between *c*_*i*_ and *c*_*j*_. In this sense, the embedding function *F*(*g*(*c*)) is said to “capture space”.

To make the previous definition precise, consider a *k* -nearest neighbor (“knn”) graph whose nodes are all cells *c* ∈ *C*, and a directed edge from *c*_*i*_ to *c*_*j*_ is included if and only if cell *c*_*j*_ is among the *k* nearest neighbors of cell *c*_*i*_ based on their physical locations *x*. Here, *k* is a hyperparameter of the method, which is chosen as the smallest integer for which the graph is a connected graph, i.e., there is a path from every node to every other node. We now define the distance *d*_*ij*_ between any two cells *c*_*i*_ and *c*_*j*_ as the shortest path distance between their corresponding nodes in the knn graph. The function *F* introduced above is learnt so as to minimize the loss function 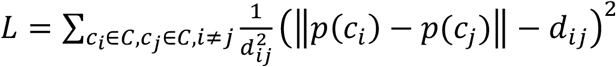, where *p*(*c*) = *F*(*g*(*c*)) and ‖⋅‖ represents the *L*_2_-norm of a vector. This loss function, henceforth called “KK” loss, is a special form of the Kamada-Kawai loss function used in network layout applications^53^. It is a sum over all pairs of cells (*c*_*i*_, *c*_*j*_) of the squared difference between *d*_*ij*_ (their spatial distance, as defined above) and ‖*p*(*c*_*i*_) − *p*(*c*_*j*_)‖ (Euclidean distance between their embedding vectors), but with each term being weighted by 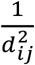, thus down-weighting errors made on more distant pairs of cells. The training of function *F*, also called the “model”, can be done on one ST data set, as explained above, or on multiple data sets; in the latter case, the loss function used is the sum over the KK loss on each data set. Once trained, the model may be used to map the cells of any new data set, using only their expression vectors *g*(*c*), to their corresponding embeddings *p*(*c*), without requiring their spatial locations. SpaceExpress determines the optimal value of *k* by iteratively increasing *k* from 4 to 20 and selecting the smallest *k* that results in a fully connected graph. If no such *k* is found within this range, it first generates a graph with the smallest *k* that minimizes the number of connected components, then connects the physically closest pairs of nodes from different components to achieve full connectivity. Specifically, for n disconnected components, it adds n−1 edges to connect them: it finds the closest cell pair straddling two disconnected components and connects them with an added edge, repeating this process until all components are mutually connected.

When training on multiple ST datasets, SpaceExpress performs joint training to learn a shared embedding function *F* that captures spatial relationships across all datasets. Suppose we have *n* datasets. For each dataset, the same number of cell pairs are randomly selected to ensure equal contribution from each dataset. The KK loss is computed separately for each dataset, resulting in individual losses *L*_*i*_ for *i* = 1,2, …, *n*. The total loss function used for training is the sum of the individual losses: 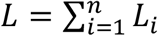. This approach allows the model to generalize across different samples and conditions, effectively integrating multiple datasets into a unified embedding space. In the implementation, during each training epoch, a fixed number of cell pairs from each dataset are randomly sampled to compute the KK loss to prevent the model from being biased toward larger datasets.

The function *F* that generates *p*(*c*) = *F*(*g*(*c*)) is a neural network with an input layer comprising *G* neurons (one for each gene’s expression value), an output layer with *d* neurons and a single hidden layer with *h* neurons (*h* is a hyperparameter), and two fully connected layers, for a total of *O*(*Gh* + *dh*) free parameters. Parameter optimization is implemented using Adam optimizer. The full collection of 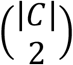 cell pairs is partitioned at random into *N*_*mb*_ equal subsets and these subsets are used as mini-batches in each epoch. Batch normalization is used as a form of regularization, and though we explored the use of dropouts and L1/L2 regularization, these features were not used in the final implementation as they did not yield significant improvement on unseen data. The embedding dimension was fixed to 4 throughout all analyses, as we observed minimal performance differences across a range of dimensions (4, 8, 16, 32). A detailed evaluation of dimensionality effects is provided in **Supplementary Figure S13**.

Preprocessing: Each dataset was preprocessed by filtering out genes expressed in fewer than 3 cells and cells expressing fewer than 3 genes. The data was normalized to a target sum of 10,000 (each cell by total counts over all genes) and log transformed. To minimize the influence of extreme values, gene expression values were capped at the 95th percentile. The user may optionally select to use either all genes or only the most highly variable genes (HVGs); the number of HVGs to use (*n*_*HVG*_) is a configurable parameter. If the user selects to use HVG-based analysis, SpaceExpress identifies the *n*_*HVG*_ highest variance genes in each sample, calculates the intersection of these HVGs across all samples, and creates a limited version of each sample’s data that includes only these genes, and scales each gene in each sample to unit variance and zero mean.

### Synthetic data sets

“Early embryo slice” data set: Each such data set comprises 3,915 cells placed uniformly in an elliptical area whose major and minor axes, called “anterior-posterior” (AP) or “dorso-ventral” (DV) axes, have lengths of 100 and 50 units, respectively (Figure 2a). The transcriptomic data synthesized spans 100 genes that fall into the following categories:

- AP genes (20): these include two “core” AP genes AP1 and AP2, whose expression in a cell with coordinates (*x, y*) is given by 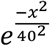and 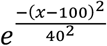respectively. Here, the major (AP) and minor (DV) axes defined the *x* and *y* coordinates of a cell, with *x* ranging from zero at the left (anterior) end of the major axis to one at the right (posterior) end, and *y* ranging from zero at the top (dorsal) end of the minor axis to one at the bottom (ventral) end. An additional 18 “derivative” AP genes are assigned expression values according to the formula *α*_*i*_ ⋅ *AP*_1_ + (1 − *α*_*i*_) ⋅ *AP*_2_, i.e., a weighted combination of AP1 and AP2 expression profiles. Each derivative gene uses a weight *α*_*i*_ chosen at random from the range. Note that all 20 AP genes have an “AP” expression pattern, i.e., the expression is identical in all cells at a fixed *x* coordinate (position along the AP axis).
- DV genes (20): these are assigned expression patterns similarly to the AP genes except that their expression varies along the DV axis and all cells with the same *y* coordinate have the same expression value for a DV gene. There are two core DV genes – DV1 and DV2 – with expression values given by 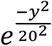and 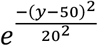respectively, and 18 derivative DV genes whose expression is a weighted combination of DV1 and DV2.
- APDV genes (20): each such gene is defined by the formula 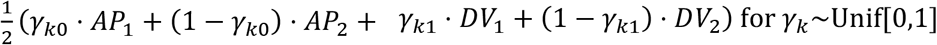, i.e., sum of an AP and a DV expression pattern, each being a weighted sum of the respective core genes.
- Randomly expressed genes (40): For each of these genes, we assign an expression pattern in each cell independently at random in the range [0,1]. These genes have no spatial pattern.

Once each gene’s expression level is assigned in each cell, an expression “count” is sampled from the Poisson distribution with parameter 1000*e*.

### Use of existing software

In this study, we utilized SEDR^54^, GraphST^15^, and STAGATE^14^ as benchmark methods to compare their embeddings with SpaceExpress. For SEDR, during the preprocessing step, if a dataset contained fewer than 200 genes, we removed the PCA step used for dimensionality reduction to 200. For STAGATE, we set each spot or cell to have six neighboring cells, following the original paper’s protocol. All other parameters were kept at their default settings as specified by the original authors.

### Evaluation of learned natural coordinate

We jointly trained SpaceExpress on two synthetic embryo samples to learn a shared four-dimensional coordinate system. To evaluate this coordinate system, we defined two complementary metrics: inter-sample agreement and natural coordinate fidelity. Inter-sample agreement measures whether cells with known correspondences across samples occupy nearby positions in the learned coordinate system (“embedding space”). Let 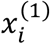 and 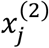denote the true spatial coordinates of cells *i* (in sample 1) and *j* (in sample 2), respectively, and let 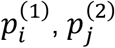 be their learned embedding coordinates. This spatial coordinatesystem is defined along a biologically meaningful axis common to both samples, specifically the anterior–posterior (AP) axis in the synthetic embryo (Figure 2b,d), cortical depth in the mouse visual cortex (Figure 2e), or histologically defined depth in the DLPFC samples (Figure 2g, Supplementary Figure S14). For each cell *i* in sample 1, we find its nearest neighbor in the other sample in embedding space:

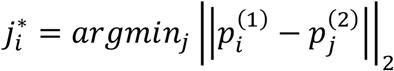

We then compute a normalized spatial score

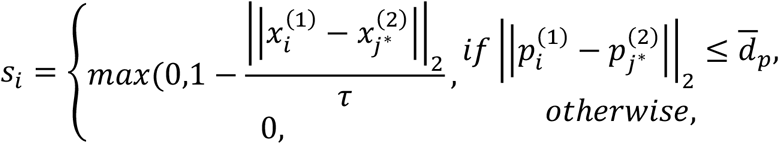

where 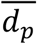 is the mean distance to the *k*-th nearest neighbor (*k*=10) in embedding space across all cells, and τ is the mean physical distance to the *k*′-th nearest neighbor (*k*^′^ = ⌊0.5N⌋) in true spatial coordinates.

The overall inter-sample agreement is then

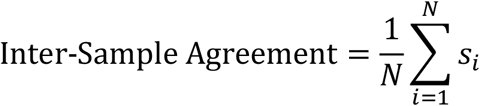

ranging from 0 to 1, with higher values indicating stronger agreement between samples. The condition 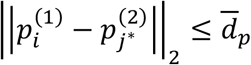 checks if the nearest neighbor of cell *i* in the other tissue sample is within a small distance. If this condition is met, we examine the physical distance 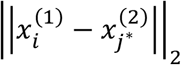between cell *i* and that nearest neighbor (such a distance is defined via the correspondence between cells in the two tissue samples, known as part of ground truth). The physical distance is converted to an “agreement” score by a linear transformation 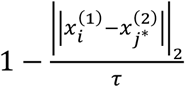, truncated at zero. This is the contribution of cell *i* to the overall inter-sample agreement. If the condition 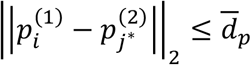 is not met, we ignore cell *i* in calculating inter-sample agreement. For the Drosophila embryo (Figure 2i), the coordinate system is defined by annotated spatial domains, so agreement is measured as the fraction of cells whose nearest neighbor from the other sample in embedding space belongs to the same domain.

Natural coordinate fidelity measures whether any single embedding axis closely aligns with the true spatial organization. For each sample, let *P* ∈ ℝ^N×D^ be the embedding matrix (with *N* cells and *D* dimensions), and let *z* ∈ ℝ^D^ denote the known natural coordinate of each of the *N* cells. Here, *z* is the position along the AP axis in the synthetic embryo (Figure 2b,d), cortical depth in mouse visual cortex (Figure 2e), or histologically defined depth in DLPFC samples (Figure 2g, **Supplementary Figure S14**. Natural coordinate fidelity is defined as:

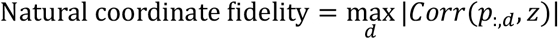

That is, we compute the absolute Pearson correlation between the natural coordinate of each cell and its learned coordinate, for each dimension *d* = 1, …, *D* separately, and the take the maximum of these correlations. This yields a value between 0 and 1, with higher values indicating that at least one embedding axis aligns closely with true natural coordinate. For the Drosophila embryo (Figure 2i), spatial organization is defined by annotated domains, so fidelity is measured as the fraction of cells whose nearest neighbor in embedding space falls within the same domain.

### Real data sets

We utilized SpaceExpress to analyze spatial transcriptomic datasets generated by various platforms, including 10x Visium, Stereo-seq and MERFISH. The human dorsolateral prefrontal cortex (DLPFC) dataset, generated using the 10x Visium platform, includes 12 sections from three individuals^17^. This dataset contains 33,538 genes, with the number of spots per section ranging from 3,460 to 4,789. The Drosophila embryo dataset, generated using the Stereo-seq platform^18^, includes 15 sections from the same embryo. For evaluation, we used three consecutive sections (IDs 6 to 8), with each section containing between 815 and 1,272 spots and 13,668 genes. The honey bee brain dataset was generated in-house using MERFISH, profiling 130 behavior-related genes in one soldier and one forager brain. The dataset includes 11,373 cells for the soldier and 10,334 cells for the forager; additional details of data generation are provided in the Methods section. The vision-dependent transcriptomic dataset of the mouse primary visual cortex^16^, generated using MERFISH, includes data from postnatal day 28 (P28) mice. It comprises two samples from dark-reared (DR) mice and two from normal-reared (NR) mice, with each sample providing data for both anterior and posterior regions, which are approximately 550 *μ*m apart, were profiled, yielding eight slices in total. This dataset contains 500 genes, with the number of cells ranging from 2,333 to 8,768. The mouse brain hypothalamic preoptic dataset^33^, which consists of six samples, each with 4,908 to 5,998 cells and 155 genes. For 1:1 comparison between naïve and pup-exposed samples, we used one male naïve sample (ID 5) and one male pup-exposed sample (ID 36), both at bregma 0.16. For the analysis of multiple biological replicates from different phenotype groups, we selected male naïve samples (IDs 5, 6, 7) and male pup-exposed samples (IDs 34, 35, 36), all from the same bregma (0.16).

### Synthetic data sets with differential spatial expression

To test the ability of SpaceExpress to learn “shared” coordinate systems, we synthesize a pair of “early embryo slice” data sets that are both organized using the AP and DV axes as coordinate axes, but exhibit a varying degree of divergence in that organization. Specifically, we follow the procedure outlined above to generate expression data for all but the “APDV” category of genes. Note that a gene in this category was assigned its expression value in a cell with coordinates (*x, y*) according to the formula 1⁄2 (*γ*_*k*0_ ⋅ *AP*_1_ + (1 − *γ*_*k*0_) ⋅ *AP*_2_ + *γ*_*k*1_ ⋅ *DV*_1_ + (1 − *γ*_*k*1_) *DV*_2_), where *γ*_*k*_∼Unif[0,1], i.e., a random combination of an AP pattern and a DV pattern. To simulate differential spatial expression of these genes, we now modify this procedure as follows: in the first data set, we zero out the contribution of the AP pattern, so the new formula is 1⁄2 (*γ*_*k*1_ ⋅*DV*_1_ + (1 − *γ*_*k*1_) ⋅ *DV*_2_), meaning that such an “APDV” gene exhibits an exclusively DV pattern. In the second data set, we weight the AP pattern by a constant *μ* ∈ {0.1,0.2, … 0.9}, so the new formula is 1⁄2 ((*μ* ⋅ *AP*_1_) + (*γ*_*k*1_ ⋅ *DV*_1_ + (1 − *γ*_*k*1_) ⋅ *DV*_2_)) and the “APDV” genes exhibit both a DV and an AP pattern with the strength of the latter being controlled by *μ*. All 20 APDV genes use the same value of *μ* in a data set. Thus, a pair of data sets comprises one embryo where APDV genes have zero AP pattern and one embryo where the AP pattern is weighted by a fixed value of *μ*. In this way we can generate pairs of data sets exhibiting varying degrees of DSE, by progressively increasing the value of *μ*.

### Benchmarking SpaceExpress and an Alignment-Based Baseline method for DSE Detection

We implemented an alignment-based approach as a complementary method for DSE detection that does not rely on learned coordinate systems. This approach aligns two ST tissue samples using PASTE2^50^ under default settings, with the overlap parameter set to the ratio of the number of cells between the two samples. A gene’s expression in aligned cell pairs (one cell from each tissue) is then examined, and the correlation coefficient is calculated. A perfect correlation of +1 indicates no differential spatial expression, while smaller values denote greater DSE, with -1 representing the strongest DSE. This is repeated for every gene and produces a ranked list of genes ordered by their DSE scores. On the other hand, SpaceExpress also produces a ranked list of DSE genes ordered by their FDR values.

To compare SpaceExpress and the alignment-based method, we first used either method on pairs of tissues from the same condition (intra-group comparison) to obtain their respective ranked lists of DSE genes. For either list, the 5th percentile DSE score (FDR for SpaceExpress-DSE and correlation for the alignment-based method) across all genes was noted. This was repeated for two intra-group tissue pairs (see Figure 3i,j) and the average of the 5th percentile DSE scores in these two analyses was designated as the DSE score threshold for the respective method. This threshold was then applied to designate DSE genes in inter-group comparisons. (We assume that intra-group comparisons provide a null model for DSE scores, so the threshold derived in this way controls false positive errors at 5% for both methods.) To evaluate robustness, the number of times each gene was designated as DSE across the four inter-group comparisons was noted. A robust method should consistently detect shared DSE genes across multiple inter-group comparisons. Additionally, River^11^ was employed as a competing method to evaluate DSE detection in comparison with SpaceExpress. For the prediction model in River, we defined each cell’s corresponding label as its condition group.

### Differential Spatial Expression analysis between two samples

#### Goal

In this section, we describe how SpaceExpress assesses differential spatial expression (DSE) of genes between two ST samples. We assume that the first step has learnt the shared coordinate system of the two samples, by embedding each cell in a latent Euclidean space. For each embedding dimension, SpaceExpress analyzes each gene separately and then combines the results via appropriate multiple-hypothesis testing procedure. As a result, we get a set of DSE genes for each embedding dimension.

#### Pre-processing

For a given gene, we log-transform the expression and then remove the cells that are beyond 4 standard deviations away from the mean. We drop a gene from the analysis if the transformed expression values are all zero in any group. We scale (without centering) the transformed expression within each group to have unit variance.

#### Linear Models

As mentioned in the goal, the analysis involves regressing each gene on each embedding dimension separately. For the sake of interpretation and statistical inference, we decided to use a linear model to fit gene expression against a given embedding dimension. However, a linear function of the embedding values will not be sufficient to capture the variation. Therefore, we use a spline regression strategy, where we create spline basis functions based on an embedding dimension. These spline basis functions are non-linear functions of the embedding dimension. Consequently, we can regress gene expression as a linear function of these spline basis functions. This approach allows us to retain the statistical interpretation advantages of a linear model while achieving a fit that is a non-linear function of the embedding. The generation of spline basis functions depends on a crucial parameter known as degrees of freedom. Higher degrees of freedom correspond to more non-linearity. We generate natural cubic spline bases with 300 degrees of freedom using the *ns* function from the *splines* package in R. For cell *i*, we denote the transformed and scaled gene expression by *Y*_*i*_, the generated spline basis vector by 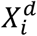 (*d* denotes the embedding dimension), and the group identity by *G*_*i*_. We fit the following two linear models using the *lm* package in R:

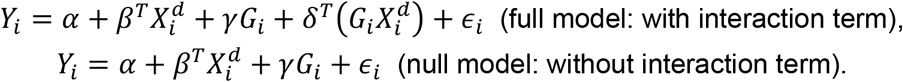

Here ∈_*i*_ are assumed to be independent Gaussian errors with zero mean. The interaction term δ represents the differential spatial expression between the two groups. Therefore, we test the significance of *δ* via a likelihood ratio test (LRT) between full and the null models. Furthermore, the *γG*_*i*_ term in the null model accounts for the mean expression difference between the two groups, ensuring that the observed effects are not simply due to overall differential expression (DE).

#### Multiple Hypothesis testing via Empirical Null method

For embedding dimension *d*, we collect the LRT based chi-squared test statistic for each gene. We estimate the empirical null distribution and then the false discovery rate (FDR) for each gene using the method described by Ren et al.^19^. As an implementation detail, the quantile parameter, q, for empirical null calculation that we use is 0.75.

### Multi-replicate Differential Spatial Expression analysis

Goal: In this section, we describe how SpaceExpress assesses differential spatial expression (DSE) of genes across two groups with multiple replicates.

#### Pre-processing

For a given gene, we log-transform the expression and then remove the cells that are beyond 4 standard deviations away from the mean. We drop a gene from the analysis if the transformed expression values are all zero in any group. We scale (without centering) the transformed expression within each group to have unit variance. Furthermore, we generate natural cubic spline bases with 50 degrees of freedom using the *ns* function from the *splines* package in R.

#### Linear Mixed Effects Models

]For cell *i* in replicate *j* we denote the transformed and scaled gene expression by *Y*_*ji*_, the generated spline basis vector by 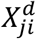 (here *d* denotes the embedding dimension), the group identity by *G*_*ji*_, and the cell type of cell *i* by *C*_*i*_. We fit the following linear mixed effects model using *lmer* package in R:

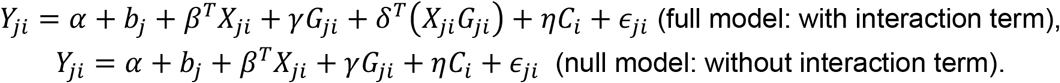

Here *b*_*j*_ represents zero mean random intercept term. Other coefficients *α, β, γ, δ*, and η are assumed to be fixed effects across replicates. Furthermore, ∈_*ji*_ are assumed to be independent Gaussian errors with zero mean. The interaction term *δ*^*T*^(*X*_*ji*_*G*_*ji*_) represents the differential spatial expression between the two groups. Therefore, we test the significance of *δ* via a likelihood ratio test between full and the null models. We can drop the *C*_*i*_ term and the coefficient η if we have just one cell type in the data or we do not want to capture the effect of cell types on the gene expression.

#### Multiple Hypothesis testing via Empirical Null method

For each embedding dimension, we collect the LRT based chi-squared test statistic for each gene. We estimate the empirical null distribution and then the false discovery rate (FDR) for each gene using the method described in Ren et al.^19^. As an implementation detail, the quantile parameter, q, for empirical null calculation that we use is 0.7.

### MERFISH data generation for honey bee brain

#### MERFISH sample preparation

Honey bee MERFISH samples were prepared using a slightly modified published method^33^. Specifically, honey bee brains embedded in OCT were cryo-sectioned at a thickness of 5-7 µm. To reduce spot density, samples were embedded in an expandable poly-acrylamide gel using a previously published method^55^.

#### MERFISH imaging

All images were acquired using a Zeiss Axiovert-200m widefield microscope (Carl Zeiss AG) located in the IGB core imaging facility. The sample was placed into a flow cell (Bioptechs, FCS2), filled with RNAse free 2x SSC, and connected to a lab built automated flow system. Briefly, computer-controlled valves (Hamilton, MVP/4, 8-5 valve) are used to select which solution was pulled across the sample by a computer controlled pump (Gilson, Minipuls 3). All systems are controlled by a custom designed Python script that can communicate with the microscope to start imaging or start flowing after an imaging round is done. In brief, a single round of imaging involves staining with fluorescently labeled readout probes (10% (v/v) ethylene carbonate (Sigma Aldrich), 0.1% Triton X-100 in 2x SSC, fluorescent DNA probe), washing with readout wash buffer (10% (v/v) ethylene carbonate, 0.1% Triton X-100 in 2x SSC) to remove unbound probes, and imaging buffer (5mM 3,4-dihydroxybenzoic acid (PCA; Sigma Aldrich), 2 mM trolox (Sigma Aldrich), 50 µM trolox quinone, 1:500 of recombinant protocatechuate 3,4-dioxygenase (rPCO; OYC Americas), was flowed into the flow cell prior to imaging to reduce photobleaching. A single quad band excitation filter (Chroma, ZET402/468/555/638x) and dichroic (Chroma, ZT405/470/555/640rpc-UF1) were used to image all samples. Excitation was provided by a 7 laser system (LDI WF, 89 North). Alexa Fluor 647 (Fisher scientific) labeled probes were excited using a 647 nm laser (0.5 W) with a ET700/75m (Chroma) emission filter. Atto 565 (Atto tec) labeled probes were excited using a 555 nm laser (1 W) with an ET610/75m (Chroma) emission filter. Fiducial beads were imaged with a 405 nm laser (0.3 W) with a ET440/40m emission filter. Samples were imaged with a 63x oil immersion objective (Carl Zeiss AG, 420782-9900-000), and focus was maintained between imaging rounds using Definite Focus (Carl Zeiss AG). After imaging is complete, a cleavage buffer (0.05 M TCEP HCl, adjusted to a pH of 7-7.2 using 1 N NaOH, in 2x SSC) was flowed across the sample to remove the fluorophores from the probes. The cleavage buffer was washed away using RNAse free 2x SSC (0.5 mL/minute for 10 minutes). This process was repeated for a total of 8 rounds of imaging. PolyA probes were stained after the final imaging round using the same method as described above.

#### MERFISH data processing

Individual FOVs were exported from czi format into 16 bit tiff format using Zen (Carl Zeiss AG) using the image export method. Images then were reformatted into image stacks by FOV and round. A modified copy of MERLIN^56^ was used to decode MERFISH spots. In brief, for each FOV, images from different rounds are aligned using fiducial beads that were imaged in each round. Aligned images are then normalized, decoded, and identified spots filtered using previously published methods^57^. Cell segmentation was done separately from MERLIN using Cellpose^58^ on PolyA and DAPI images for each FOV. To improve FOV alignment to neighboring FOVs, the DAPI channel was used with the restitching function found in Zen (Edge detection: on, minimal overlap: 5%, maximal shift: 15%, comparer: best, Global optimizer: best). Using the aligned images, segmented cells that cross FOV boundaries were merged into single cells, and global positions were generated for each spot. Spots are then assigned to cells based on their spatial coordinates. Spots were then filtered to remove any spot smaller than 3 pixels in size.

### Obtaining neuroanatomical annotation of mouse brain

We obtained the neuroanatomical annotations of the hypothalamic preoptic area in mouse brains based on the manual annotations in Li et al.^34^. These annotations were made in tissue sections at Bregma -0.14 mm from animal 1, and included eight domains (V3: third ventricle; BST: bed nuclei of the stria terminalis; fx: columns of the fornix; MPA: medial preoptic area; MPN: medial preoptic nucleus; PV: periventricular hypothalamic nucleus; PVH: paraventricular hypothalamic nucleus; and PVT: paraventricular nucleus of the thalamus). We used this annotated section as reference and aligned cells from sections of naïve mice (IDs 5, 6, 7) and pup-exposed mice (IDs 34, 35, 36) at bregma 0.16mm using SLAT^8^ with default settings. The domains from the reference cells were assigned to the mapped cells.

## DATA AVAILABILITY

Dorsolateral Prefrontal Cortex (DLPFC) Dataset: The 10x Visium data used in this study is available at https://github.com/LieberInstitute/spatialLIBD.git^17^. Drosophila Late-Stage Embryo Dataset: Stereo-seq data can be accessed at https://db.cngb.org/stomics/flysta3d/^18^. Mouse Hypothalamic Preoptic Area Dataset: MERFISH data was obtained from Moffitt et al.^33^, available at https://datadryad.org/stash/dataset/doi:10.5061/dryad.8t8s248. Honeybee Dataset: Data is available at https://doi.org/10.13012/B2IDB-5536668_V1.

## CODE AVAILABILITY

SpaceExpress is implemented in Python, and the code is available at https://github.com/YeojinKim220/SpaceExpress. The package utilizes the AnnData format and can be installed by following the instructions on the GitHub page. A tutorial demonstrating SpaceExpress with data from the mouse hypothalamic preoptic region is also provided^33^.

## ACKNOWLEDGEMENTS

We thank Anurendra Kumar for valuable discussion, and Seokjin Yeo and Sunny Sun for image segmentation, Marisa Asadian for help for imaging, and Adithya Karthik for assistance with alignment analysis. Funding: This work was supported by the National Institutes of Health (R35GM131819 to S.S., R35GM147420 to H.-S.H, R21HG013180 to S.D.Z. and H.-S.H.), Georgia Institute of Technology (Wallace H. Coulter Distinguished Faculty Chair: S.S.), and the European Union’s Horizon 2020 Research and Innovation Program under ERC-2017-StG Grant Agreement 757583 (Brain2Bee; to Jennifer L Cook and Gene E. Robinson). Facilities: We acknowledge Core Facilities at the Carl R. Woese Institute for Genomic Biology for their microscope and staff support.

