## Supplementary Figure for "SpaceExpress enables differential spatial transcriptomics with natural coordinate systems"

**Supplementary Note 1**

**SpaceExpress embeddings accurately capture space in synthetic and real data sets**

The first step of SpaceExpress is to embed cells of the given ST sample(s) in a new, intrinsic coordinate system. We evaluated the extent to which these embeddings can accurately capture the relationship between expression and location in synthetic ST datasets. We synthesized an "early *Drosophila* embryo slice" with cells laid out uniformly in an elliptical area, with a subset of genes varying in expression along the longer ("anterior-posterior" or "AP") axis and/or shorter ("dorso-ventral" or "DV") axis (**Supplementary Figure S15a**, **Supplementary Table S1**). We used SpaceExpress to map each cell to a 4-dimensional embedding vector. To assess the accuracy of these embeddings, we calculated the Jaccard Similarity (JS) score between the nearest neighbor graph of cells based on their Cartesian coordinates (the "x-knn" graph) and their embeddings (the "p-knn" graph) (**Supplementary Figure S15b**). To evaluate how well a given set of embeddings ($p(c)$ for all cells $c$ in a data set) capture space, we construct a $k$-nearest neighbor graph based on the locations $x(c)$ of cells, called the "x-knn" graph, and a $k$-nearest neighbor graph based on the embeddings $p(c)$, called the "p-knn" graph. In both cases, Euclidean distances are used to find nearest neighbors. We then calculate the Jaccard similarity score $JS($x-knn, p-knn) as $\frac{|E_x \cap E_p|}{|E_x \cup E_p|}$, where $E_x$ and $E_p$ are sets of edges (cell pairs) in the x-knn and p-knn respectively. To evaluate how well embeddings capture gene expression, we adopt a similar procedure as above, now constructing a $k$-nearest neighbor graph based on the gene expression vectors $g(c)$ of cells ("g-knn" graph), but using cosine similarities instead of Euclidean distances to define nearest neighbors, on account of the high dimensionality of expression vectors. We then calculate the Jaccard similarity score $JS$(g-knn, p-knn) as $\frac{|E_g \cap E_p|}{|E_g \cup E_p|}$, where $E_g$ is the set of edges (cell pairs) in the g-knn. We observed the JS score (theoretical range of 0-1) to be 0.45 on average (**Supplementary Figure S15c**), significantly greater than the random expectation of 0.001 as well as that of state-of-the-art ST cell embedding approaches like STAGATE, SEDR and GraphST (Supplementary Figure S15c). A Principal Components Analysis (PCA) visualization of the cell embeddings (**Supplementary Figure S15d, Supplementary Figure S16**) and visualizations of the learnt dimensions using colormaps (**Supplementary Figure S15e, Supplementary Figure S17**) provide further evidence that our embeddings accurately capture the AP and DV axes and thus learn the intrinsic coordinate system of this data set.

A tissue's spatial organization often comprises a set of discrete spatial domains or regions[1]. To test our model's ability to capture this type of organization, we synthesized a "domain grid" ST data set with nine different spatial domains arranged in a square grid (**Supplementary Figure S15f**), each domain distinguished by a marker gene's expression**.** Each such data set comprises 900 cells placed uniformly in a 30 x 30 grid, organized as 3 x 3 = 9 "regions" of 10 x 10 = 100 cells each (Supplementary Figure S15f). Each region is "marked" by a gene that is assigned an expression level of 1 in all 100 cells of that region and 0 in all remaining 800 cells. Thus, there are nine marker genes G1, G2, …, G9, corresponding to the nine regions respectively. An additional 91 genes are assigned random expression levels selected uniformly from $[0,1]$ independently for each cell. Once expression values have been assigned in this manner, an expression "count" is sampled from the Poisson distribution with parameter equal to 1000 times that assigned value.

SpaceExpress embeddings (JS score of 0.8 on average) were substantially more accurate than other methods (**Supplementary Figure S15g,h, Supplementary Figure S18**). We also assessed the extent to which SpaceExpress embeddings capture expression (**Supplementary Figure S19**) and observed that, compared to other embedding methods, they reflect spatial proximity more strongly than expression similarity. This underscores the different trade-offs made by different cell embedding methods when integrating expression and spatial information.

We next undertook similar assessments on real ST datasets obtained using different technologies. Our first evaluation was on the commonly analyzed dorsolateral pre-frontal cortex (DLPFC) data obtained by Maynard et al.[2] using 10X Visium technology. We analyzed 12 different samples (from 3 different donors) separately, and observed significantly higher JS scores for SpaceExpress embeddings than STAGATE, SEDR, and GraphST (**Supplementary Figure S20a, Supplementary Data S3**). In all samples, one of the learnt dimensions captures a transverse axis through the cortex layers**,** despite substantial differences in tissue spatial structure (**Supplementary Figure S20b,c**, **Supplementary Figure S21)**. In an additional evaluation motivated by the layered structure of these tissue samples, we validated that (a) cells in more spatially separated layers are embedded more distally to each other than cells in proximal layers, and (b) cells in the same layer are embedded more proximally to each other if they are located nearby in physical space; in both regards SpaceExpress is competitive with or better than other methods (e.g., see **Supplementary Figure S22, Supplementary Figure S20d** for results on sample ID 151507). We also evaluated the tool's embedding step on 16 samples of Drosophila late stage embryo (E14-16) profiled via STEREO-seq technology (Wang et al.[3]). Once again, we noted significantly higher JS scores than those of other methods (**Supplementary Figure S20e**) and a clear ability to capture the spatial organization of different regions in the embryo (**Supplementary Figure S20f,g**). In summary, the evaluations reported here confirmed the ability of SpaceExpress embeddings to capture the spatial location of cells based on their transcriptome, across a variety of ST data sets.

### Supplementary Note 2

#### SpaceExpress embeddings generalize to unseen tissue samples

The intrinsic coordinate system learnt by SpaceExpress is capable of describing cellular locations across multiple spatial tissue samples. To demonstrate this ability, we show here that an embedding function trained on one tissue sample can predict spatial location from expression in an unseen tissue of a similar type. Specifically, we trained SpaceExpress embedding functions on a DLPFC sample (ID 151507) and used the trained model to embed cells of a different ("test") sample from another donor (**Supplementary Figure S23)**, without utilizing spatial information about them. We found these embeddings to successfully capture separations between layers (**Supplementary Figure S24a**) as well as intra-layer distances between cells (**Supplementary Figure S24a**) in the test sample. A visualization of the embedding dimensions for the test sample **(Supplementary Figure S25**) shows that they represent spatial directions analogous to those in the training sample. (See **Supplementary Figure S26** for additional tests.) The ability of SpaceExpress embeddings to capture spatial organization of the test sample is superior to that of an advanced spatial reconstruction tool called TANGRAM[4] (**Supplementary Figure S24b**). We conducted similar tests using the Drosophila embryogenesis data set, training SpaceExpress on the central slice, S08 (**Supplementary Figure S20f**), and using it to assign spatial embeddings to other slices (**Supplementary Figure S24c,d**) without accessing spatial information from the test slice. As shown in **Supplementary Figure S24e** and **Supplementary Figures S27**, the spatial organization of cell types in these embryonic slices is well captured by the SpaceExpress embeddings. These tests demonstrate that the SpaceExpress cell embedding model captures space in a way that transcends the specific spatial coordinate system and cellular arrangements of the given tissue and is relevant even for other specimens of the tissue type. We note that no existing tool is capable of such generalizability: current ST embedding tools[1, 5, 6] require spatial location information to embed cells, while tools like TANGRAM[4] are intended to *map* a non-spatial scRNA-seq sample onto a reference spatial sample.

### Supplementary Note 3

#### DSE analysis on multiple replicates of mouse hypothalamic preoptic region from two behavioral groups

We applied SpaceExpress to find DSE genes across three control mice and three pup-exposed mice collected by Moffitt et al.[7], using the multiple replicate mode. Here, we included cell type as an additional covariate of the model, to account for the potential confounding effect that varying cell types may have on spatial expression patterns (Methods). **Supplementary Figure S28** visualizes the common four-dimensional intrinsic coordinate system learned by SpaceExpress. We found 19 DSE genes at an FDR of 0.001, including well-known genes such as *oxytocin* and *urocortin-3*. (The FDR threshold here is higher than above because of the much smaller sample size – number of replicates rather than number of cells.) For instance, the *Oxt* (oxytocin) gene is DSE in dimension 4 (FDR ~ 0), with the two groups of replicates diverging in their expression levels for embedding values greater than 0 (**Supplementary Figure S29a**). This roughly corresponds to the BST and PVH regions, as seen in **Supplementary Figure S28** and **Supplementary Figure S30**, with the pup-exposed mouse brains exhibiting lower *Oxt* expression than naïve mice. This is consistent with recent work demonstrating that oxytocin activation in the PVH region inhibits pup-directed aggression[8] and with the known role of the BST region in such behavior[9].

In addition, our multi-replicate analysis found several genes that were not DSE in the previous no-replicate analysis, e.g., Gad1 (FDR $2.0 \times 10^{-4}$) in dimension 1. Visualizations of spline plots (**Supplementary Figure S29b**) and the embedding dimension (**Supplementary Figure S28**) show that in pup-exposed mice *Gad1* expression drops and then rises between embedding values of -5 and 0, indicating that *Gad1* is high in the V3 and PV, high in the VLPO, and lower as cells get further from these two structures. In naïve mice *Gad1* rises continuously between embedding values -5 and 0 before also dropping at 0. Gad1 is a glutamate decarboxylase involved in synthesizing the inhibitory neurotransmitter GABA[10], suggesting that differing levels of inhibitory circuit activity in the regions around the VLPO may be related to pup-directed aggression. This finding is not visually obvious and requires all three replicates in each group, highlighting the unique advantages of an interpretable quantitative framework for capturing and summarizing differential spatial expression across multiple biological replicates.

| Gene | Equation |
|---|---|
| $AP_1$ | $e^{-\frac{x^2}{40^2}}$ |
| $AP_2$ | $e^{-\frac{(x-100)^2}{40^2}}$ |
| $DV_1$ | $e^{-\frac{y^2}{20^2}}$ |
| $DV_2$ | $e^{-\frac{(y-50)^2}{20^2}}$ |
| $AP_{3-20}$ | $\alpha_i \cdot AP_1 + (1 - \alpha_i) \cdot AP_2$ for $\alpha_i \sim$ Unif[0,1) |
| $DV_{3-20}$ | $\beta_j \cdot DV_1 + \left(1 - \beta_j\right) \cdot DV_2$ for $\beta_j \sim$ Unif[0,1) |
| $APDV_{1-20}$ | $\frac{1}{2}(\gamma_{k0} \cdot AP_1 + (1 - \gamma_{k0}) \cdot AP_2 + \gamma_{k1} \cdot DV_1 + (1 - \gamma_{k1}) \cdot DV_2)$ for $\gamma_k \sim$ Unif[0,1) |
| $R_1$ | $R_1$~Unif(0,1) |
| $R_2$ | $R_2$~Unif(0,1) |
| $R_{3-40}$ | $\delta_l \cdot R_1 + (1 - \delta_l) \cdot R_2$ for $\delta_i \sim$ Unif[0,1) |

**Supplementary Table S1.**

Equations of simulated gene expression for synthetic "early *Drosophila* embryo slice". $x$ and $y$ represent the coordinates of a cell along the anterior-posterior (AP) axis and dorso-ventral (DV) axis respectively. The "core" AP genes ($AP_1, AP_2$) and DV genes ($DV_1, DV_2$) have gradients of expression in opposing directions along their respective axes. Remaining AP genes ($AP_{3-20}$) and DV genes ($DV_{3-20}$) have an expression profile that is a linear combination of the two core genes along their respective axes. Each "APDV gene" has its expression pattern generated as a linear combination of the AP and DV patterns. Apart from these 60 genes (20 AP, 20 DV, 20 APDV), the remaining genes ($R_{1-40}$) consist of two genes ($R_1$ and $R_2$) with randomly assigned expression values, and the expression patterns of the other genes ($R_{3-40}$) are generated as a linear combination of these two. For all 100 genes, the above equations are used to generate a "clean" expression pattern, to which noise is added through a Poisson distribution.

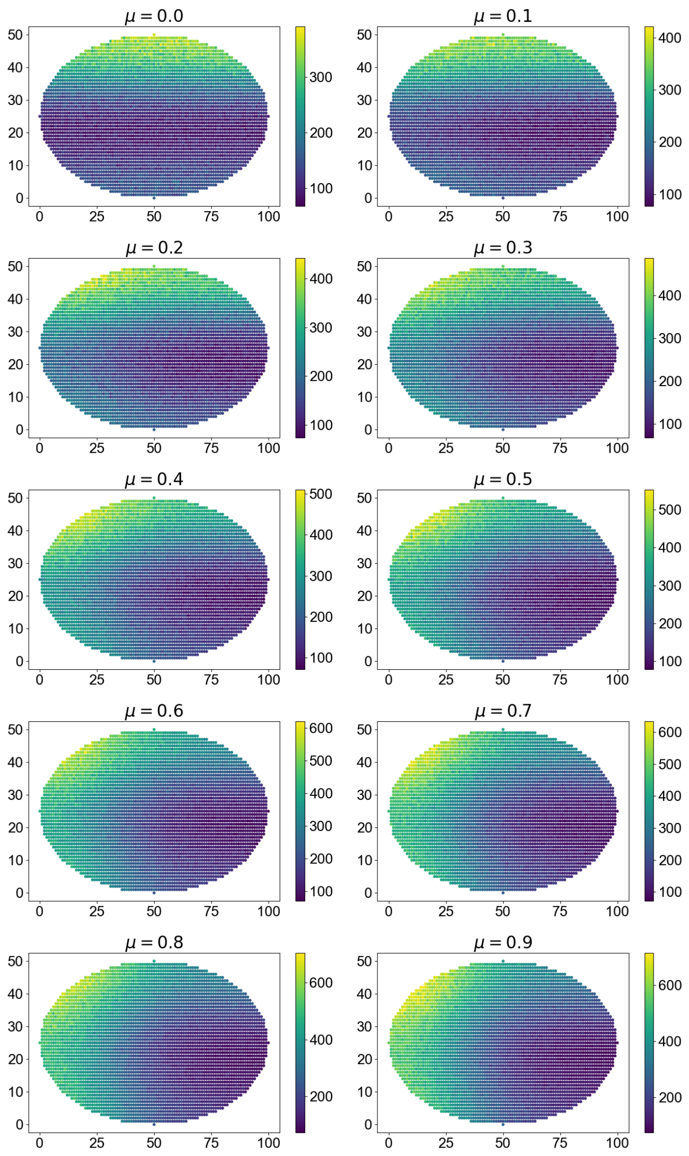


### Supplementary Figure S1

Example expression pattern of an “APDV” gene at different values of the parameter $\mu$, which controls the weight of the AP pattern, with μ set to 0 (purely DV pattern) to 0.9 (nearly equal combination of AP and DV patterns).

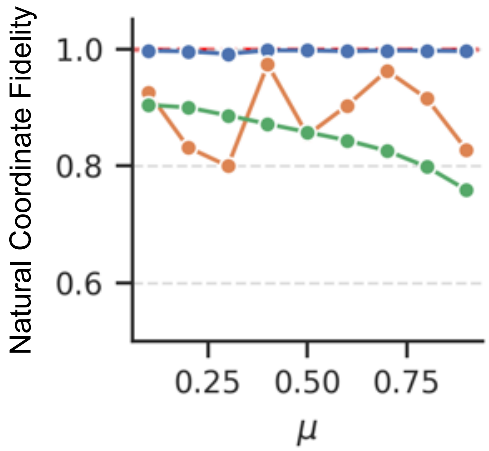


**Supplementary Figure S2**

Natural coordinate fidelity across the dorsal-ventral (DV) axis for synthetic embryo samples, shown across varying values of $\mu$. This analysis complements the AP-axis results shown in Figure 2b (right), with DV axis now treated as the “ground truth” natural axis. SpaceExpress consistently achieves scores near 1.0 regardless of the level of variation in gene expression patterns, indicating robust preservation of biologically meaningful structure. STAGATE shows moderate variability in fidelity depending on $\mu$, while GraphST steadily declines. The red dashed line marks the theoretical maximum score of 1. These results confirm that SpaceExpress reliably maintains spatial organization even as expression patterns diverge.

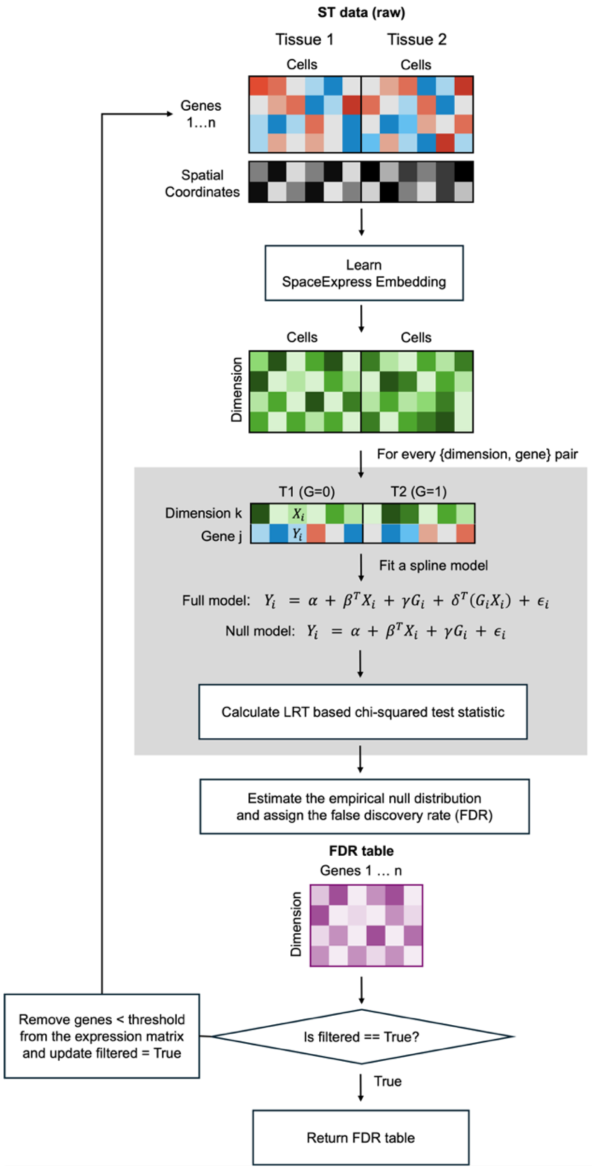


**Supplementary Figure S3**

Schematic description of DSE detection pipeline in SpaceExpress. Spatial transcriptomic (ST) data from two tissues are used as input. The method begins by learning a low-dimensional embedding that encodes the spatial structure of cells in each tissue. For each dimension-gene pair, a likelihood ratio test (LRT) is performed to compare a full model (which includes an interaction term) against a null model (without the interaction term). The LRT statistic is used to compute a false discovery rate (FDR) for each gene-dimension pair. Genes with FDR below a predefined threshold (in any dimension) are filtered out of the expression matrix for both tissues, and the process is repeated on the modified ST data. The final output is an FDR table. Note: the genes identified as DSE in first round are tested for DSE again in the second round.

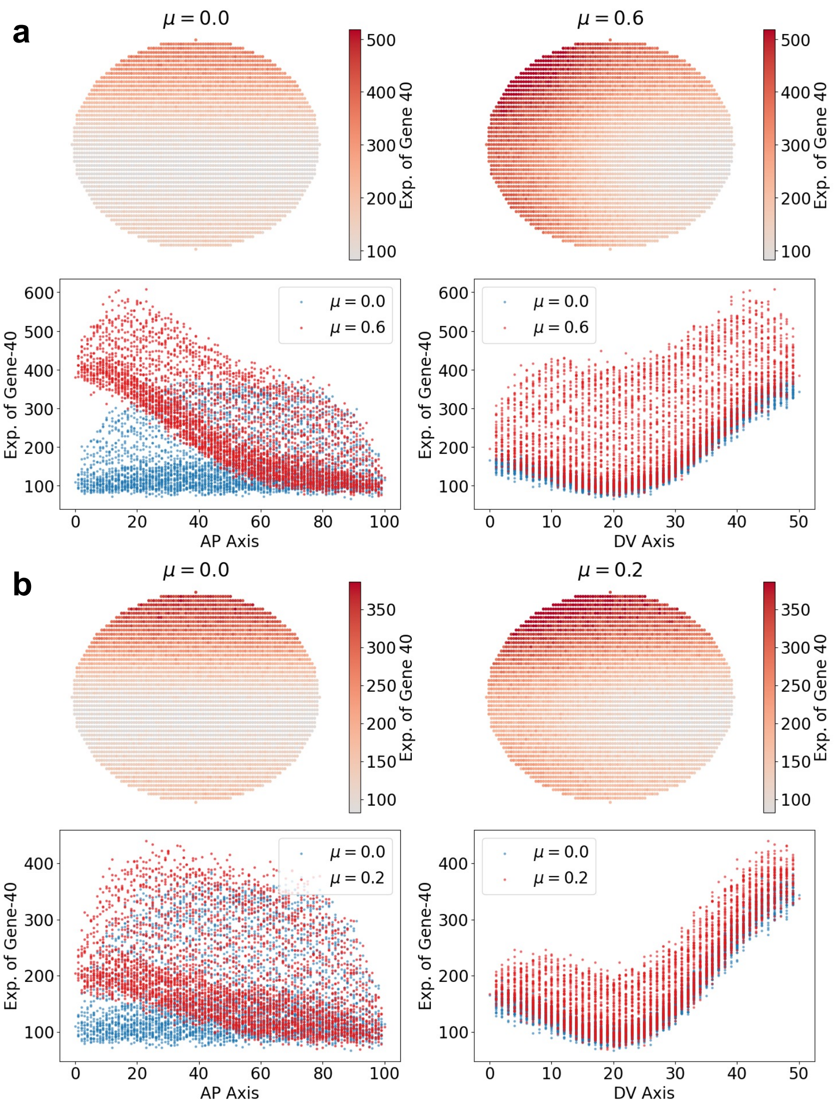


### Supplementary Figure S4

(a, c) Same as Figure 3f, this panel visualizes APDV gene expression patterns in two synthetic early embryos. In one embryo, the parameter $\mu$ is set to 0, representing a purely DV pattern, while in the other, $\mu$ is set to 0.6 (a) and 0.2 (c), representing a mixed AP and DV pattern. (b, d) Gene expression values from both embryos (blue and red) plotted along the AP and DV axes, showing differences in expression between the samples on the AP and DV axis.

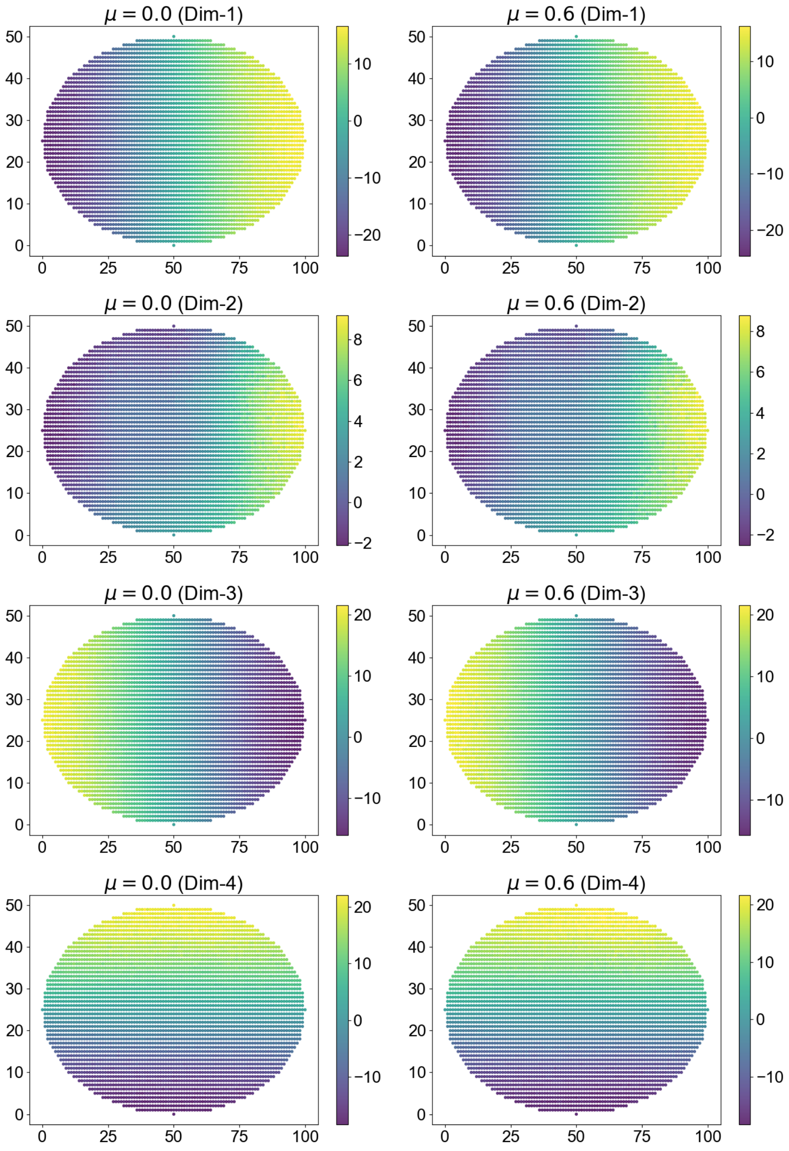


**Supplementary Figure S5**

Visualization of jointly trained SpaceExpress embedding dimensions in the two synthetic early embryos with $\mu$ set to 0 and 0.6.

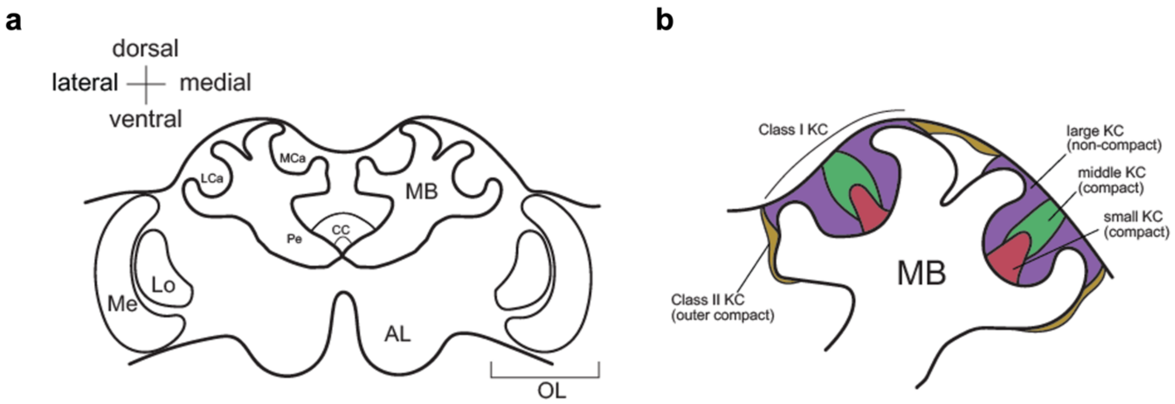


**Supplementary Figure S6**

Schematic of the honey bee adult worker brain. (a) Major bilateral neuropils such as the mushroom bodies (MB), optic lobes (OL), antennal lobes (AL), and central complex (CC) are shown. Within the MB, the lateral (LCa) and medial (MCa) calyx, as well as the peduncle (Pe) are identified on the left hemisphere of the schematic, as well as the lobula (Lo) and medulla (Me) of the OL; the outermost layer of the OL, the lamina, is not shown. (b) Expanded depiction of the MB from the right hemisphere in (a); Class I and II Kenyon cells (KC) are shown, and Class I KC are further divided into large, middle, and small cellular subtypes. Compact or non-compact cell patterning for each subregion is listed below each subregion's name.

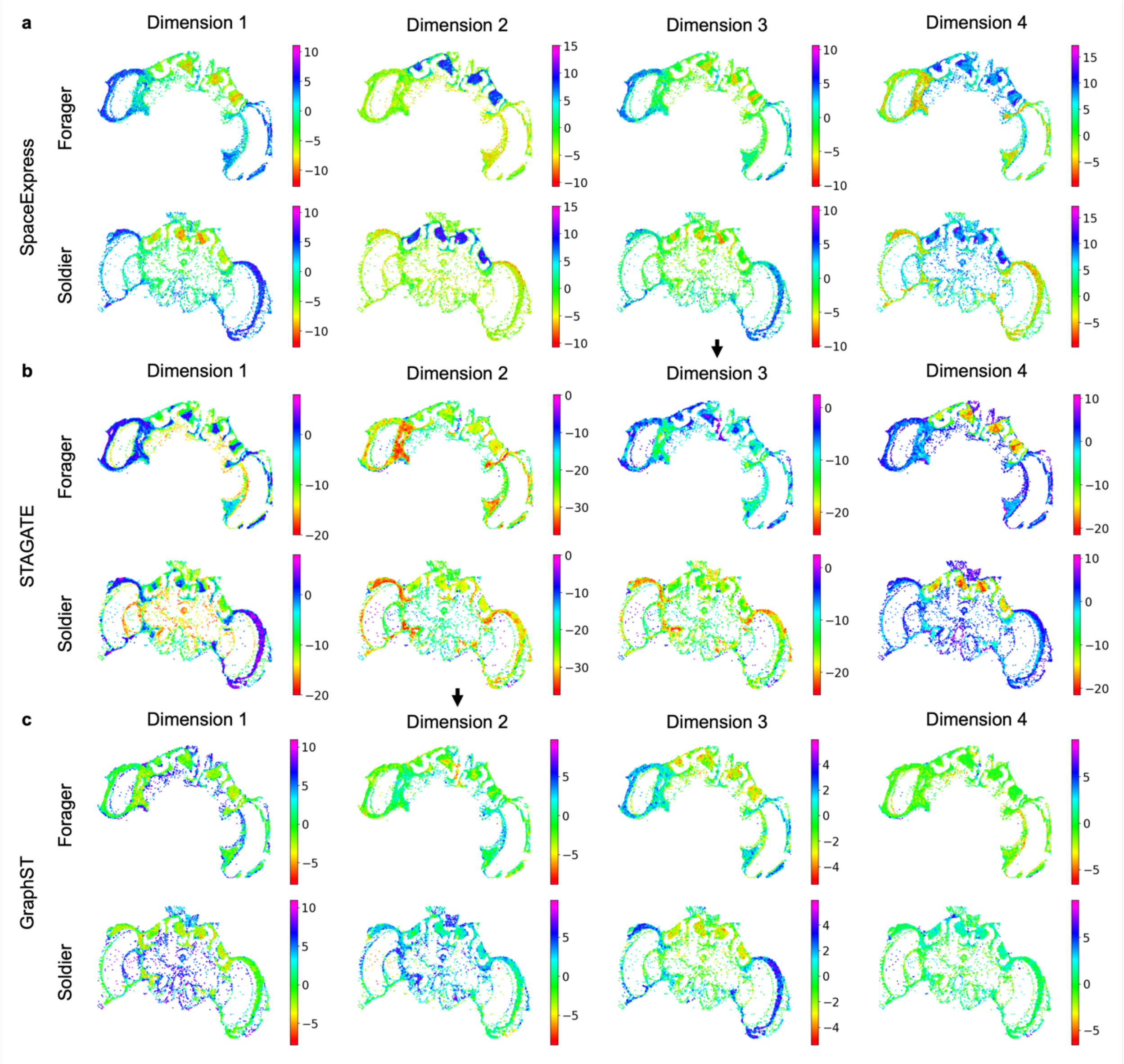


**Supplementary Figure S7**

Comparative assessment of approaches to learning shared coordinate systems between tissues. Shown are colormap visualization of cell embeddings generated by SpaceExpress (a), STAGATE (b), and GraphST (c) for two honeybee brain samples (forager and soldier) across four embedding dimensions. Each point corresponds to an individual cell at its spatial location, colored by the embedding value for the respective dimension. (Note: dimension “i" (i=1,2,3,4) of different methods, arranged in a column, have no correspondence to each other.) This figure helps assess the ability of each method to generate a shared coordinate system across samples from different biological conditions. Establishing a shared coordinate system is essential because each dimension serves as an intrinsic axis, allowing cells in different samples to be positioned and compared along the same coordinates. This alignment enables direct comparisons of gene expression based on each embedding dimension, ensuring that cells from corresponding anatomical regions (even if they come from different samples) occupy similar coordinates and can be compared reliably. For consistency, SpaceExpress and STAGATE embeddings were configured to be four-dimensional by construction, and GraphST embeddings (X-dimensional) were reduced to four dimensions using PCA. **(a)** SpaceExpress effectively captures the underlying anatomical structures of honeybee brains, assigning similar embedding coordinates to corresponding regions in both the forager and soldier samples, for each of the four dimensions. **(b)** STAGATE demonstrates some inconsistencies in creating a shared coordinate system. For instance, in Dimension 3, the forager brain has embedding values that cluster between -10 and 0, whereas the soldier spans from -20 to 10, suggesting a discrepancy of coordinates assigned globally to one brain versus the other. **(c)** GraphST embeddings also show limitations when used as a common coordinate system. For instance, in Dimension 2, forager brain cells are mapped to coordinates between -5 and 0, while soldier brain cells cluster between coordinates 0 and 5, indicating that analogous regions in the two samples are not mapped to a shared region in the learnt coordinate space. Furthermore, Dimension 4 lacks variation across different regions within each sample, rendering it ineffective as a meaningful coordinate system. Overall, these results underscore the effectiveness of SpaceExpress in generating a consistent embedding framework for cells across samples under different conditions, an essential capability for accurately comparing gene expression profiles across different biological conditions.

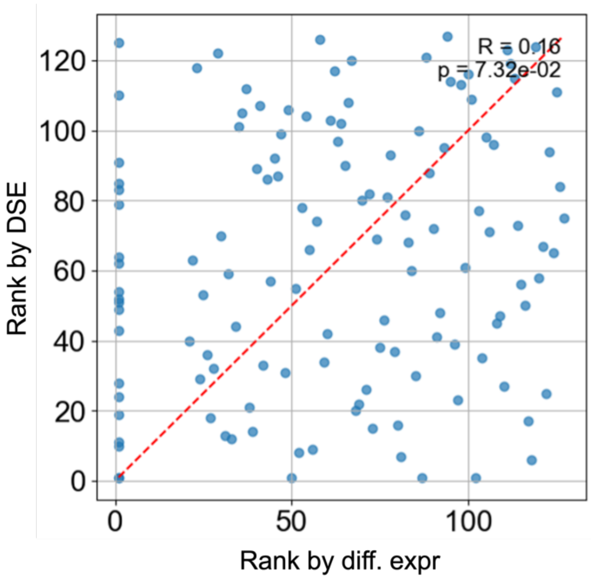


**Supplementary Figure S8**

Scatter plot showing the ranks of genes under differential expression analysis (FDR from Wilcoxon test, x-axis) and under differential spatial expression analysis (FDR from SpaceExpress, y-axis), for honey bee brain data set.

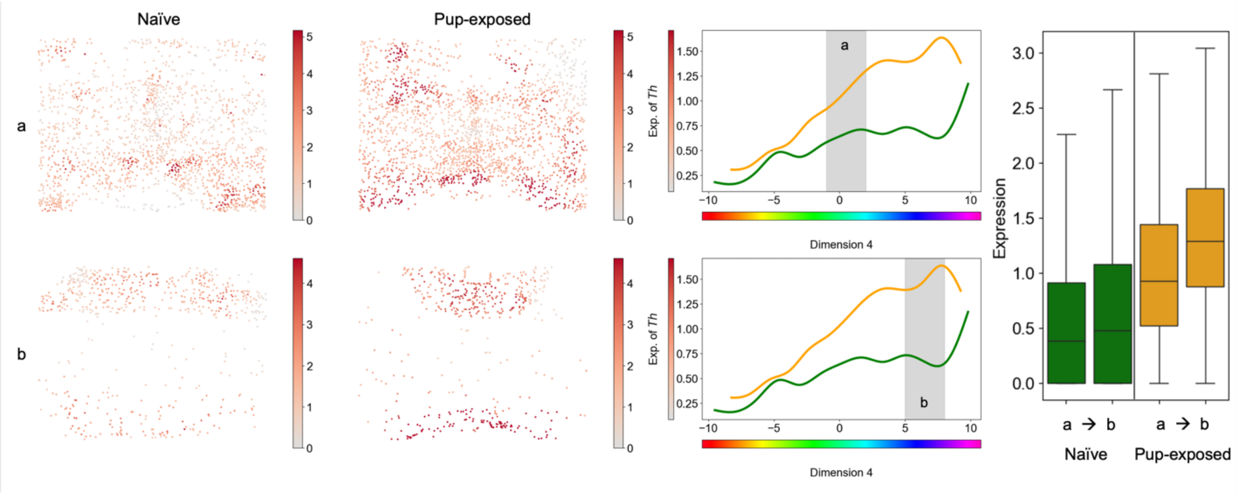


**Supplementary Figure S9**

Spatial expression visualization and gene expression versus embedding dimension plots for select genes. Expression levels of *Th* are shown in both naïve and pup-exposed samples (grey-red colormaps). For easier visualization, the colormaps in each row show cells in a subregion of dimension 4 (gray-shaded in the adjacent plot of expression versus dimension 4), where significant expression differences were detected. To emphasize the pattern differences, neighborhood smoothing was applied (n = 20), assigning each cell the average expression of its 20 nearest neighbors. (No such smoothing is used in the results shown in Figure 4 or 5.) The spline model fits illustrate gene expression trends along the embedding dimension where significant DSE was observed. The box plot on the right shows that expression of *Th* gene is similar between the subregions “a” and “b” for the naïve mouse brain but different between these two subregions for the pup-exposed mouse brain.

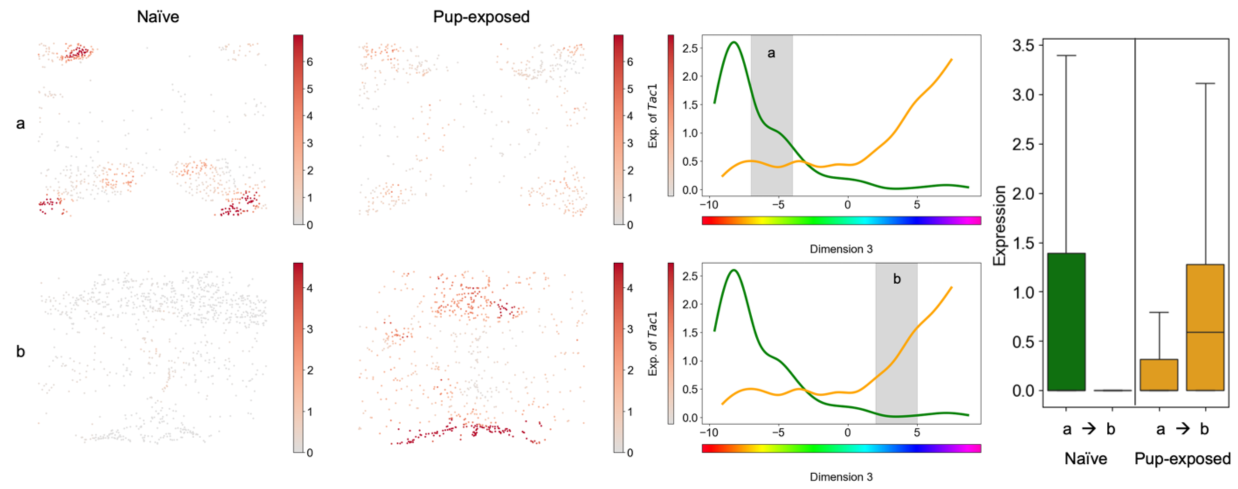


**Supplementary Figure S10**

The description is identical to Supplementary Figure S9, except this figure focuses on the *Tac1* gene, found to be DSE along dimension 3.

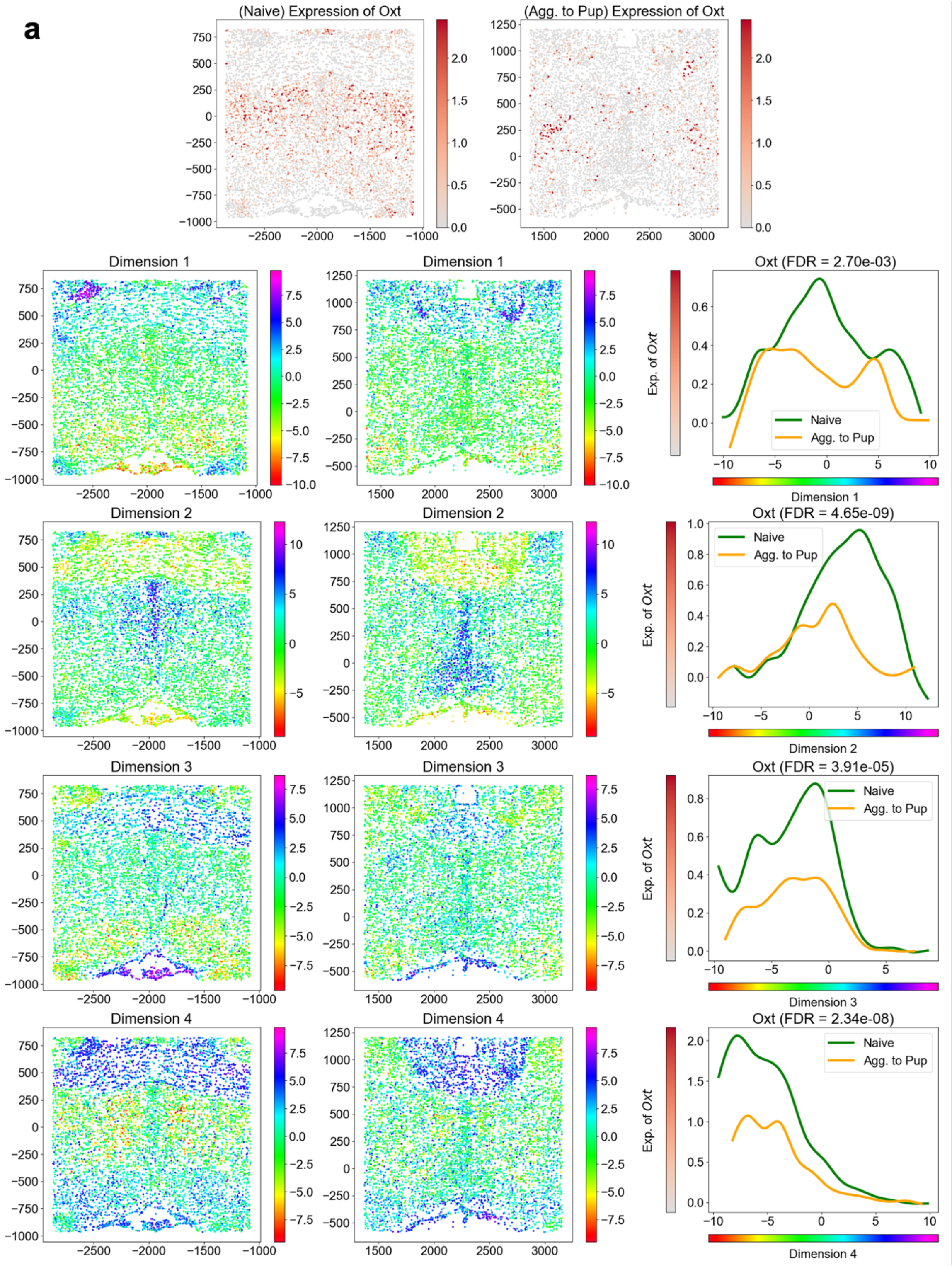

a
(Naive) Expression of Oxt
(Agg. to Pup) Expression of Oxt
Dimension 1
Dimension 2
Dimension 3
Dimension 4
Oxt (FDR = 2.70e-03)
Oxt (FDR = 4.65e-09)
Oxt (FDR = 3.91e-05)
Oxt (FDR = 2.34e-08)
Exp. of Oxt
Naive
Agg. to Pup

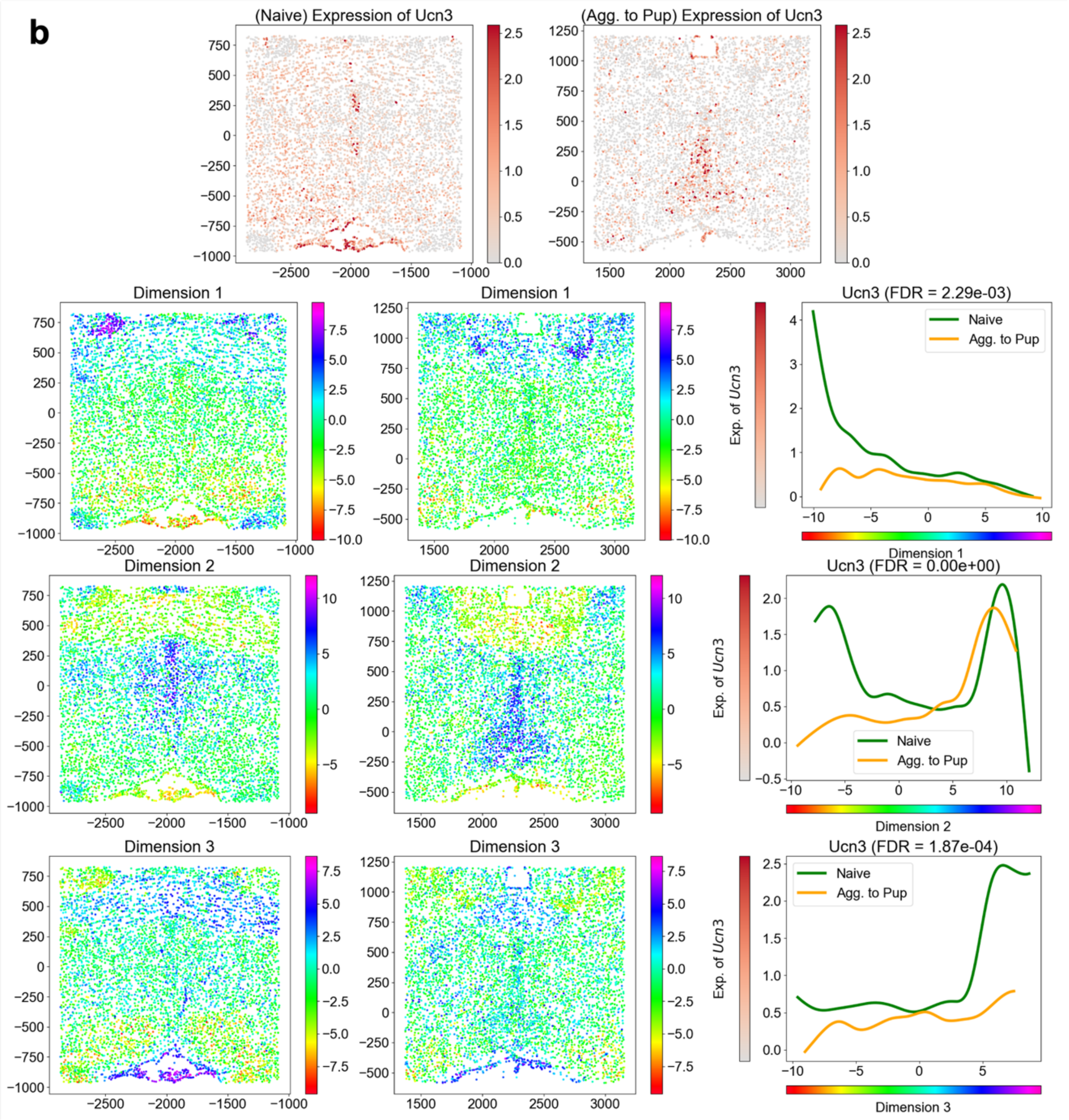

b
(Naive) Expression of Ucn3
(Agg. to Pup) Expression of Ucn3
Dimension 1
Dimension 2
Dimension 3
Ucn3 (FDR = 2.29e-03)
Ucn3 (FDR = 0.00e+00)
Ucn3 (FDR = 1.87e-04)
Naive
Agg. to Pup
Exp. of Ucn3

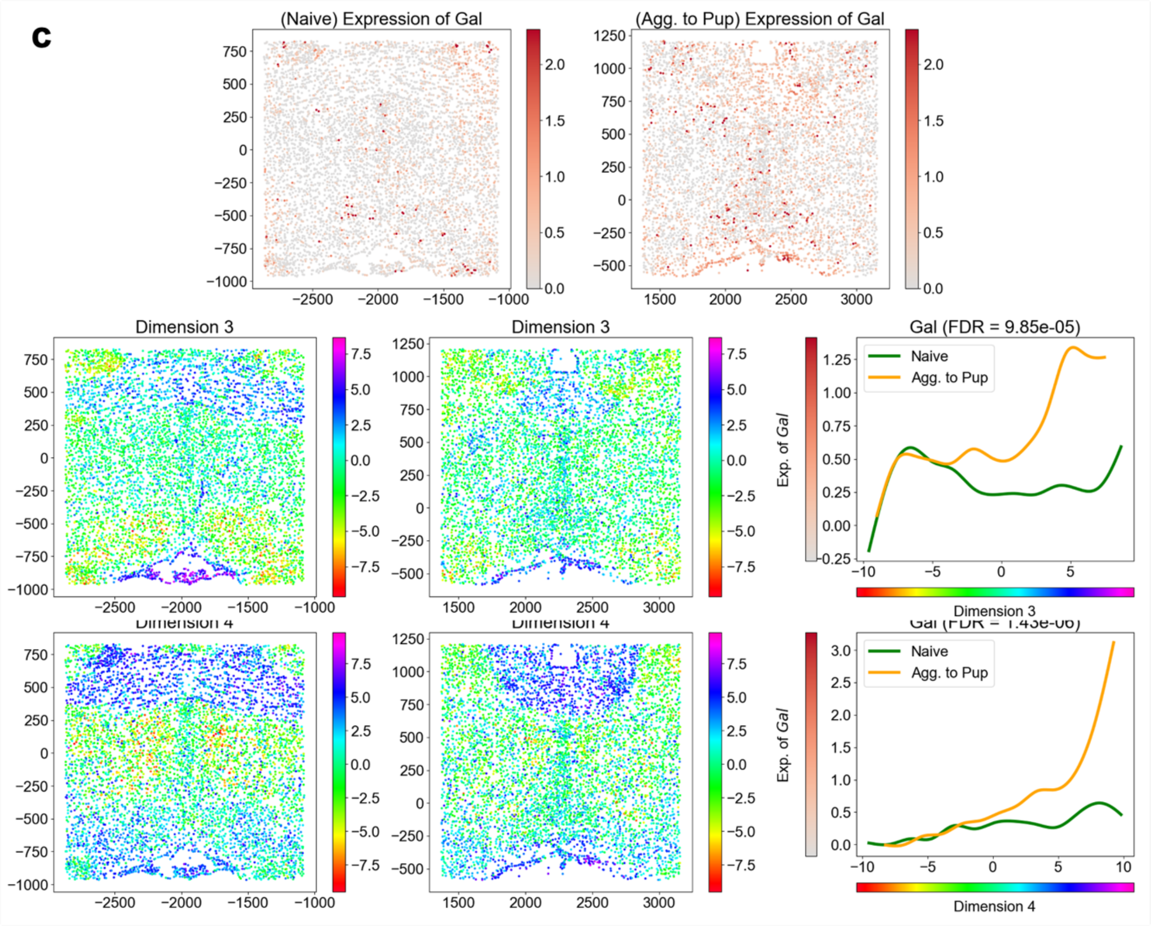


**Supplementary Figure S11**

Examples of DSE genes detected by SpaceExpress by comparison of whole brain MERFISH data from naïve and pup-exposed mice (sample IDs 5 and 36 respectively). Top panels display spatial expression patterns of (a) Oxt, (b) Ucn3, and (c) Gal in naïve (left) and pup-exposed (right) male mice. Color intensity represents the log-normalized expression of each gene across the tissue sections. The subsequent rows present visualizations of embedding dimensions along with spline model fits for gene expression, shown for the dimensions where significant DSE was detected (the same gene may be found DSE along more than one dimension). The full-spectrum color gradient reflects the embedding values of cells, capturing the spatial structure of the tissue. The corresponding spline plots (rightmost panel in each row) illustrate how gene expression varies along the embedding dimensions in naïve (green) and pup-exposed (orange) mice.

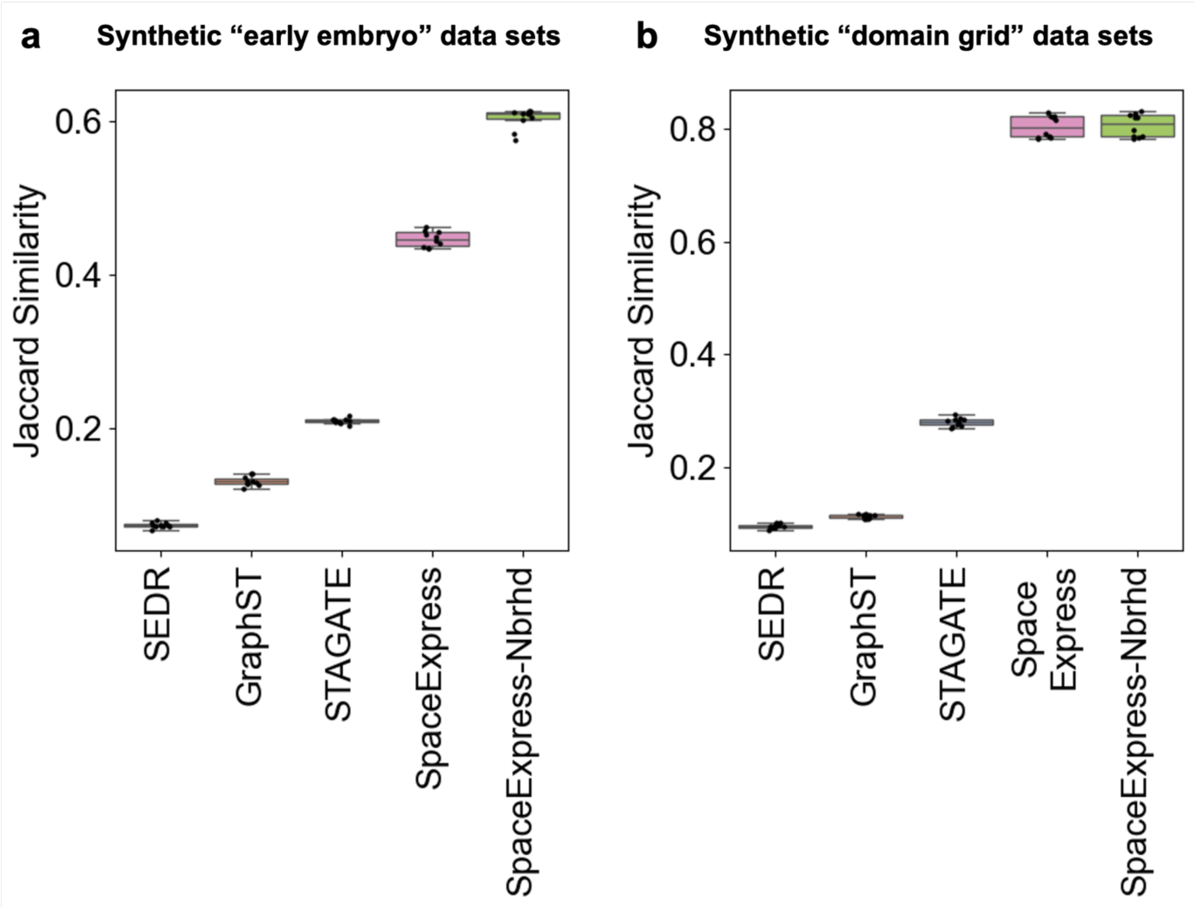


**Supplementary Figure S12**

Box plots of JS scores on synthetic early embryo data sets (a) and synthetic domain grid data sets (b) (n=10 replicates), comparing the performance of different embedding methods. We trained a variant model, called “SpaceExpress-Nbrhd”, that represents each cell by its own expression profile as well as the average expression profile of its neighboring cells, and maps the combined expression profile to the cell’s location using a neural network. We found the SpaceExpress-Nbrhd model to capture space even more accurately than SpaceExpress, though its embeddings are no longer determined solely by a cell’s own transcriptome. While superiority in capturing spatial locations is an encouraging sign, this alone does not mean that the learnt embedding dimensions are biologically meaningful. A model that learns embeddings from the expression and location information about cells may, in theory, “memorize” the expression-location mapping of those cells and thus excel in our evaluations above, yet not have captured the fundamental aspects of spatial organization of the tissue.

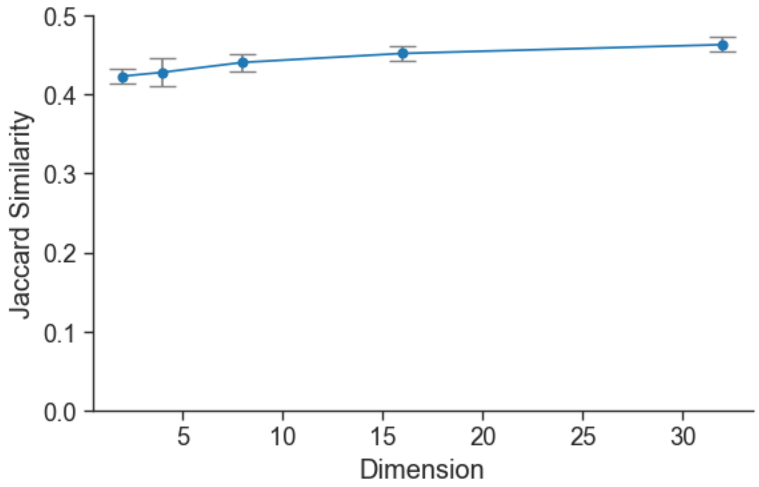


**Supplementary Figure S13**

Jaccard Similarity (k=10), as defined in Supplementary Note 1, between true spatial coordinates and the SpaceExpress embeddings is plotted for embedding dimensions 2, 4, 8, 16 and 32. Data points denote the mean Jaccard Similarity over five independent replicates; error bars indicate standard deviation. The consistency of Jaccard Similarity across all tested dimensions indicates that SpaceExpress reliably preserves spatial relationships regardless of the embedding dimensionality.

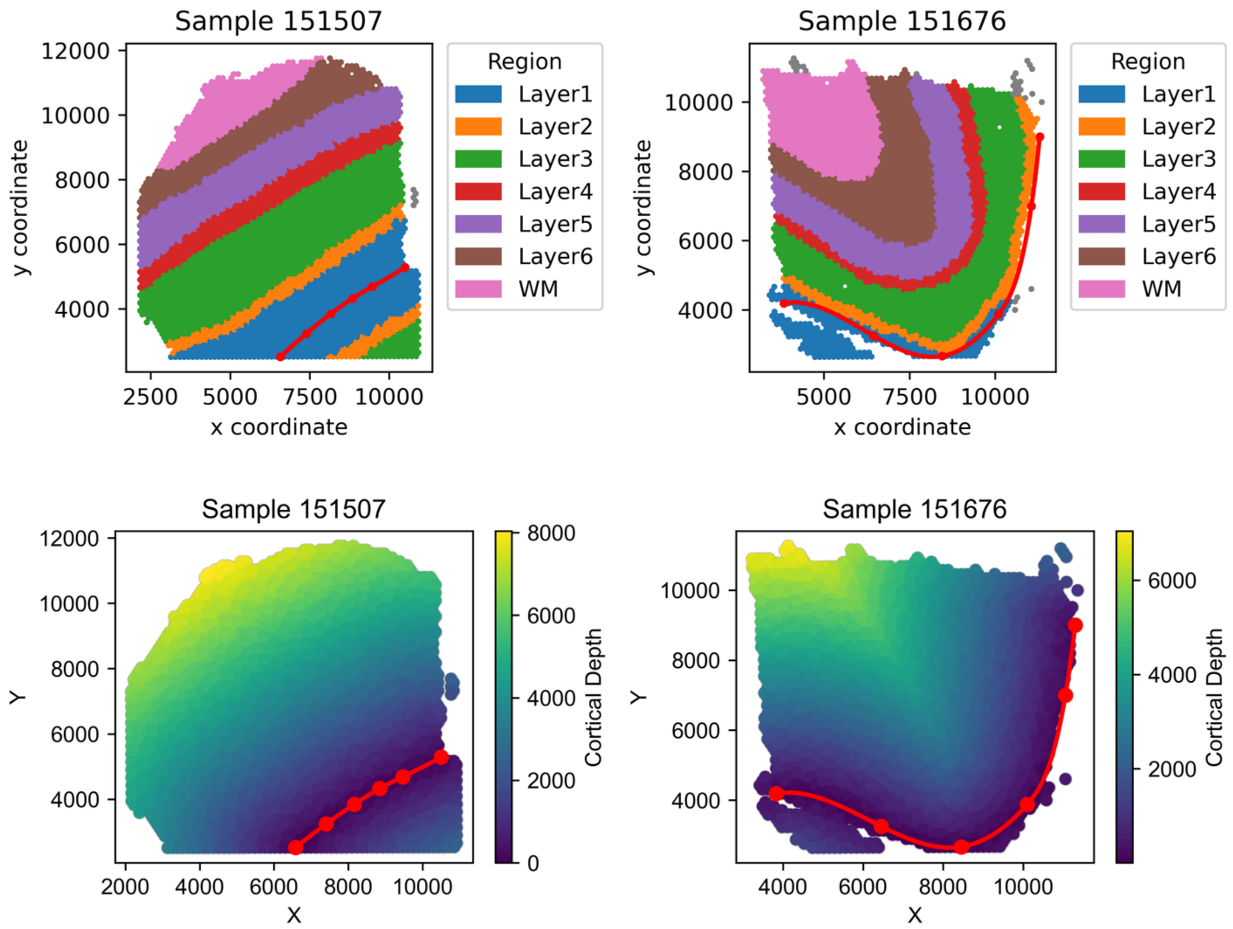


**Supplementary Figure S14**

Histologically defined cortical depth of DLPFC samples (151507, 151676). For each spatial transcriptomic sample, we manually annotated a series of anatomical landmarks along the cortical boundary (red circles). We then sorted these points by their x-coordinates and fit a cubic spline (red line) through them to approximate the cortical surface. Every cell's two-dimensional spot coordinate was compared to 100 equally spaced points along the spline, and we computed the Euclidean distances between the cell and each of these points. The minimum distance among these was assigned as the cell's cortical depth, shown in the bottom panels. The top panel shows the "layer" annotation of each sample, from Maynard et al.[2].

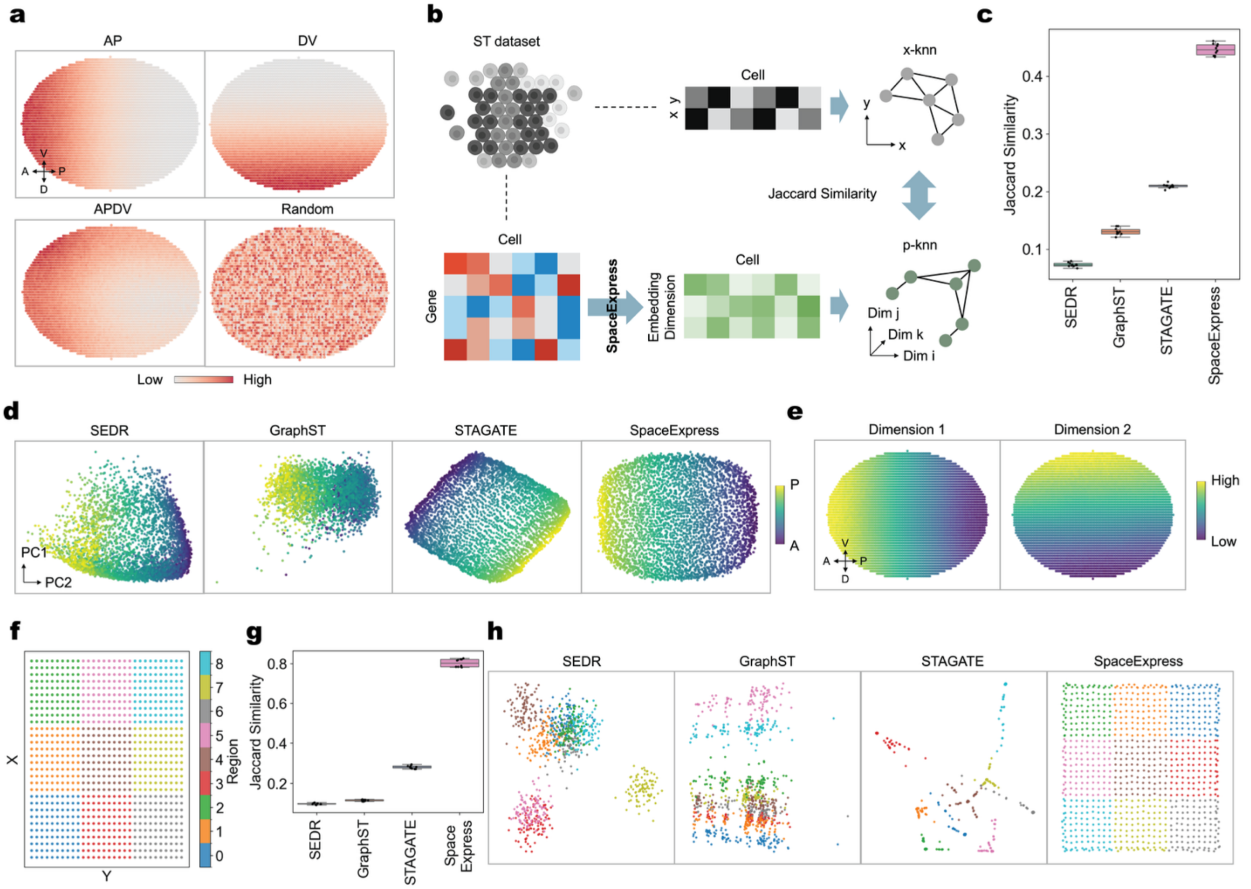


### Supplementary Figure S15

**(a)** Synthetic "early embryo" data set comprises cells arranged in an elliptical idealized embryo, with expression data on 100 genes whose spatial expression distribution has an anterior-posterior (AP) pattern, a dorsal-ventral (DV) pattern, a combination of AP and DV patterns (APDV) or a random distribution. In total, 10 replicates with different random noise were generated. **(b)** Schematic of Jaccard Similarity (JS) measure for evaluating SpaceExpress cell embeddings. Two neighborhood networks are constructed: the x-knn network, based on physical cell locations (x, y), and the p-knn network, based on the cell embeddings (panel b). JS (size of intersection divided by size of union) of the edges in these two networks is then used to quantify the similarity between these networks, providing a measure of how well the embedding preserves spatial relationships. JS has maximum value (1) when the two graphs are identical and its minimum value (0) if the two graphs share no edges. **(c)** Box plots of JS scores on synthetic early embryo data sets (n=10 replicates), comparing the performance of different embedding methods—SEDR, GraphST, STAGATE, and SpaceExpress. Higher JS scores indicate a better preservation of spatial structure in the embedding. Box plots depict interquartile range with median marked, with whiskers extending to 1.5 times the interquartile range; outliers are not displayed. **(d)** PCA visualizations of cell embeddings generated by SEDR, GraphST, STAGATE, and SpaceExpress for a synthetic early embryo data set. Each point represents a cell, colored by its position along the anterior-posterior (AP) axis. **(e)** Visualization of two of the embedding dimensions learnt by SpaceExpress from the synthetic early embryo data. Each point represents a cell at its physical location, colored by its coordinate value in the respective embedding dimension. **(f)** Illustration of the spatial regions within the synthetic “domain grid” data set. Nine square regions, each comprising 10 x 10 cells, are arranged in a 3 x 3 grid representing the tissue sample. Each region (shown in a different color) is marked by exclusive expression of one gene, while 91 additional genes are assigned random expression values. **(g)** Box plot of JS scores on synthetic “domain grid” data sets (n=10 replicates) for the four methods (SEDR, GraphST, STAGATE, and SpaceExpress). Box plots depict interquartile range with median marked, with whiskers extending to 1.5 times the interquartile range; outliers are not displayed. **(h)** PCA visualizations of spatial embeddings across methods, with points colored by their spatial regions (panel h) in the synthetic tissue.

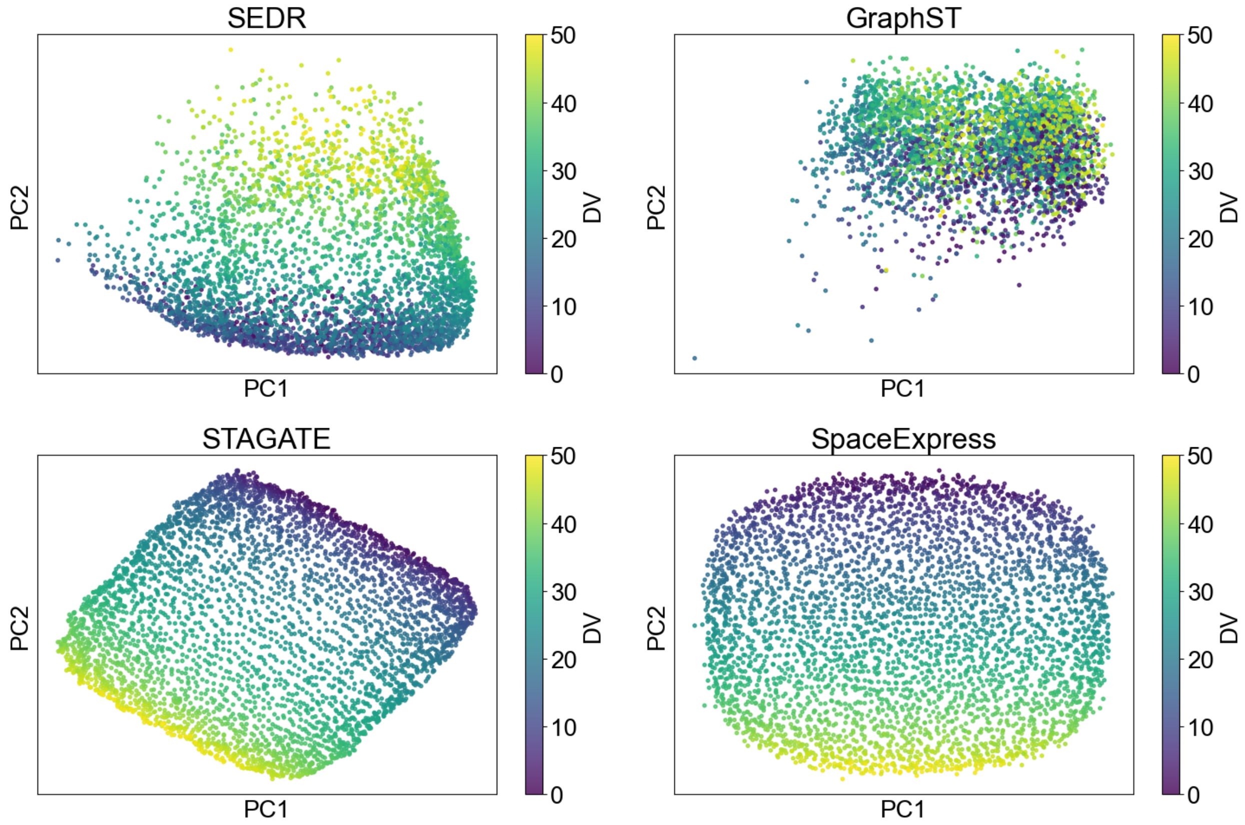


**Supplementary Figure S16**

PCA visualizations of cell embeddings generated by SEDR, GraphST, STAGATE, and SpaceExpress for the synthetic early embryo data. Each point represents a cell, colored according to its position along the dorsal-ventral (DV) axis. SpaceExpress embeddings accurately capture the DV locations of cells in the embryo.

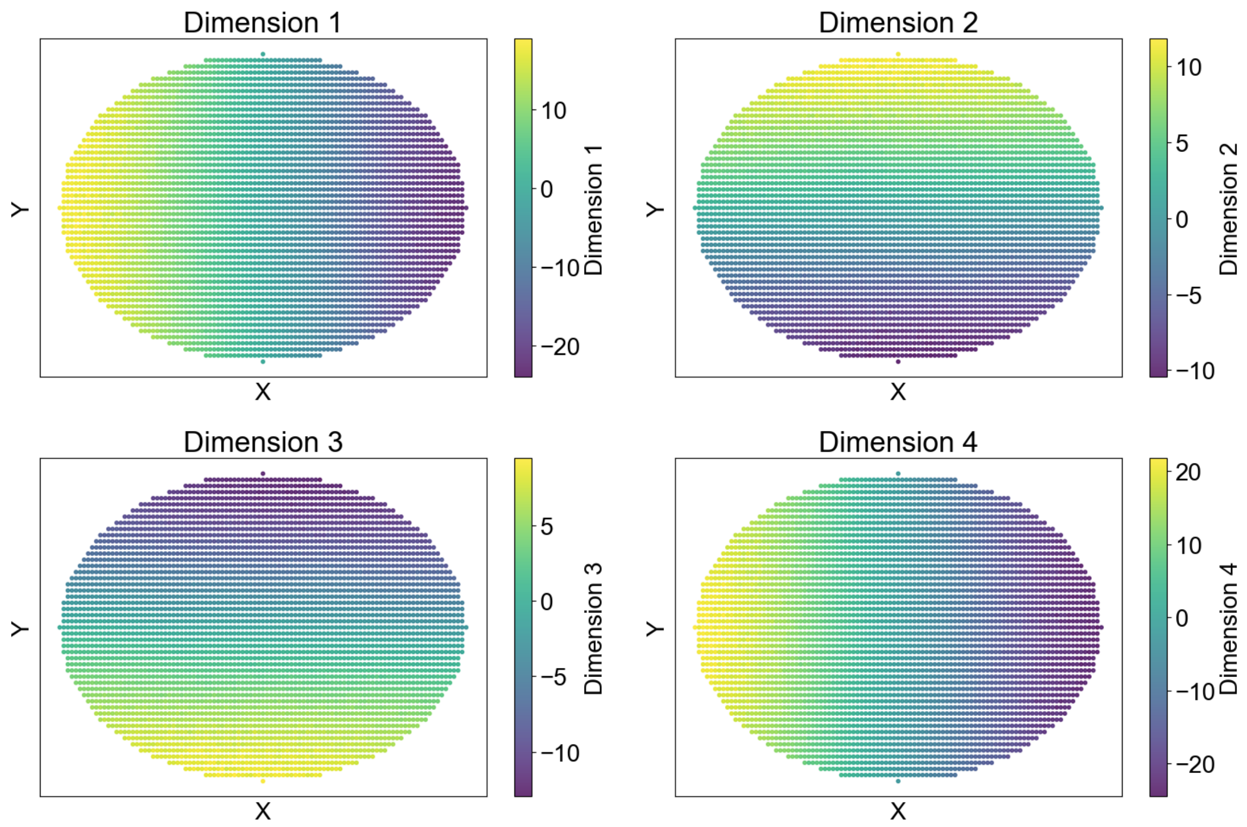


**Supplementary Figure S17**

Visualization of the SpaceExpress embedding dimensions derived from the synthetic early embryo data. Each point corresponds to a cell, positioned according to its physical coordinates, and is colored based on its embedding dimensions. Dimensions 1 and 4 capture the AP axis while dimensions 2 and 3 capture the DV axis; these two axes form an “intrinsic” coordinate system of the embryo, by construction.

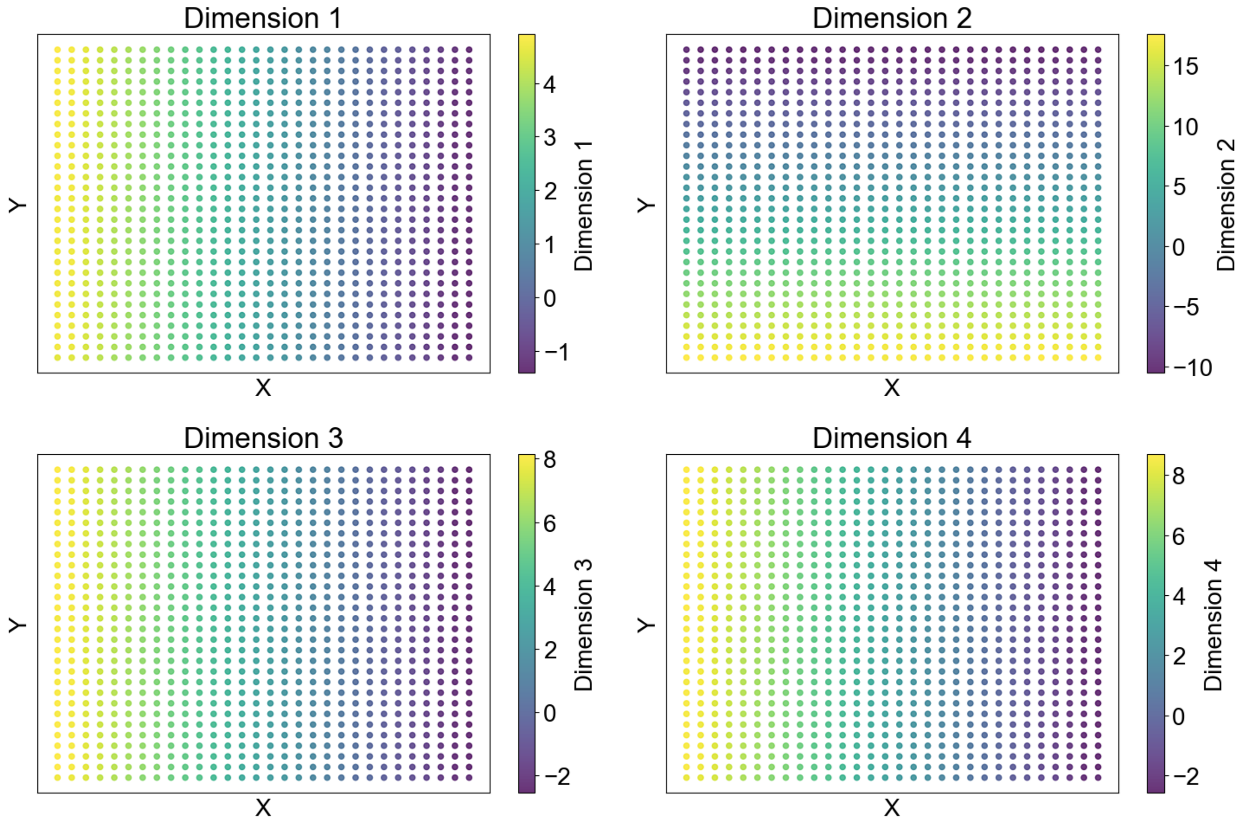


**Supplementary Figure S18**

Visualization of SpaceExpress embedding dimensions learned from the synthetic “domain grid” dataset. Each point represents a cell located at its physical position within the tissue, colored according to the values in each embedding dimension. The combination of these four dimensions captures the domain grid organization of cells in this synthetic tissue, as shown in Figure 1J.

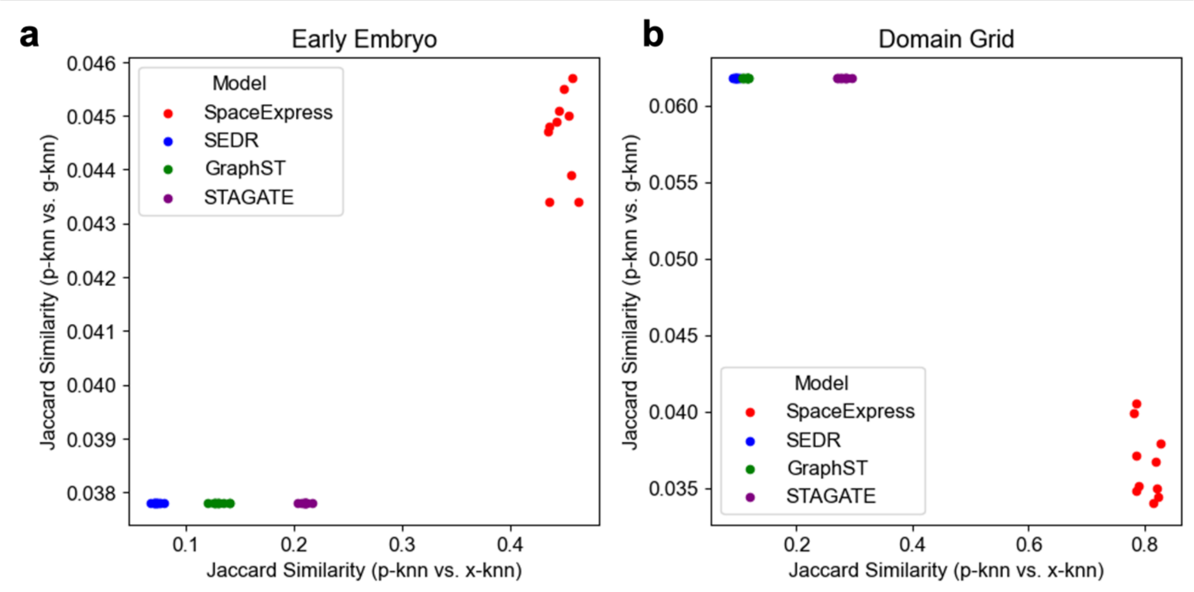


**Supplementary Figure S19**

Jaccard Similarity (JS) analysis comparing neighborhood networks in the synthetic early embryo (a) and domain grid (b) datasets. The x-axis is the JS score between a k-nearest neighbor network based on physical cell coordinates (“x-knn”) and that based on spatial embeddings (“p-knn”), assessing how well the embeddings preserve spatial relationships. The y-axis is the JS score between the p-knn and a nearest neighbor network based on expression profiles of cells (“g-knn”), evaluating the extent to which embeddings reflect cellular transcriptomes. We observe that SpaceExpress embeddings capture space substantially better than other methods, in both types of data sets (a,b). They reflect transcriptomes more strongly than other methods for the early embryo data set but less strongly than other methods for the domain grid data set. These analyses underscore the different tradeoffs an embedding of ST data makes in capturing spatial and expression information about cells.

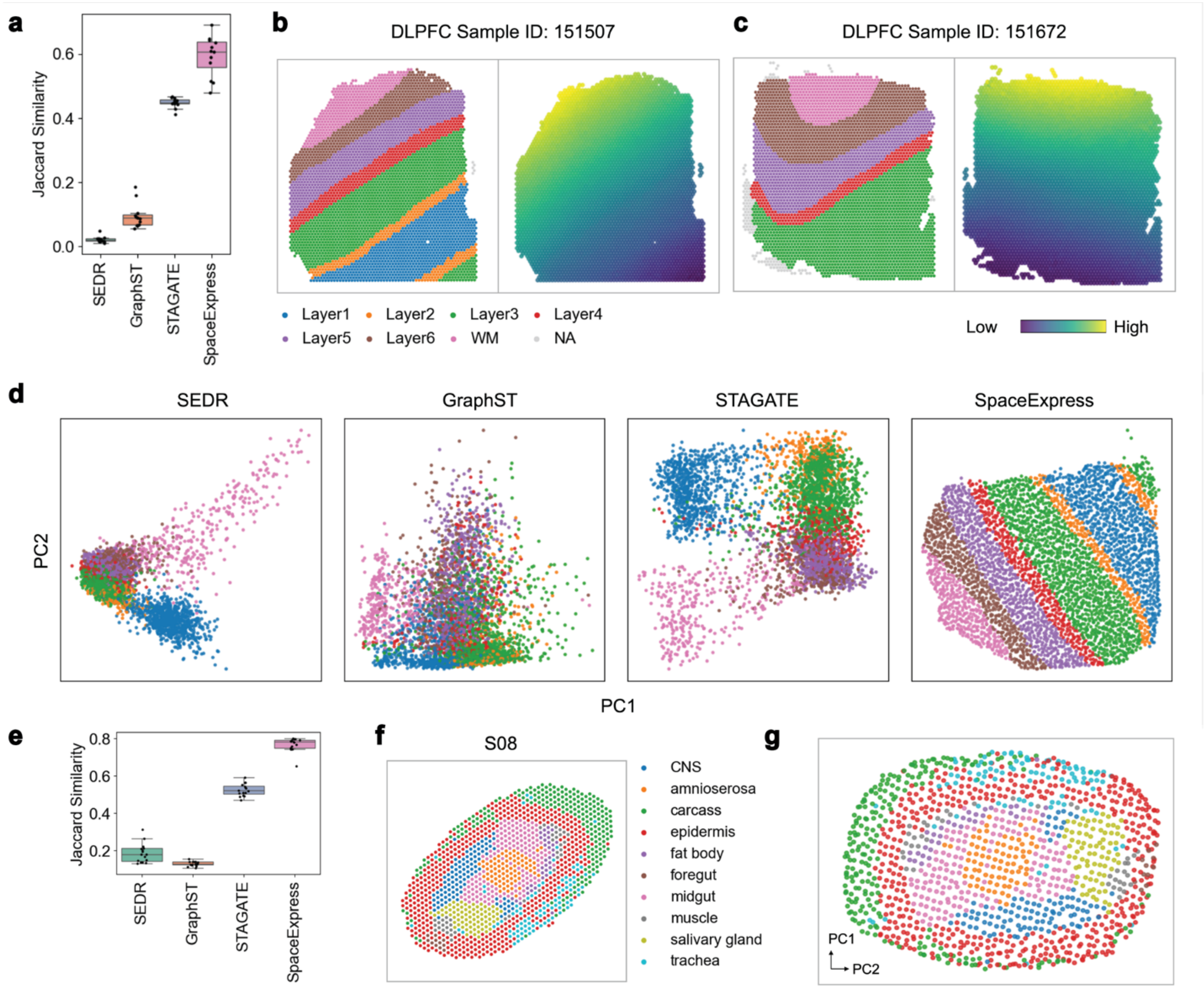


**Supplementary Figure S20**

**(a)** Box plots showing Jaccard Similarity (JS) scores on 12 tissue samples from the DLPFC dataset when embedding cells in a tissue using SEDR, GraphST, STAGATE, or SpaceExpress. Box plots depict interquartile range with median marked, with whiskers extending to 1.5 times the interquartile range; outliers are not displayed. **(b, c)** Visualization of tissue samples ID 151507 (b) and ID 151672 (c), showing spatial tissue organization into cortex layers and white matter (left panel) and a colormap visualization of one of the embedding dimensions learnt by SpaceExpress (right panel). (**d**) PCA plots of cell embeddings learnt by different methods on tissue sample ID 151507. Points are colored according to their cortex layer assignments (panel b). **(e)** Box plot of JS scores on 16 slices of Drosophila embryo (E14-16) data, comparing the performance of the four embedding methods. Box plots depict interquartile range with median marked, with whiskers extending to 1.5 times the interquartile range; outliers are not displayed. **(f)** Visualization of Drosophila embryo slice S08, with colors indicating annotated spatial regions/tissues. **(g)** PCA plots of SpaceExpress cell embeddings of Drosophila embryo slice S08, showing that the learnt embeddings capture known tissue structure.

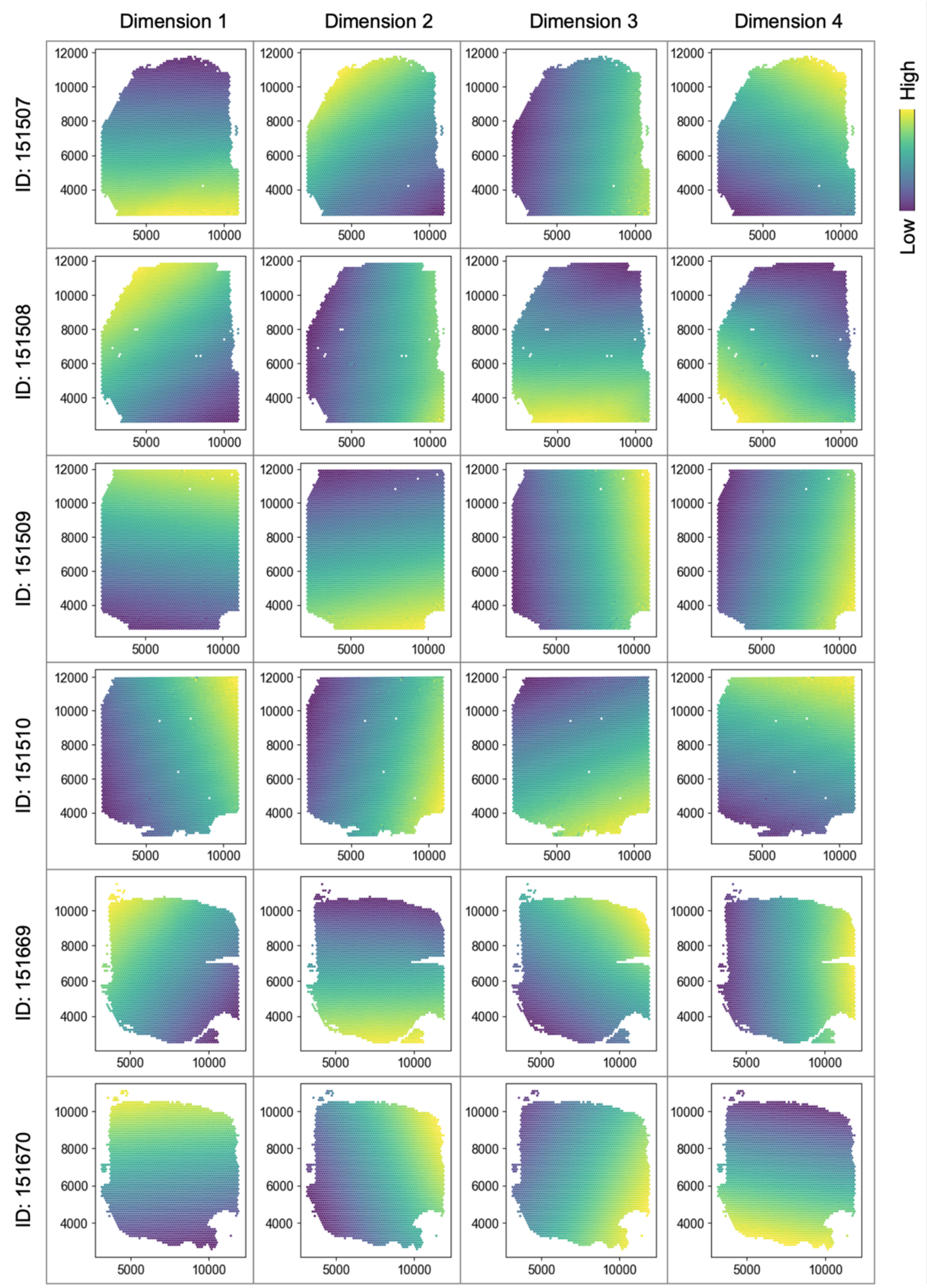

Dimension 1
Dimension 2
Dimension 3
Dimension 4
ID: 151507
ID: 151508
ID: 151509
ID: 151510
ID: 151669
ID: 151670
High
Low

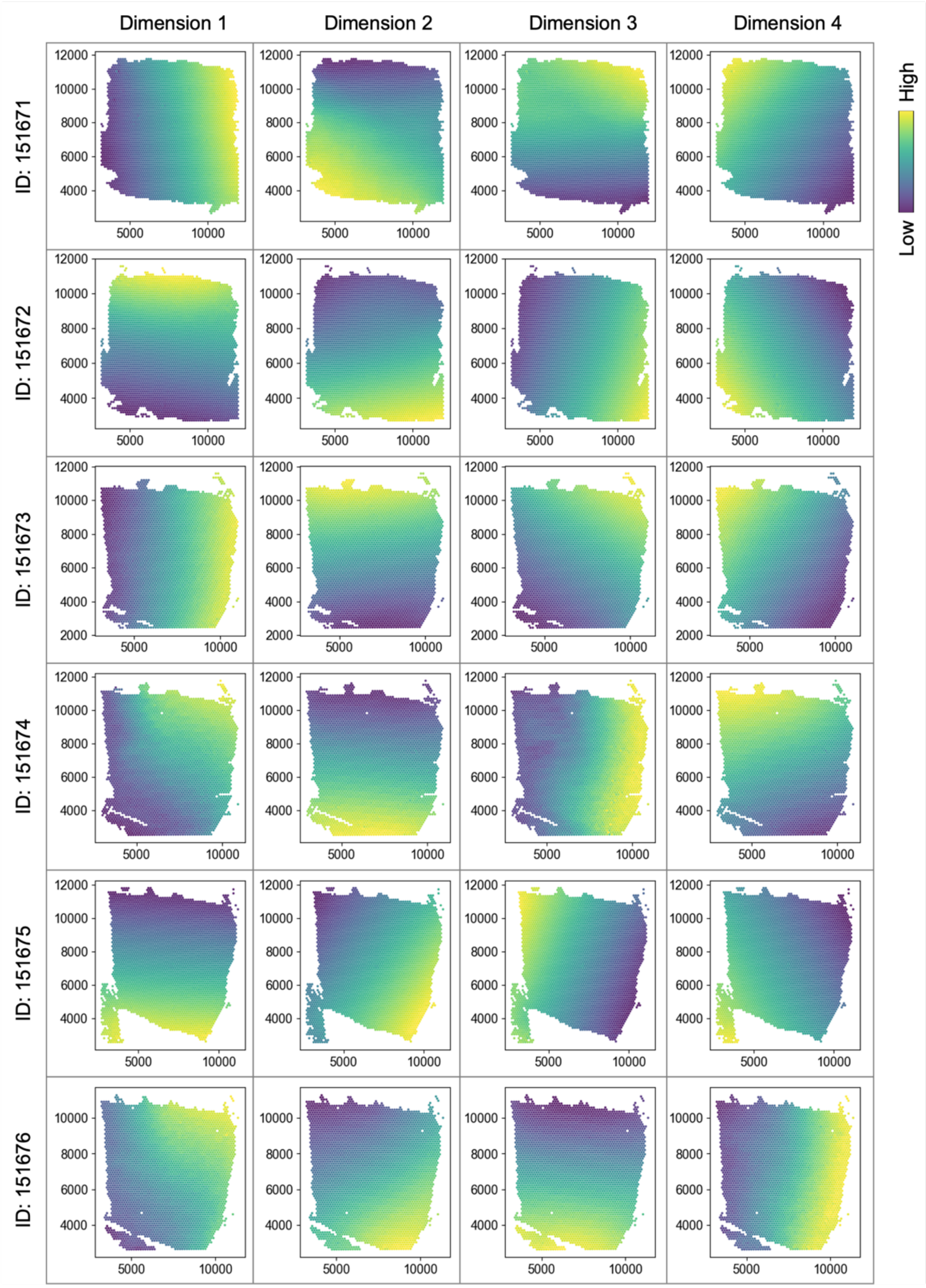


**Supplementary Figure S21**

Visualization of SpaceExpress embedding dimensions for each of 12 DLPFC tissue samples. Each point represents a cell at its physical position, colored by its values in the embedding dimension 1-4. These are results of training the SpaceExpress embedding function on each sample separately.

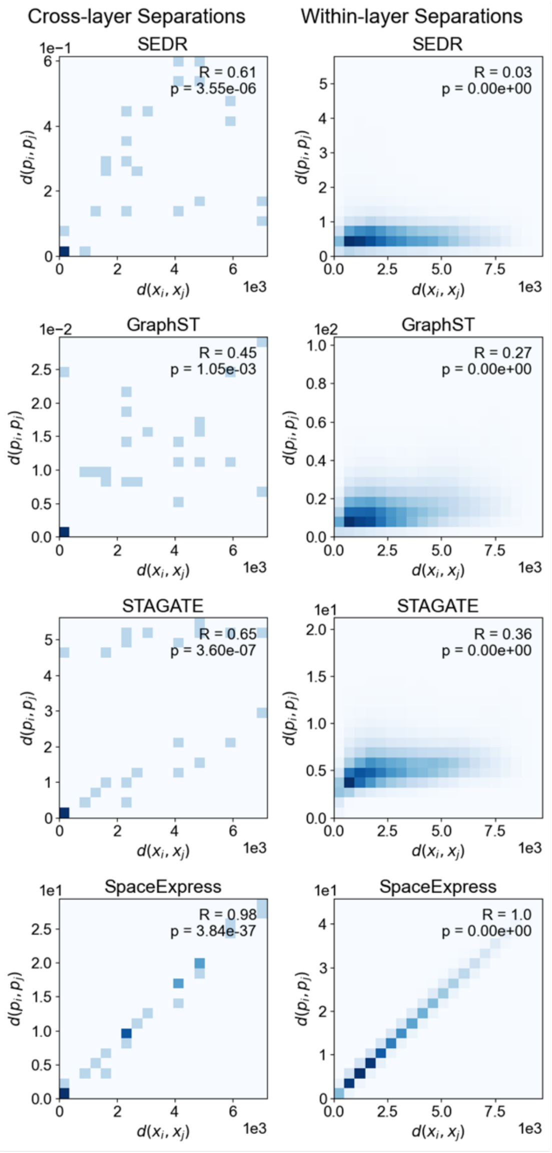


**Supplementary Figure S22**

Comparison of cell embedding distances (y-axis) generated by SEDR, GraphST, STAGATE, and SpaceExpress, trained on DLPFC sample ID 151507, to physical distances (x-axis) between cell pairs in the same sample. The left plot shows inter-layer distances, where each point represents the centroid distance between pairs of tissue layers. The right plot displays a 2D histogram of the joint distribution of embedding distances and physical distances between cell pairs within the same layer, aggregated across all layers.

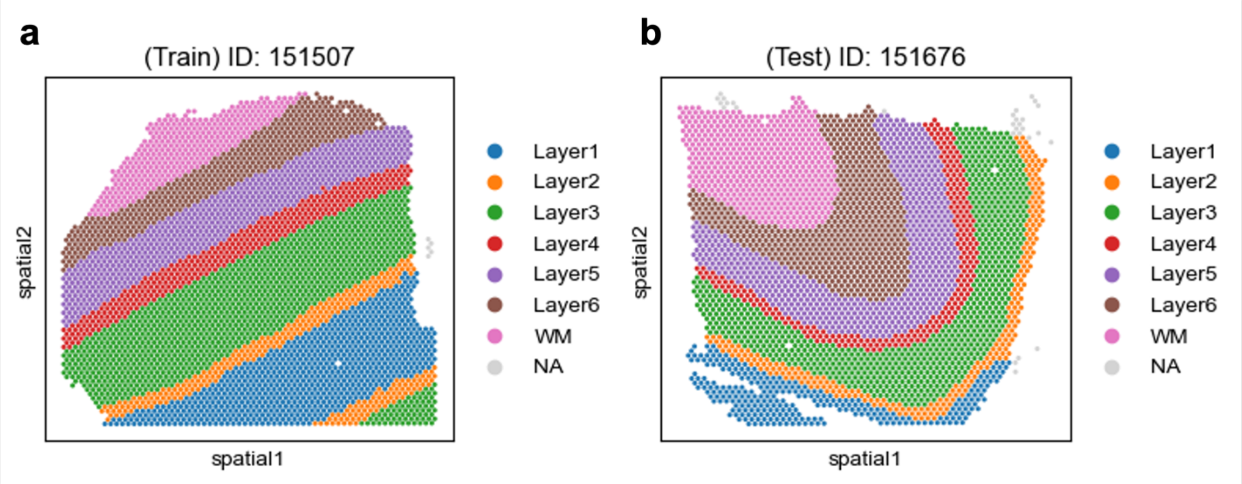


**Supplementary Figure S23**

Spatial organization of DLPFC tissue samples ID 151507 (a) and ID 151672 (b), showing cortex layers and white matter. The SpaceExpress embedding model trained on sample ID 151507 was applied to embed cells in a different sample, ID 151676, using only gene expression data (excluding spatial information). (Results shown in other figures.)

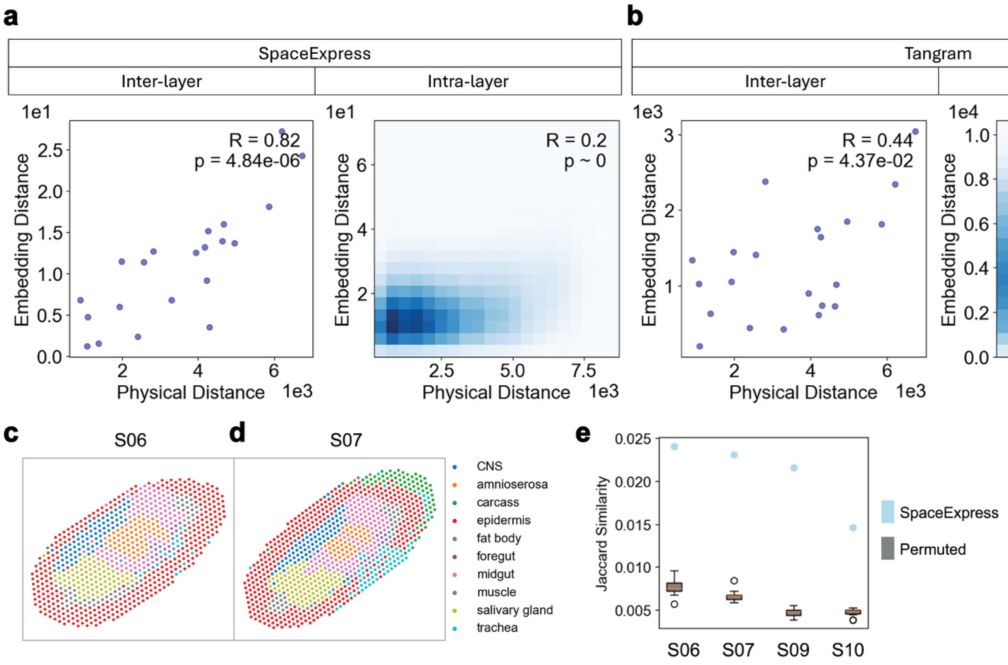


### Supplementary Figure S24

**(a)** SpaceExpress cell embedding function trained on one DLPFC sample (ID 151507, Supplementary Figure S23a) was used to embed (assign coordinates to) cells in a different sample (ID 151676, Supplementary Figure S23b), using only expression (without spatial) data from the latter. Distances between embeddings of cell pairs (y axis) were compared to their physical distance (Euclidean distance) (x axis), for cell pairs belonging to the different layers (left) or same layer (right) of the tissue. Each point in the inter-layer plot (left) represents one pair of layers, showing the (physical or embedding) distances between centroids of the two layers**.** The intra-layer plot (right) is a 2D histogram of the joint distribution of embedding and physical distances between cell pairs in the same layer, aggregated over all layers. **(b)** Evaluation of a cell mapping method, TANGRAM, using the same procedure as in (a). **(c, d)** Visualization of Drosophila embryo slices S06 (c) and S07 (d), where cells are colored based on their annotated spatial regions/tissues. **(e)** Box plot of Jaccard Similarity (JS) values of SpaceExpress embeddings for slices S06 and S07, obtained without using spatial information on these tissues. Also shown for comparison are corresponding JS scores of a random baseline ("Permuted") where SpaceExpress embeddings were trained on a version of slices S08 where the spatial locations of cells were randomly permuted. Box plots depict interquartile range with median marked, with whiskers extending to 1.5 times the interquartile range; outliers are not displayed.

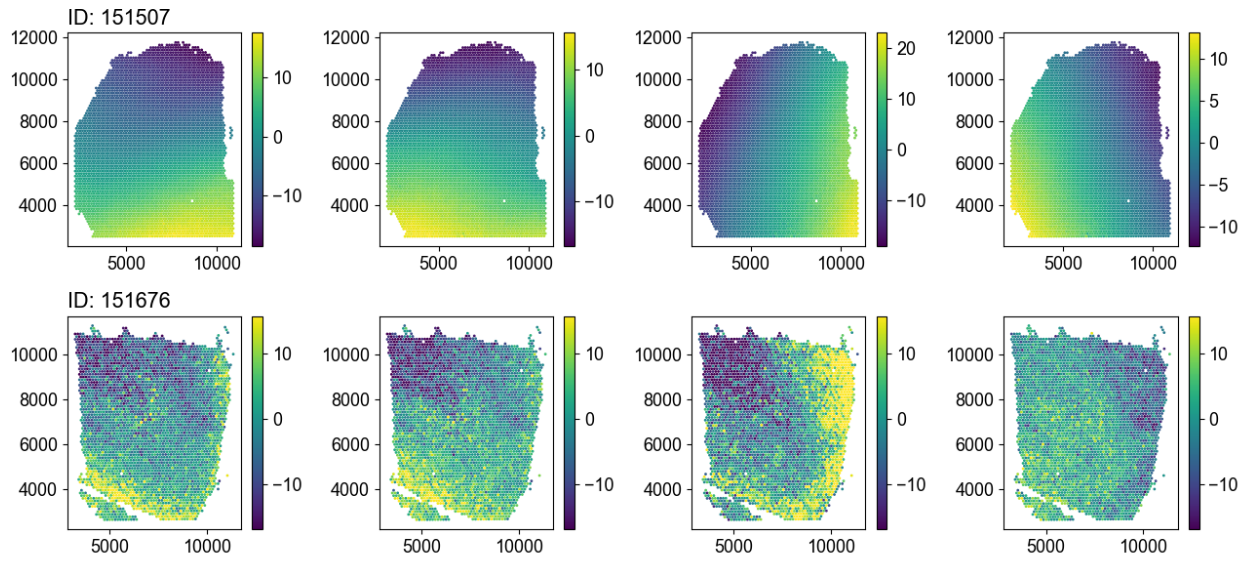


**Supplementary Figure S25**

SpaceExpress embedding dimensions trained on DLPFC sample ID 151507 (using expression and spatial data) and used to embed cells in sample ID 151676 using only expression data. Each point represents a cell at its physical location, colored by the respective values in the learned embedding dimensions. For instance, dimension 3, which is approximately parallel to the layers in the training sample (ID 151507), accurately captures the curved shape of those layers in the test sample (ID 151676), indicating robustness in the face of structural distortions.

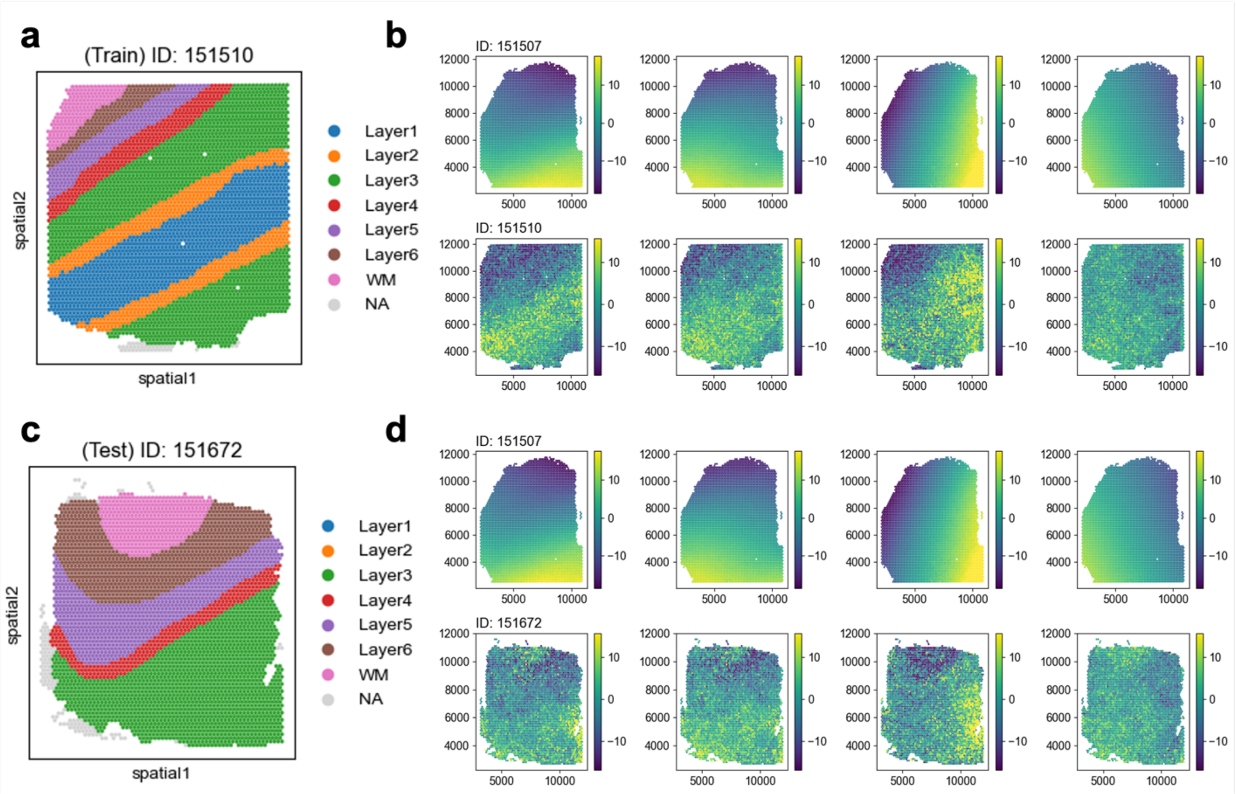


**Supplementary Figure S26**

(a, c) Visualization of tissue samples ID 151510 (a) and ID 151672 (c). (b, d) Visualization of SpaceExpress embedding model trained on sample ID 151507 (Figure S7a) and used to embed cells in sample ID 151676 (b) and ID 151672 (d) using only expression data on these “test” samples. Cells are shown at their physical locations, colored by embedding values. We see that analogous regions of the test and training samples are assigned similar coordinate values in each embedding dimension, supporting the generalizability of SpaceExpress embeddings.

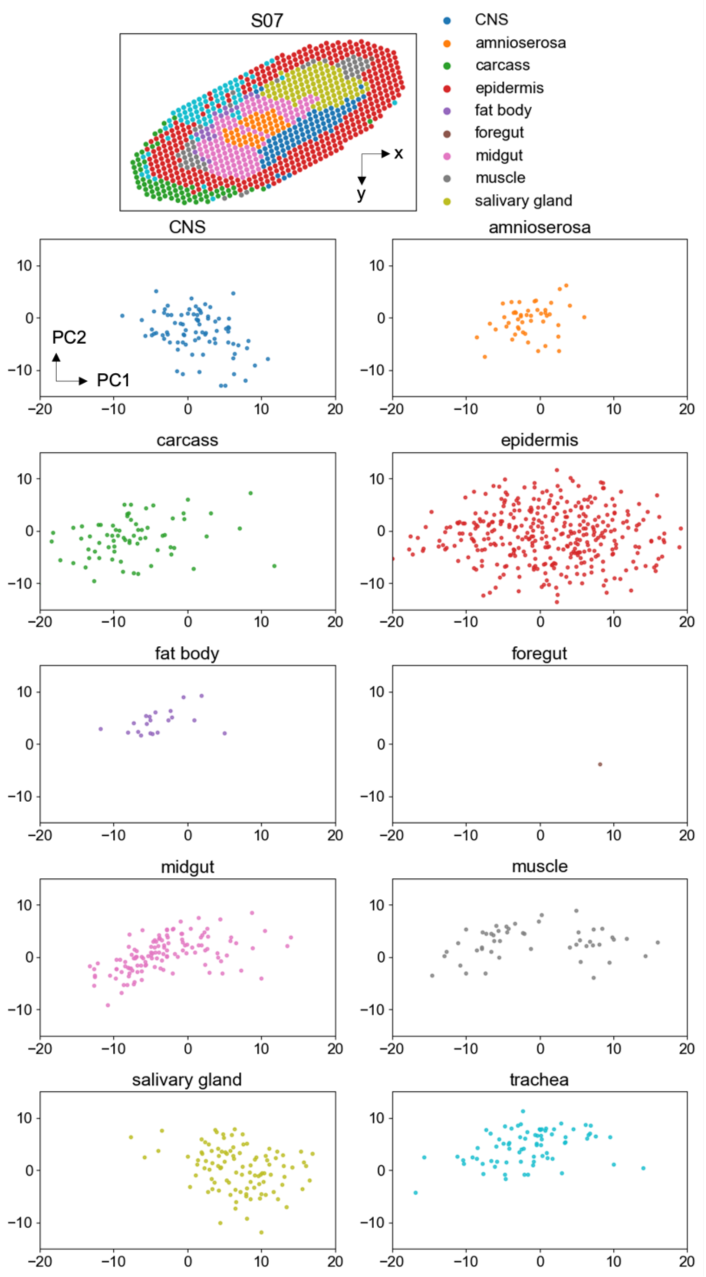


### Supplementary Figure S27

PCA plots of SpaceExpress cell embeddings for Drosophila embryo slice S07 (top), generated without utilizing spatial information. Each point represents a cell, colored based on its assigned spatial regions/tissues. All cells were projected to the PC1, PC2 space, and then plotted in separate panels grouped by their spatial region, for clarity.

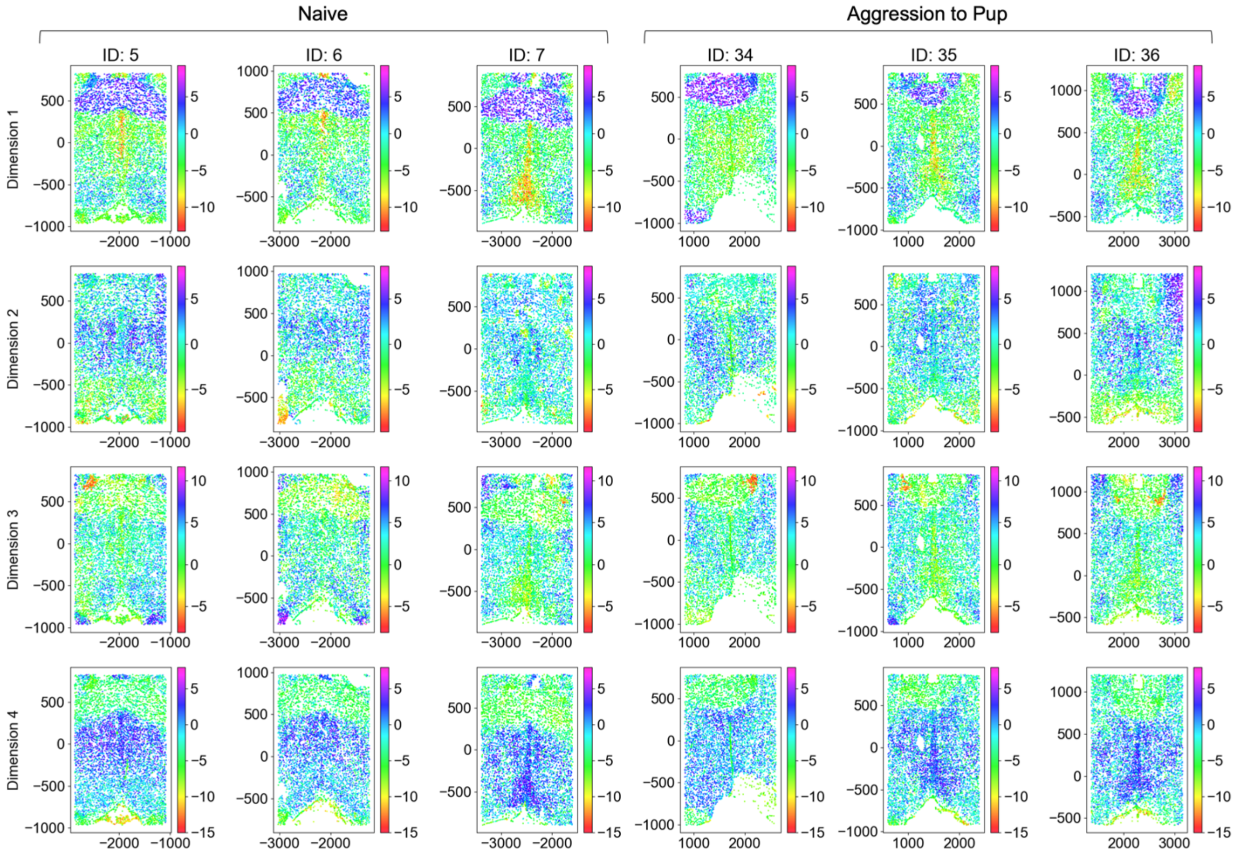


**Supplementary Figure S28**

Visualization of embedding dimensions learnt by SpaceExpress in the joint analysis of MERFISH data on six mouse brains (hypothalamic preoptic area). Each panel shows the embedding values learned by SpaceExpress for a different sample, with the left section depicting naïve mice (IDs 5, 6, and 7) and the right section showing pup-exposed aggressive mice (IDs 34, 35, and 36). The rows represent the four different embedding dimensions, and the color gradient from red to purple reflects the embedding values of cells.

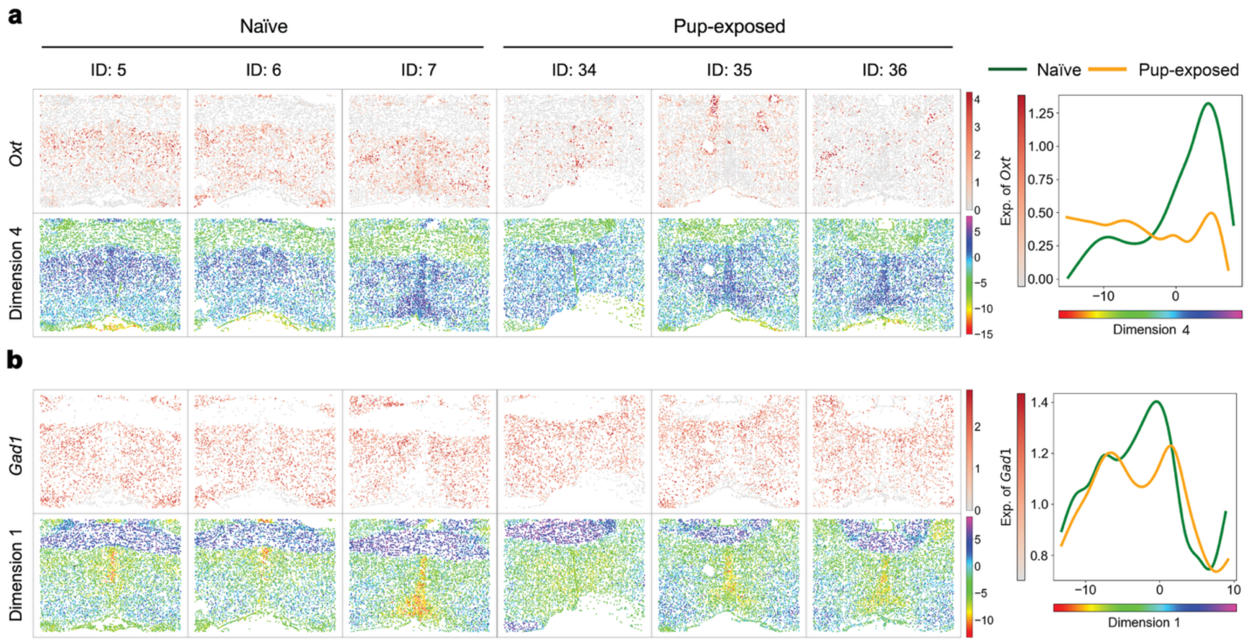


**Supplementary Figure S29**

Shown are spatial expression (grey-red colormaps) and embedding dimension (full spectrum colormaps) visualizations for *Oxt* (a) and *Gad1* (b) in whole brain of naïve and pup-exposed mice. The six columns represent three replicates from the naïve group and three from the pup-exposed group. The plots on the right show spline model fits of gene expression against coordinate values for the embedding dimension visualized.

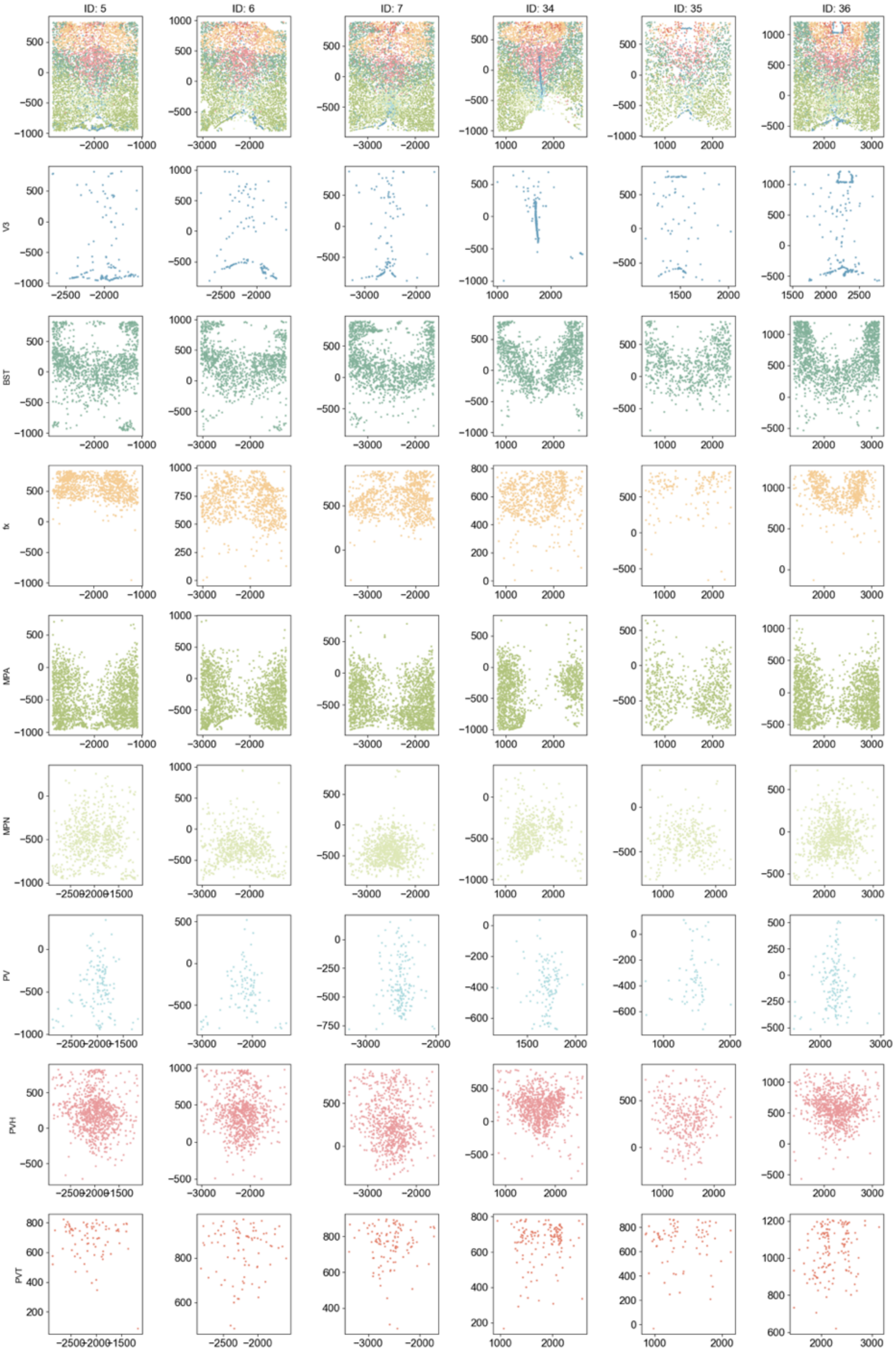
ID: 5
ID: 6
ID: 7
ID: 34
ID: 35
ID: 36
V3
BST
fx
MPA
MPN
PV
PVH
PVT

**Supplementary Figure S30 (previous page)**

Annotations of regions in MERFISH data of the hypothalamic preoptic area of six mouse brains. Each column corresponds to a different sample, with naïve mice (IDs 5, 6, 7) on the left and pup-exposed mice (IDs 34, 35, 36) on the right. The top row shows the neuroanatomically annotated regions transferred using SLAT from the manually annotated samples with colors representing distinct anatomical domains that capture spatial structures within the tissue. The subsequent rows display the spatial expression patterns for each domain across the tissue.

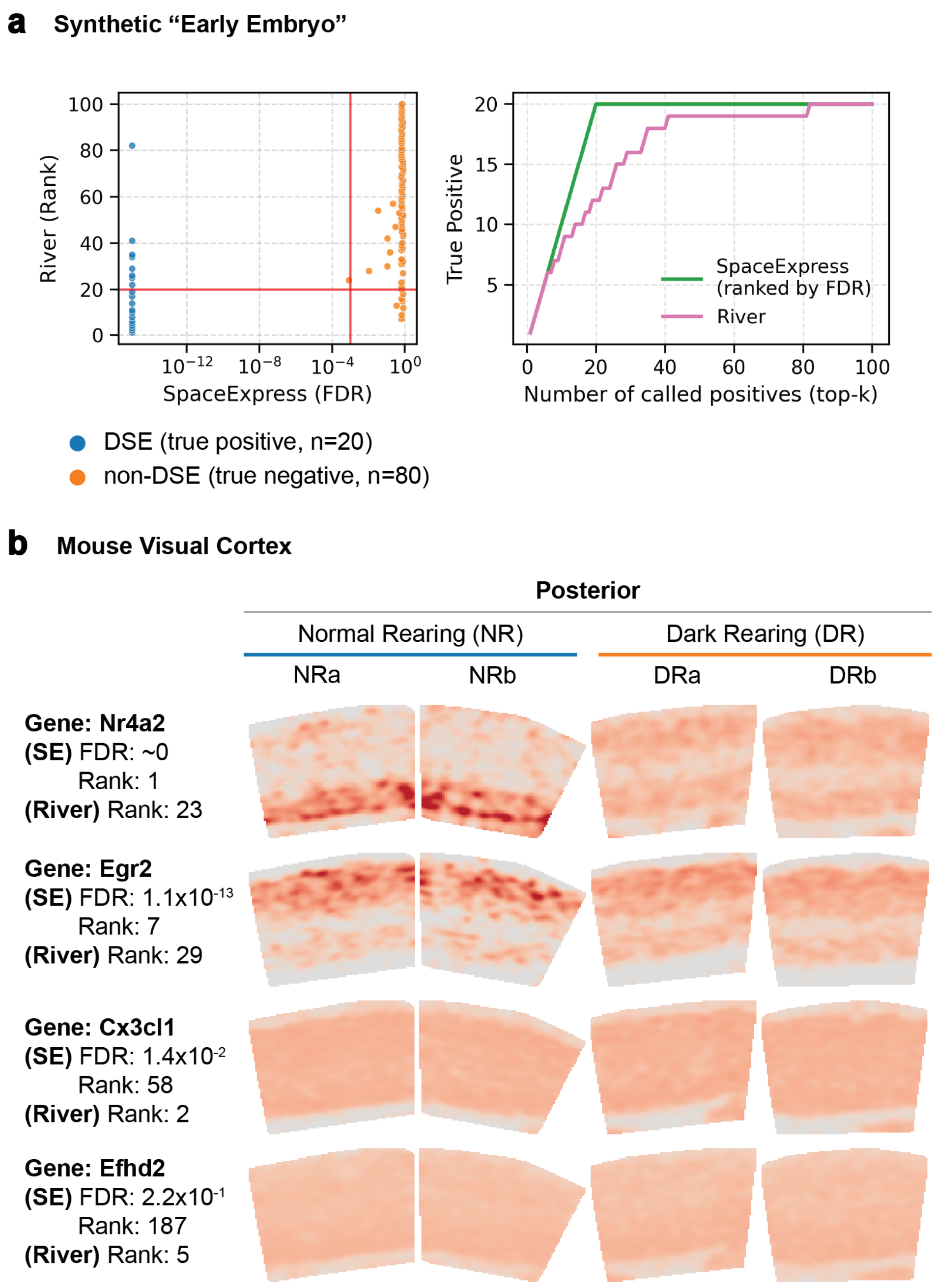


### Supplementary Figure S31

Comparison of DSE detection by SpaceExpress and River on real and synthetic data sets. **a.** Synthetic "early embryo" dataset with two slices simulated using $\mu$=0 and $\mu$=0.6 respectively, thus representing a modest level of expression divergence for 20 of the 100 genes simulated (see Fig. 2a and 3a). We ran both methods with default settings. (Right) The scatter plot show SpaceExpress-assigned FDR versus River-assigned rank for 100 genes, of which 20 were designed to be DSE across the two synthetic slices. Red lines mark the SpaceExpress FDR threshold ($1\times10^{-3}$) and the top-20 cutoff for River. SpaceExpress recovered all 20 true DSE at FDR $\leq1\times10^{-3}$ with one non-DSE among its discoveries, whereas River placed eight non-DSE genes within its top 20 ranks. (Left) The same data is shown in terms of true positives detected as a function of rank, for either method. **b.** Mouse visual cortex (posterior) dataset (see Fig. 5a,b). Each row shows one gene across two normal-rearing (NRa, NRb) and two dark-rearing (DRa, DRb) replicates; labels on the left indicate the SpaceExpress FDR and River rank. The top two rows are genes highly ranked by River but not considered significant by SpaceExpress at FDR $\leq 1\times10^{-3}$. The bottom two rows are genes with very significant SpaceExpress FDR that were not highly ranked by River. Since “ground truth” is not available for this real data set, we have shown examples where visual inspection strongly suggests a DSE designation, along with the DSE scores/ranks from either method to assess consistency with that designation.
